# Fragment-based discovery of a new class of inhibitors targeting mycobacterial tRNA modification

**DOI:** 10.1101/564013

**Authors:** Sherine E. Thomas, Andrew J. Whitehouse, Karen Brown, Juan M. Belardinelli, Ramanuj Lahiri, M. Daben J. Libardo, Pooja Gupta, Sony Malhotra, Helena I. M. Boshoff, Mary Jackson, Chris Abell, Anthony G. Coyne, Tom L. Blundell, R. Andres Floto, Vítor Mendes

## Abstract

Translational frameshift errors are often deleterious to the synthesis of functional proteins as they lead to the production of truncated or inactive proteins. TrmD (tRNA-(N(1)G37) methyltransferase) is an essential tRNA modification enzyme in bacteria that prevents +1 errors in the reading frame during protein translation and has been identified as a therapeutic target for several bacterial infections. Here we validate TrmD as a target in *Mycobacterium abscessus* and describe the application of a structure-guided fragment-based drug discovery approach for the design of a new class of inhibitors against this enzyme. A fragment library screening followed by structure-guided chemical elaboration of hits led to the development of compounds with potent *in vitro* TrmD inhibitory activity. Several of these compounds exhibit activity against planktonic *M. abscessus and Mycobacterium tuberculosis*. The compounds were further active in macrophage infection models against *Mycobacterium leprae* and *M. abscessus* suggesting the potential for novel broad-spectrum mycobacterial drugs.

## Introduction

Mycobacteria are a group of diverse organisms that include many important human pathogens. Within this group, *Mycobacterium tuberculosis*, the causative agent of tuberculosis, is responsible for over 1.7 million deaths per year (Floyd et al., 2018). Nevertheless, *Mycobacterium abscessus*, a rapidly growing species of nontuberculous mycobacteria (NTM), has recently emerged as a major threat to individuals with Cystic Fibrosis (CF) and other chronic inflammatory lung conditions (Bar-On et al., 2015; Sood and Parrish, 2017), with infection rates increasing around the world (Floto et al., 2016). *M. abscessus* is intrinsically resistant to most antibiotics and is consequently associated with extremely high treatment failure rates (Floto et al., 2016). There is therefore an urgent unmet need to develop new antibiotics.

Several structurally diverse, modified nucleosides, found at different locations of tRNAs, help in the maintenance of the reading frame and avoidance of translational frame-shift errors. Many such nucleoside modifications are found in regions near the anticodon, particularly at position 34 (the wobble position) and 37 (3’ and adjacent to the anticodon) of tRNA (Ahn et al., 2003; Urbonavicius et al., 2001). TrmD, tRNA-(N(1)G37) methyltransferase, catalyzes the methylation of G_37_ (Guanosine at position 37) in prokaryotic tRNAs **(Figure 1A)**. This modified nucleotide N^1^-methylguanosine at position 37 (m^1^G_37_) is present in tRNAs containing a G_36_G_37_ sequence in the anti-codon region from all three domains of life, where G_37_ is the base adjacent to the anticodon at the 3’ end (Ahn et al., 2003; Bjork et al., 2001; Bjork et al., 1989). Mutations in *trmD* result in growth defects associated with increased translational frameshifting leading to defective protein production (Bjork et al, 1989; Urbonavicius, 2001). TrmD belongs to a distinct class of *S*-adenosyl-L-methionine (SAM)-dependent methyltransferases known as the SpoU-TrmD (SPOUT) RNA methyltransferase superfamily or Class IV methyltransferases. Proteins belonging to this family are structurally unique due to the absence of a consensus methyltransferase fold. TrmD and other proteins of the SPOUT family consist of a deep trefoil knot architecture at the catalytic region, which provides an L-shaped pocket for binding of SAM. In eukaryotes however, G_37_ methylation is carried out by the enzyme Trm5, belonging to the Class I methyltransferase family (Anantharaman et al., 2002; Hori, 2017; Ito et al., 2015).

**Figure 1:**
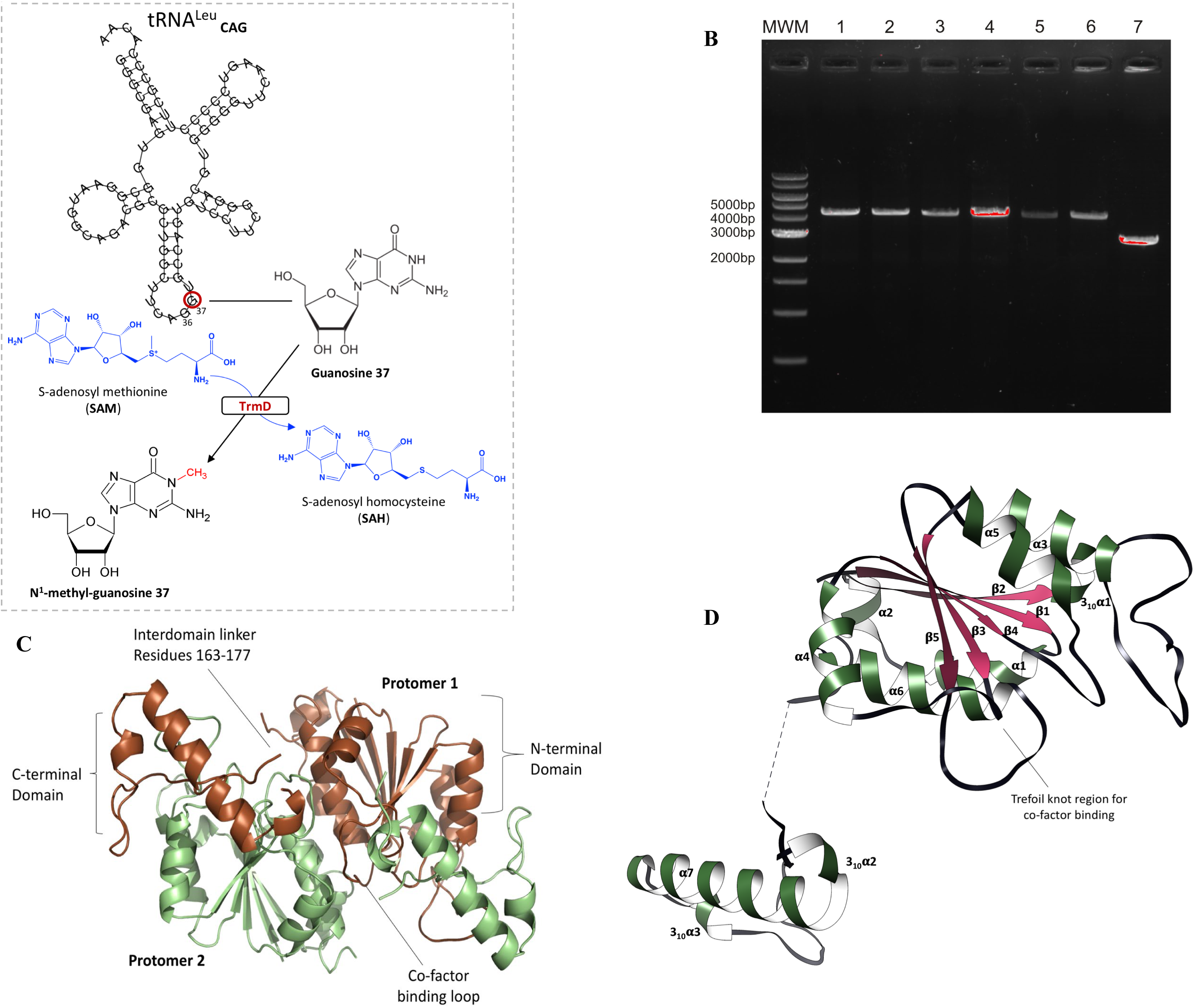
**A)** TrmD reaction scheme illustrated with tRNA^Leu^CAG as substrate. The methylation of G37 base adjacent to the anti-codon region leading to N1-methyl-guanosine 37 is mediated by methyl donor *S*-adenosyl-L-methionine, which in turn gets converted into *S*-adenosyl-L-homocysteine **B)** Allelic replacement at the *trmD* locus of *M. abscessus*. Candidate *trmD* mutants were analysed by PCR using a set of primers annealing outside the allelic exchange substrate. Lanes 1 to 6 show *M. abscessus subsp. massiliense* CIP108297 clones harboring the pMV306H::*trmD* plasmid whose endogenous chromosomal *trmD* locus was replaced by the Str cassette (expected size of the PCR fragment is 4,200 bp). Lane 7 shows the amplification product of the *trmD* locus in wild-type *M. abscessus subsp. massiliense* CIP108297 (expected size of the PCR fragment is 2,571 bp). MWM, molecular weight marker. **C)** TrmD homodimer (PDB code 6NVR) with domain architecture illustrated. Protomers 1 and 2 are represented in brown and green ribbon diagrams respectively. **D)** TrmD protomer is coloured and illustrated based on secondary structure elements. The disordered inter-domain linker is shown as black dotted lines.

Previous research (Goto-Ito et al., 2009; Ito et al., 2015) has shown that TrmD and Trm5 have distinct substrate requirements with RNA. While Trm5 recognizes the overall L-shaped tertiary structure of tRNA possessing a G_37_ base, TrmD recognition involves mainly the D stem and anticodon stem loop of tRNA with G_36_G_37_ bases. Trm5 functions as a monomer and binds to SAM at the Rossmann fold region of the active site, in contrast to dimeric TrmD with a trefoil knot methyl donor binding region. Further, SAM adopts a unique bent conformation in TrmD as compared to the extended conformation in Trm5 and many other canonical methyltransferases. These distinct structural features, substrate requirements and ligand binding conformations provide the potential for designing novel and selective inhibitors of bacterial TrmD (Goto-Ito et al., 2017).

A previous drug discovery effort targeting *Haemophilus influenzae* TrmD (Hill et al., 2013) led to the development of selective inhibitors with potent biochemical activity against TrmD isozymes *in vitro*. However, these compounds in general only showed weak antibacterial activity when profiled against a range of Gram-positive and Gram-negative pathogens, including against recombinant strains of *E. coli* and *H. influenzae* debilitated in the AcrAB TolC efflux pumps (Hill et al., 2013). A recent drug discovery work was reported against *Pseudomonas aeruginosa* TrmD which reported potent inhibitors of this enzyme. However, the antibacterial activity of these was weak not only against *P. aeruginosa* but also against other bacteria including mycobacteria (Zhong et al., 2019).

Fragment-based drug discovery (FBDD) is a promising approach for the identification of new drugs, whereby the complexity of the chemicals screened is reduced by decreasing their molecular weights (typically < 300 Da), which at the same time increases their promiscuity in binding targets. Initial fragment hits usually exhibit lower potency than the more complex drug-like molecules found in typical high-throughput screening compound libraries. However, such fragments bind by making well-defined and directional interactions, giving rise to highly ligand efficient (LE) molecules. These fragments can then be chemically optimized into lead candidates, thereby more effectively exploring the chemical space available for binding to the target protein (Erlanson et al., 2016; Mendes and Blundell, 2016; Murray et al., 2014; Thomas et al., 2017). In this work we validate TrmD as a mycobacterial target and describe the application of an FBDD approach to generate a new family of small-molecule inhibitors of *M. abscessus* TrmD, having anti-microbial activities against a range of pathogenic mycobacteria.

## Results

### TrmD is essential for *M. abscessus*

Previous studies have demonstrated that *trmD* is essential in diverse bacteria including *M. tuberculosis*, where different transposon mutagenesis studies have shown that it is essential or that mutants had a growth defect (DeJesus et al., 2017; Griffin et al., 2011; Sassetti et al., 2003). However, confirmation of essentiality of *trmD* for *M. abscessus* was lacking. Three initial attempts to disrupt the *trmD* gene of *M. abscessus* subsp. *massiliense* CIP108297 by homologous recombination using a recombineering approach proved unsuccessful. To determine whether the *trmD* gene is required for *M. abscessus* growth, allelic replacement experiments were repeated in the background of a *M. abscessus* subsp*. massiliense* CIP108297 merodiploid expressing a second copy of the *trmD* gene from the integrative plasmid pMV306H::*trmD*, and in a control *M. abscessus* subsp*. massiliense* CIP108297 strain harbouring an empty pMV306H plasmid. Analysis of over 100 candidate mutants in each background from two independent experiments showed that the endogenous chromosomal copy of *trmD* could be knocked-out in the presence of an extra-copy of the gene but not in *M. abscessus* cells carrying an empty plasmid (**Figure 1B**), thus confirming *trmD* essentiality for *M. abscessus* growth.

### *M. abscessus* TrmD: overall structure and ligand binding

We determined the crystal structures of *M. abscessus* TrmD in apo form at 1.60 Å resolution (PDB code **6NVR**), as well as in complex with SAM and *S*-adenosyl-L-homocysteine (SAH) at 1.67 Å and 1.48 Å resolution respectively (PDB codes **6NW6** & **6NW7**). The crystals belong to space group P2_1_2_1_2_1_ and consist of a homodimer in the asymmetric unit. Each non-crystallographic two-fold symmetry-related protomer of TrmD interacts in an antiparallel manner and consists of two domains: a larger *N*-terminal domain spanning residues 1-161 and a smaller *C*-terminal helical domain (177-242) connected by a flexible inter-domain linker. The two domains of the individual protomers do not contact each other and the inter-domain region is largely disordered, with residues 162-177 not clearly visible in the apo structure (**Figure 1 C & D).**

The SAM binding region of TrmD is located at the base of the *N*-terminal domain and consists of a deep trefoil knot architecture, made of three distinct untwisted loop regions. The trefoil knot of *M. abscessus* TrmD is made up of a cover loop spanning residues ^84^TPAG^87^ between strand β3 and helix α4 leading to the wall loop at the edge of the methionine pocket containing residues ^109^GRYEGID^115^ between β4 and helix α5. This loop then crosses over to form the bottom loop with residues 132-140 that encompasses the SAM adenine ring between strand β5 and helix α6 **(Figure 2 A & B).** SAM and SAH occupy the deep trefoil-knot active site at the base of the *N*-terminal region and adopt an L-shaped bent conformation as previously observed with other TrmD orthologs (Christian et al., 2016; Koh et al., 2017). Both SAM and SAH form an extensive hydrogen-bonding network in this region along with hydrophobic and π-interactions as shown in **Figure 2 B & C**. The adenine ring of SAM and SAH is sandwiched between the cover loop and bottom loop of the knot with the adenine N1 and N7 forming hydrogen-bond contacts with the backbone amide-nitrogen atoms of Ile133 and Leu138 respectively, while the amino nitrogen forms additional hydrogen bonding contacts with the backbone carbonyl oxygen atoms of Gly134 and Tyr136 of the bottom loop (**Figure 2 B & S2**).

**Figure 2:**
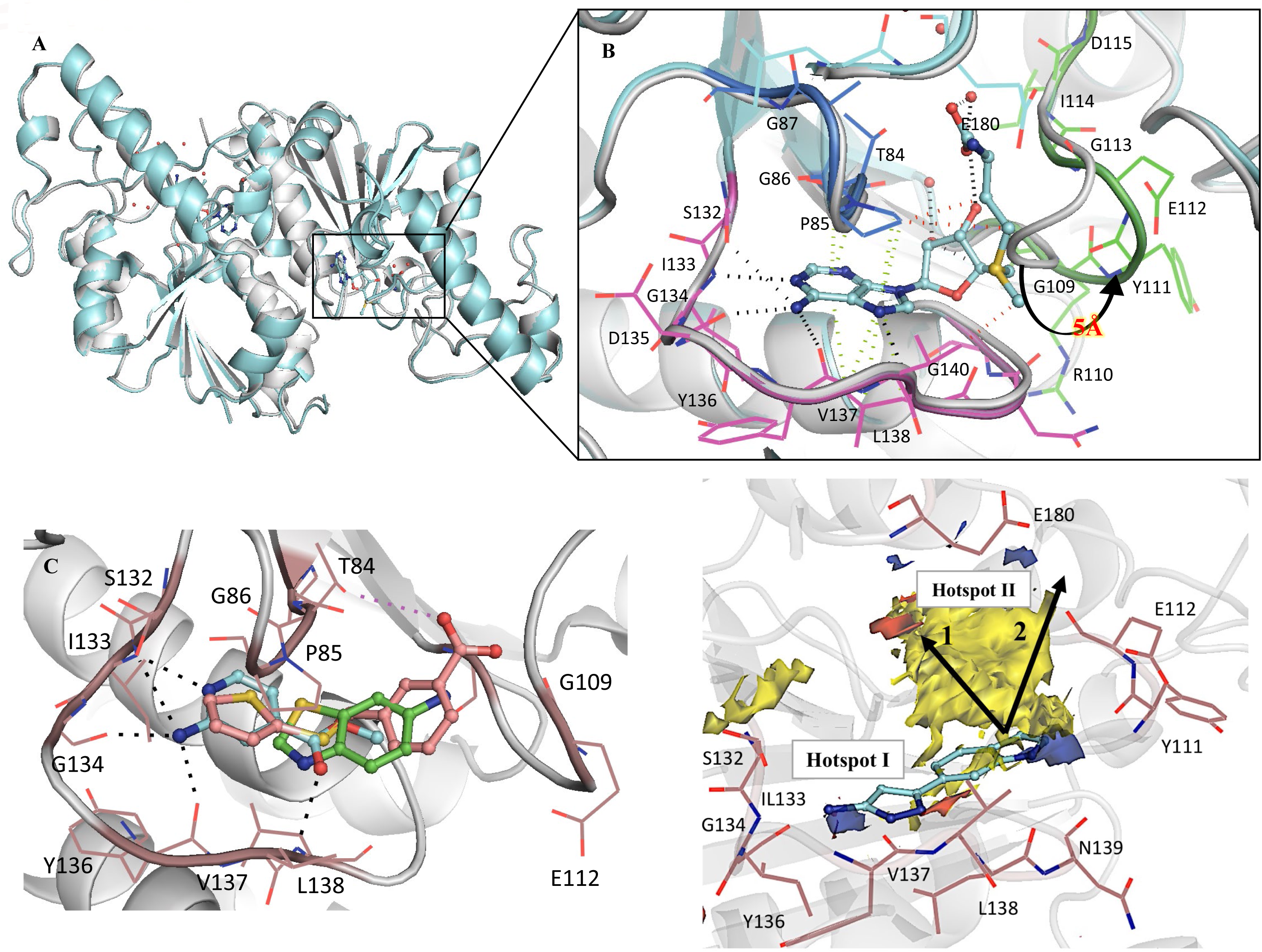
**A)** Structural superposition of TrmD apo form (white) and TrmD SAM bound form (light blue), PDB codes 6NVR and 6NW6 respectively **B)** the trefoil-knot active site of TrmD involving: cover loop (residues 84-87) shown in dark blue and bottom loop (residues 132-140) in magenta and wall loop (residues 109-115) in green and the conformational flip of the wall loop (residues ^109^GRYEGID^115^) upon SAM (light blue stick) binding are illustrated. The residues corresponding to each loop region are also shown as line representation**. C)** Three representative fragment hits from each cluster, fragment **14** (PDB Code 6QOK) coloured in blue, fragment **20** (PDB Code 6QOQ) in green and fragment **8** (PDB Code 6QOE) in salmon respectively, occupying the TrmD SAM binding site. Major Hydrogen bonds and electrostatic interactions are depicted in black and purple dotted lines respectively. **D)** Hot spot map contoured at 14 of TrmD active site superposed with crystal structure of TrmD in complex with merged compound **AW1** (light blue stick). Donor, acceptor and hydrophobic regions of the map are depicted as blue, red and yellow regions respectively. Amino acid residues contributing towards interactions in each hotspot map region are shown as brown stick representation. The arrows indicate two potential ways of fragment elaboration.

The ribose and methionine moieties of SAM and SAH interact with the wall and bottom loops of the knot. The hydroxyl oxygen atom (O2’) of the ribose ring forms a hydrogen bond with the backbone amide of Gly109. The methionine and homocysteine moieties further extend into the active site groove formed between the cover and wall loops, making further hydrogen-bonding interactions with water molecules in this region (**Figure 2 B & S2**).

A structural superposition of the apo and SAM bound forms of TrmD reveals the wall loop undergoing a switch in conformation leading to a movement of about 5 Å, when measured at the Cα of Tyr111, to the outer edge of subunit A. This conformational flip of the wall loop from apo form to SAM-bound form and the subsequent change in positions of residues 110 to 113 help to accommodate the methionine moiety of the methyl donor (**Figure 2 A & B**).

### Fragment Screening, hit validation and clustering of fragments

Having examined the conformational changes and binding interactions at the *M. abscessus* TrmD catalytic site, we initiated a structure-guided FBDD effort targeting *M. abscessus* TrmD by screening an in-house library of 960 small molecule fragments. The preliminary screening was performed using differential scanning fluorimetry (DSF), resulting in 53 hits within a thermal shift cut-off value of 3 standard deviations from the negative control, consisting of TrmD protein in the absence of any ligand. These hits were then selected for validation by X-ray crystallography. Apo crystals of *M. abscessus* TrmD were soaked with each of the 53 fragments in individual experiments. The resulting crystal structure determinations allowed characterization of the binding modes of 27 fragments (**Figure S3**).

All of the 27 fragments, validated by X-ray crystallography, were found to occupy the TrmD SAM site. These fragments can be clustered into three groups based on their binding mode at this site (**Figure 2 D & S3**). **Cluster 1** consists of 12 fragments that bind exclusively to the sub-pocket that accommodates the adenine ring of SAM, engaging residues within the cover and bottom loops of the trefoil knot. These fragments recapitulated many of the hydrogen bonding and π-interactions of the SAM adenine moiety, as shown in the example (**Figure 2 D & S3)**. These interactions include hydrogen-bond contacts to the side chain of Ser132, which in turn adopts a dual conformation, and to the backbone amides of Ile133, Gly134, Tyr136 and Leu138.

The **second cluster** consists of 12 further fragments that occupy the entire adenosine region of the TrmD active site, thus extending from adenine towards the ribose-binding pocket of the active site. These fragments, in addition to retaining several adenine moiety contacts, also interact with the wall loop residues, forming hydrogen bonds to the backbone amides of Tyr111 and Gly109 and water-mediated hydrogen bonds as shown in the example (**Figure 2 D & S3). Cluster 3** consists of three fragments that extend beyond the TrmD adenosine site, thus reaching the methionine-binding region of the pocket. One of these fragments, fragment **8** stretched further into the groove formed between the cover and wall loops of the trefoil knot, thus engaging additional hydrogen bonding contacts with the side chain of Thr84 and the back bone amide of Gly109 in this region (**Figure 2 D & S3).**

### Fragment merging & hotspot mapping for chemical elaboration

Two of the above 27 fragment hits, fragments **23** (*K*_d_ 0.17 mM, LE 0.37) and **24** (*K*_d_ 0.26 mM, LE 0.41), were chosen for further chemical development by a fragment-merging strategy. The choice of fragments for subsequent chemical optimization was based on a number of criteria, including binding affinity, ligand efficiency, synthetic tractability and their ability to make key binding interactions at the TrmD SAM binding site. Fragment **23** occupies the adenosine binding region of the TrmD AdoMet site, with its pyrazole ring making hydrogen bond contacts to the backbone amides of Tyr136 and Leu138 and the amino group making further hydrogen bonds with the backbone carbonyl oxygen of Gly134 and the side chain of Ser132, respectively. The 4-methoxyphenyl ring of the fragment extends into the ribose binding site, engaging hydrophobic and π-interactions with the residues of the cover loop (**Figure 3A & S3).**

**Figure 3:**
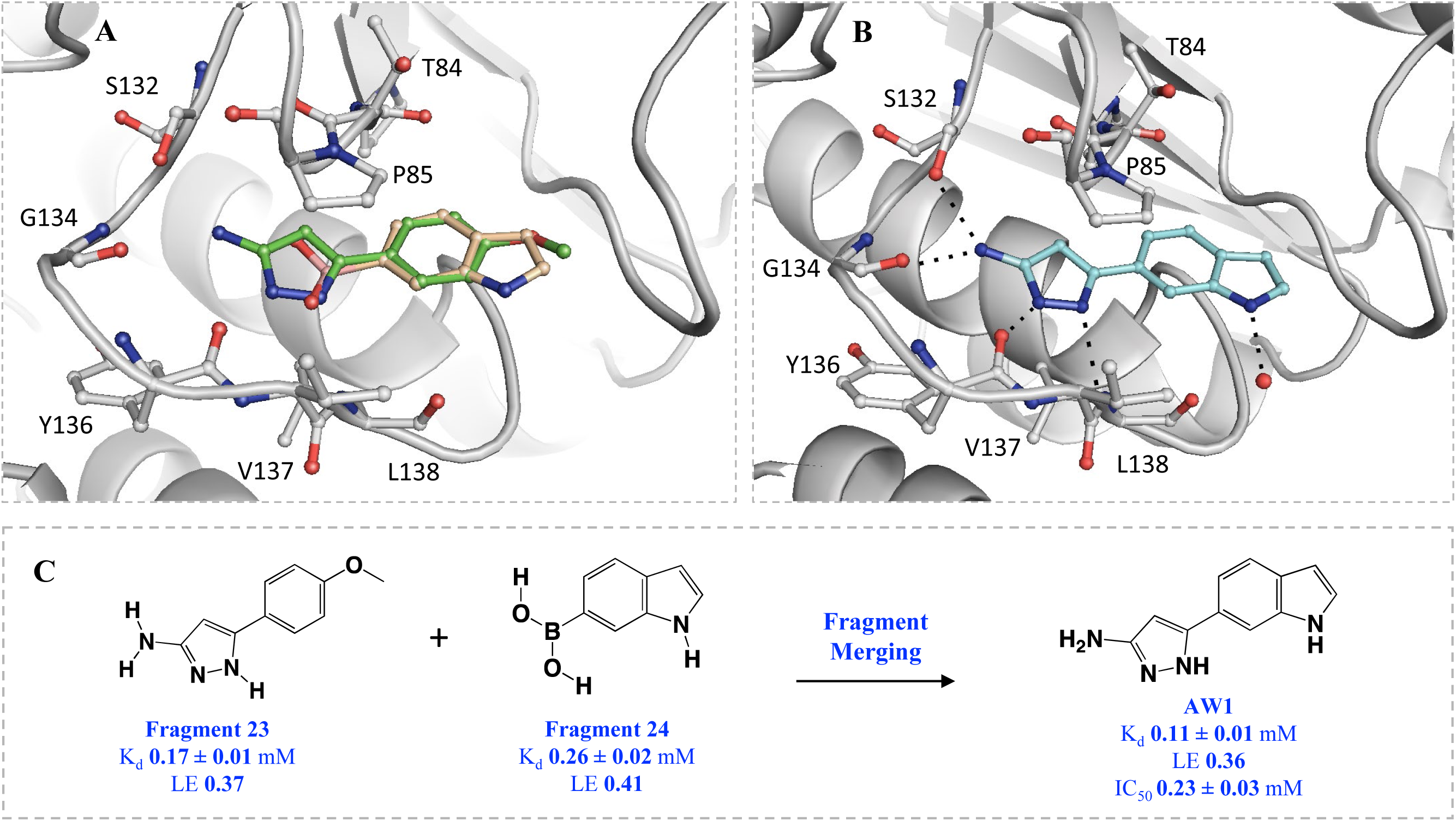
Fragment merging approach. **A)** Structural superposition of TrmD (grey) in complex with fragments **23** (green) and **24** (beige) – PDB Codes 6QOT & 6QOU, showing binding mode and interactions at the SAM site **B)** merged compound **AW1** (light blue), PDB Code 6QQS, showing binding mode at the SAM site. The corresponding amino acid interactions are illustrated in dotted lines and **C)** the overall scheme of fragment merging.

The indole ring of fragment **24** also occupies the ribose pocket where it forms a water-mediated interaction with the backbone amide nitrogen of Leu138. The 4-methoxyphenyl and indole ring systems of fragments **23** and **24** overlap perfectly, while the 6-boronic acid group of fragment **24** partially extends into the SAM adenine pocket and makes hydrogen bonds with the backbone amides of residues Tyr136 and Leu138 and further water-mediated hydrogen bond contacts to the backbone amides of Val131, Ile133, Gly134 and the side chain hydroxyl group of Ser132 (**Figure 3A & S3**). Compound **AW1** (*K*_d_ 0.11 mM, LE 0.36, IC_50_ 0.23 mM), formed by merging the two fragments, adopts a similar conformation to that of the original fragments in the TrmD SAM site, as shown in **Figures 3 B & C**, thereby providing a new chemical scaffold for further structure-guided development.

To aid the structure-guided lead discovery, the binding propensities of TrmD protein to ligands were further examined using the hotspot-mapping program developed by Radoux and co-workers (Radoux et al., 2016). Hotspots are areas within the protein that provide relatively large contributions to the overall binding affinity of ligands (Hajduk et al., 2005; Ichihara et al., 2014). This is usually mediated by the displacement of water molecules having restricted freedom owing to their location within a hydrophobic cavity or close to a patchwork of hydrogen bonds and lipophilic amino acid side chains, thus compensating for the loss of entropy on binding. These regions not only satisfy the minimum binding requirement for fragments but also maintain the original fragment binding interactions when elaborated (Radoux et al., 2016). While the observed fragment hits and the corresponding merged compound **AW1** satisfy many of the predicted protein-hotspot interactions, the map suggests further interactions that stabilise elaborated fragments, sometimes allowing them to reach other hotspots that are not yet explored. As shown in the example (**Figure 3 D**), **AW1** occupies hotspot 1 at the base of the TrmD active site, where it satisfies the hydrogen-bond donor requirements by interacting with the backbone amide oxygen atoms of Gly134 and Tyr136. The merged compound **AW1** also orients its pyrazole nitrogen atom in the acceptor map in this region where it forms a hydrogen bond with the backbone NH of Leu138 (**Figure 3 D**). The compound could be elaborated further towards the methionine end of the active site and by further extension to the second hotspot region at the top of the active site. The second hotspot is characterized by a large hydrophobic patch surrounded by the acceptor region mediated by the backbone amide group and side chain of Glu180. A second approach to fragment elaboration is by growing further upwards from the hydrophobic region of hotspot 2 over to the donor region mainly mediated by the backbone oxygen atom and side chains of Glu112 as illustrated in **Figure 3 D**.

### Structure-based lead optimization of merged compounds

A structure-guided elaboration of the merged compound **AW1** (*K*_d_ 0.11 mM, LE 0.36, IC_50_ 0.23 mM) was carried out, initially utilizing the indole nitrogen as a vector for growth. The addition of a 2-picolyl moiety successfully increased the affinity of **AW1** by an order of magnitude in compound **AW2** (*K*_d_ 12 µM, LE 0.30, IC_50_ 33 µM) (**Table 1**). The methylene linker attached to the indole nitrogen of **AW2** allowed the added pyridyl ring to occupy the region defined by Pro85, Glu112, Val137, Arg154 and Glu180 **(Figures 3E & 4A).**

**Table 1:** Summary of structure-guided optimization of merged compound AW1

| Compound | Structure | | $K_d$ (μM) | LE (kcal/ mol/<br>heavy atom) | $IC_{50}$ (μM) |
| --- | --- | --- | --- | --- | --- |
| | $R^1$ | $R^2$ | | | |
| AW1 | H | H | $110 \pm 11$ | 0.36 | $230 \pm 29$ |
| AW2 | | H | $12 \pm 1$ | 0.30 | $33 \pm 4$ |
| AW3 | | CN | $0.50 \pm 0.14$ | 0.36 | $0.31 \pm 0.01$ |
| AW4 | | CN | $0.092 \pm 0.018$ | 0.32 | ND |
| AW5 | | CN | $0.027 \pm 0.004$ | 0.34 | $0.030 \pm 0.003$ |
| AW6 | | H | $0.49 \pm 0.21$ | 0.31 | $1.4 \pm 0.1$ |
| AW7 | | CN | $0.073 \pm 0.030$ | 0.30 | $0.069 \pm 0.007$ |

**Figure 4:**
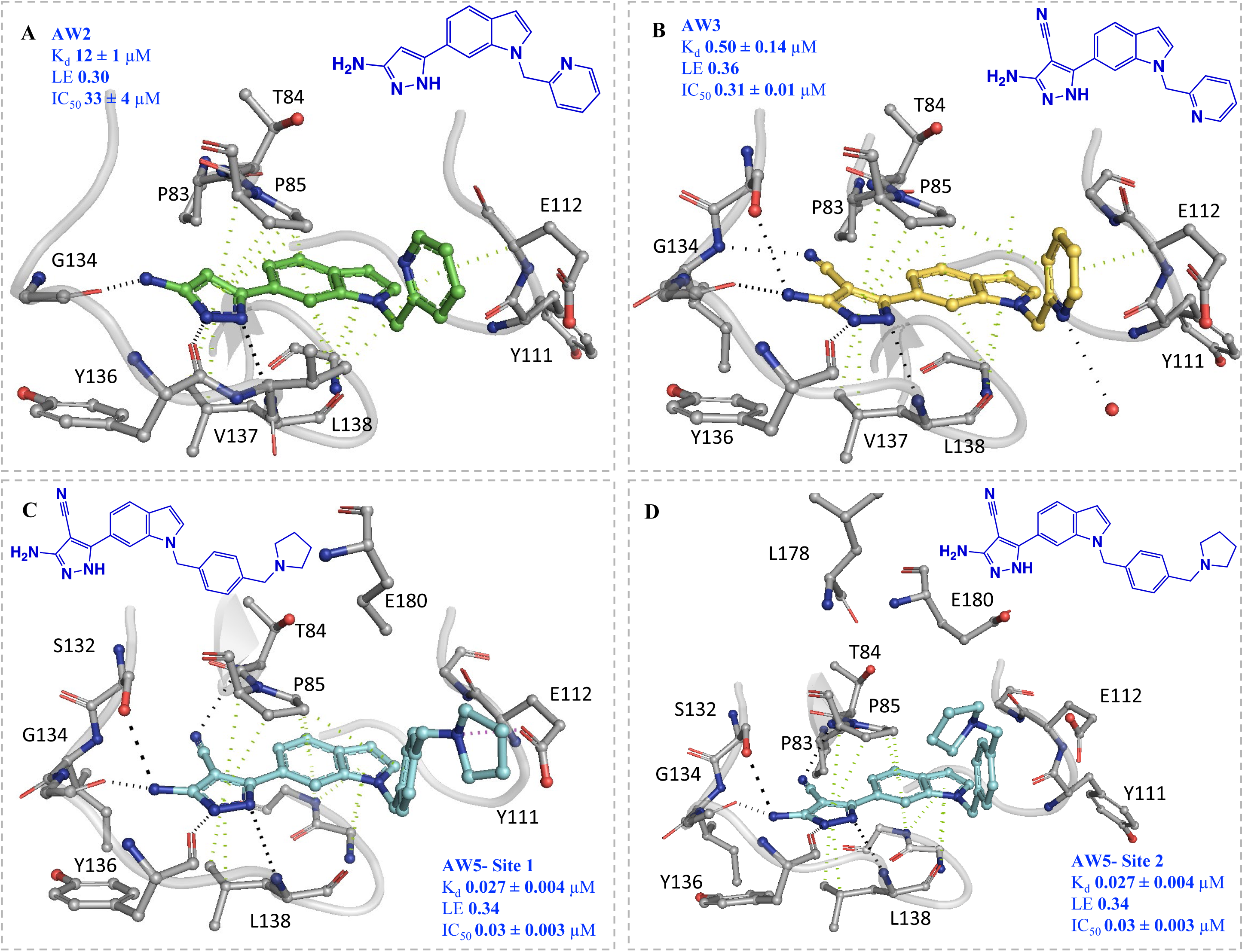
X-ray crystal structure of TrmD (grey) in complex with compounds **A) AW2** (green stick), PDB Code 6QQX **(B) AW3** (yellow stick), PDB Code 6QQY **(C) AW5** at protomer 1, (blue stick), PDB Code 6QR6 and **(D) AW5** at protomer 2 (blue stick), PDB Code 6QR6, showing binding mode at the SAM site. The corresponding amino acid interactions are illustrated with Hydrogen bonds, π-interactions and electrostatic contacts depicted in black, green and purple dotted lines respectively.

The affinity of **AW2** was further improved by the addition of a nitrile group on the 4-position of its pyrazole ring, extending into the narrow space between residues ^83^PTP^85^ of the cover loop and ^131^VSI^133^ of the bottom loop respectively **(Figure 4 A & B). AW3** (*K*_d_ 0.50 µM, LE 0.36, IC_50_ 0.31 µM) was a significant improvement on **AW2**, with the addition of two heavy atoms affording a 25-fold decrease in K_d_ (12 to 0.50 µM), increasing the ligand efficiency to the level of the original merged compound **AW1** (0.36), and a 100-fold decrease in IC_50_ (33 to 0.31 µM) (**Table 1**). The X-ray crystal structure of TrmD in complex with **AW3** shows that the original fragment contacts have been retained, with the **AW3** aminopyrazole ring orienting itself in a similar manner to **AW1** and retaining its hydrogen bonding contacts to the side chain of Ser132 and backbones of Gly134, Tyr136 and Leu138. In addition, the nitrile group of **AW3** seems to have strengthened the interactions at the active site region between residues ^83^PTP^85^ and ^131^VSI^133^ by engaging in an additional hydrogen bond contact with the backbone NH of Ile133 **(Figure 4B).**

Further elaboration was carried out from the 5-position of the pyridyl ring of **AW3** through the attachment of a pyrrolidinyl ring via another methylene linker, with **AW4** (*K*_d_ 92 nM, LE 0.34) affording an additional 5-fold improvement in affinity (**Table 1**). Modification of the scaffold of **AW4** by replacement of its pyridyl ring with a phenyl ring in **AW5** (*K*_d_ 27 nM, LE 0.34, IC_50_ 30 nM) was tolerated with a greater than three-fold improvement in affinity (92 to 27 nM) and an increase in ligand efficiency (0.32 to 0.34). The X-ray crystal structure of **AW5** shows the pyrrolidinyl ring occupying the binding site in two conformations, depending on the active site, thereby engaging either Glu112 or Glu180 in an electrostatic interaction **(Figure 4 C & D).** The removal of the nitrile group on the pyrazole ring of **AW5** in compound **AW6** (*K*_d_ 0.49 µM, LE 0.31, IC_50_ 1.4 µM) **(Figure 5A)**, had a detrimental impact on both affinity and performance in the biochemical assay, demonstrating the importance of extension of this substituent into the cavity between residues ^83^PTP^85^ of the cover loop and ^131^VSI^133^ of the bottom loop. Exploration of the active site region bordered by the Ala176 to Glu180 loop through replacement of the pyrrolidinyl ring of **AW5** with an *N*-methyl piperazinyl motif in **AW7** (*K*_d_ 73 nM, LE 0.30, IC_50_ 69 nM) showed a slight worsening of both affinity and IC_50_, possibly due to the slight change (0.3 Å) in the position of the nitrile group in comparison to that of **AW5**, thereby diminishing the hydrogen bonding contact with the backbone amide of Thr84 (**Figure 5 B** & **S8**).

**Figure 5:**
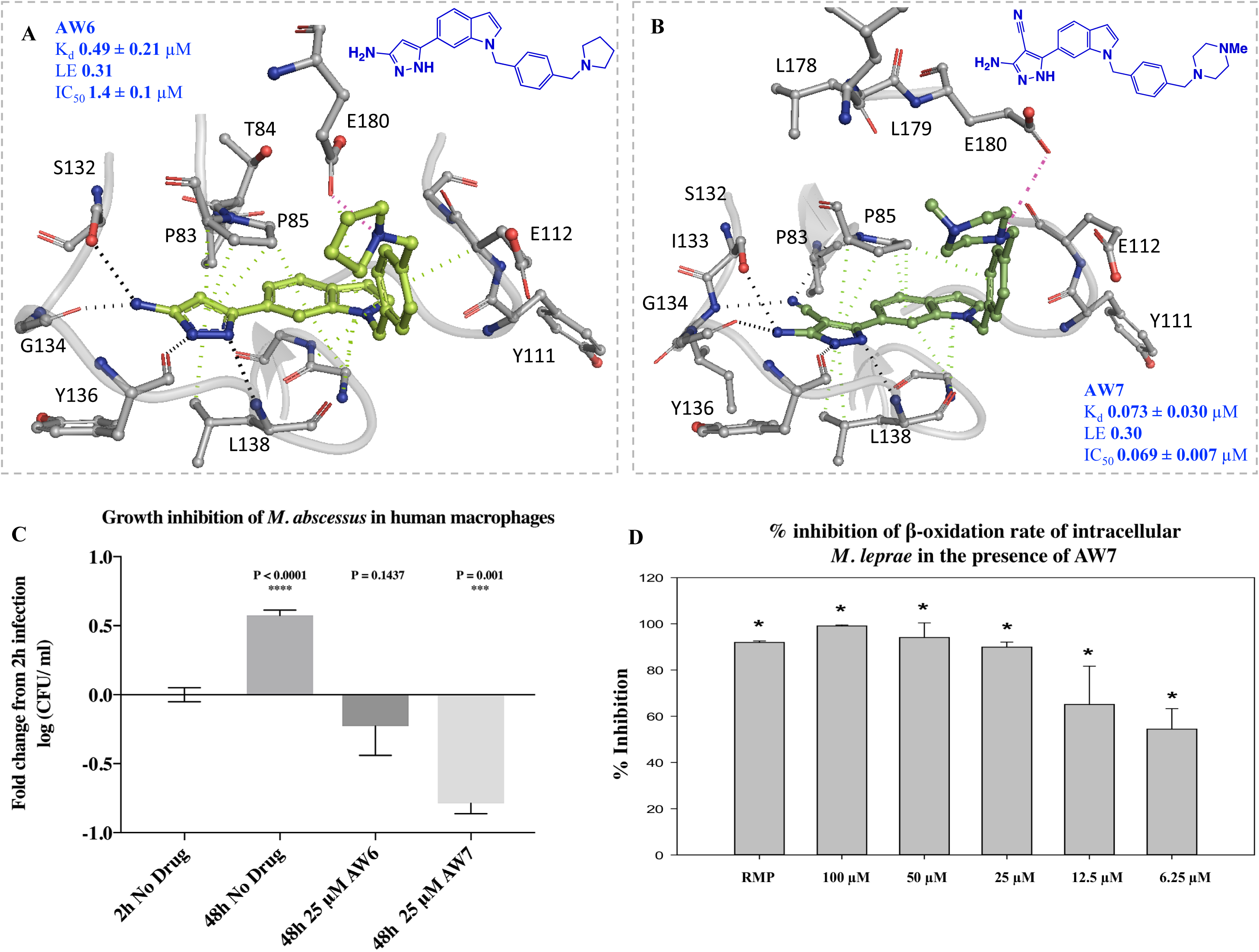
X-ray crystal structure of TrmD (grey) in complex with compound **A) AW6** (lime), PDB Code 6QR5 **B) AW7** (green), PDB Code 6QR8, showing binding mode at the SAM site. The corresponding amino acid interactions are illustrated with Hydrogen bonds, π-interactions and electrostatic contacts depicted in black, green and purple dotted lines respectively. **C)** Growth inhibition study of lead compounds **AW6** & **AW7** in *M. abscessus* infected human macrophages over a 48 h period. Data represented as fold change in log (CFU/ml) from 2 h infection in the absence of any drug. Fold change in log (CFU/ml) following 48 h infection with no drug and with 25µM **AW7** are statistically significant (P<0.05) in comparison to that in the presence of **AW6** 48 h post infection (P = 0.1437) implying that **AW6** is bacteriostatic and **AW7** is bactericidal for *M. abscessus*. **D)** Intracellular *M. leprae* palmitic acid oxidation rate (radiorespirometry) in the presence of different concentrations of **AW7** for 7 days. 7^th^ day cumulative counts per minute (CPM) were recorded and percentage inhibition of metabolism determined as compared to no drug control. **AW7** concentrations, in mM, are shown in parenthesis and rifampin (RMP) was used at 2.4 mM. * - Inhibition is statistically significant (P<0.05) compared to no drug control.

### TrmD lead compounds have anti-mycobacterial activity

The compounds were further examined for their ability to inhibit bacterial growth. While the initial fragment hits of TrmD and the early stage compounds elaborated from the fragments exhibited low levels of growth inhibition up to 250 µM (data not shown), compounds in later stages of development showed promising activity against *M. abscessus* and *M. tuberculosis* (**Table 2**). The MICs of the TrmD lead compounds against *M. abscessus* were identical and most of the lead molecules exhibited much higher inhibition against *M. tuberculosis* than *M. abscessus* (**Table 2**). Surprisingly, **AW6,** in which the nitrile group of **AW5** was removed, despite being the least active lead compound in the *in vitro* TrmD assays, showed similar MICs when compared to the other two compounds. Additionally, the replacement of the pyrrolidinyl ring of **AW5** with an *N*-methyl piperazinyl motif in **AW7** afforded a 2-fold improvement in the MIC compared to **AW5** against *M. tuberculosis*, however this wasn’t observed in *M. abscessus* (**Table 2**). Sub-micromolar affinities were subsequently determined for **AW6** (*K*_d_ 0.90 µM) and **AW7** (*K*_d_ 0.33 µM) with TrmD from *M. tuberculosis* (**Figure S18** and **S19**), supporting both the MIC values against *M. tuberculosis* and the applicability of this lead series to TrmD homologs from mycobacteria other than *M. abscessus*. The lead molecules **AW6** and **AW7** were then tested against a wider panel of NTMs (**Table 2**). The obtained MICs revealed that the compounds display limited inhibitory activity across most NTMs tested except for *Mycobacterium terrae*, where MICs are better than for *M. abscessus*. Given the high percentage sequence identity of TrmD across mycobacterial species **(Figure S1)**, it was expected that growth inhibition would be observed against some of these organisms. However, the variation in MIC observed for our lead compounds is in line with the variation in drug susceptibility profiles between mycobacterial species due to differential permeability, retention, and metabolism of compounds (Li et al., 2013; Scherr et al., 2016).

**Table 2:** Minimum inhibitory concentration (MIC) values of TrmD compounds across various mycobacterial species and strains

| No | MIC (μM) | | | | | | | |
| --- | --- | --- | --- | --- | --- | --- | --- | --- |
|  | *M. abscessus*<br>19977 | *M. tuberculosis*<br>Bluepan | *M. tuberculosis*<br>H37Rv | *M. chelonae* | *M. fortuitum* | *M. goodii* | *M. terrae* | *M. avium* |
| AW5 | 50 | 50 | 12.5 | ND | ND | ND | ND | ND |
| AW6 | 50 | 25 | 6.3 | 100 | 100 | 100 | 100 | 200 |
| AW7 | 50 | 25 | 6.3 | 100 | 100 | 100 | 25 | 100 |

### TrmD lead compounds kill intracellular *M. abscessus* and *M. leprae*

We assessed the cytotoxicity effect of **AW6** and **AW7** using lactate dehydrogenase (LDH) release assay. At or below 150 µM, neither compound caused cellular toxicity on human macrophages (**Figure S5).** The compounds were then evaluated in *M. abscessus*-infected human macrophages. Both compounds showed activity in the macrophage infection model, with **AW7** performing better than in the *in vitro* assays. At 25 µM **AW6** showed a ∼82% decrease in CFUs while **AW7** at the same concentration showed a 95% reduction in CFUs compared to the no drug control after 48h incubation (**Figure 5C**). However, only **AW7** demonstrated a bactericidal effect with a −0.8 log change after 48h incubation (**Figure 5C**).

The best lead molecule (**AW7**) was further tested against *M. leprae* maintained intracellularly in murine bone marrow macrophage. Relative inhibition of β-oxidation rates were measured using a radiorespirometry assay after 7 days of incubation. The results show that **AW7** at 6.2 µM was able to inhibit *M. leprae* radiorespirometry by ∼54% when compared to the no-drug control and by ∼89% at 25 µM, which is similar to rifampicin inhibition at 2.4 µM (91%) (**Figure 5D**). Axenically-maintained *M. leprae* showed only a 15% reduction of radiorespirometry at 25 µM and no effect at 6.2 µM, following a similar trend to what was observed for *M. abscessus* with the highest activity for the compound being observed in the macrophage infection model (**Figure 5C, S6 & S7**).

## Discussion

TrmD (tRNA-(N(1)G37) methyltransferase) is an essential tRNA modification enzyme in bacteria that prevents translational frame-shift errors by methylation of a guanosine base at position 37 of tRNAs containing G_36_G_37_ bases at the anti-codon region. This enzyme was found to be essential in *M. tuberculosis* and other organisms but direct confirmation for *M. abscessus* was lacking. This work has confirmed the essentiality of *trmD* for *M. abscessus* subsp*. massiliense* growth, validating TrmD as a drug discovery target for this organism.

A FBDD effort targeting *M. abscessus* TrmD was undertaken by screening a library of 960 fragments by DSF and resulted in 53 preliminary hits. Of these, 27 were subsequently validated by X-ray crystallographic studies of the TrmD-fragment complexes. The resulting fragments can be classed into three clusters corresponding to their different binding modes in the TrmD SAM binding site. The determination of high-resolution crystal structures informed structure-guided drug discovery and allowed us to select and develop a series of compounds by merging fragments **23** and **24**. Chemical elaboration of the merged compound **AW1** allowed the synthesis of potent *M. abscessus* TrmD inhibitors. This was achieved through both the addition of a nitrile group on the 4^_^position of the pyrazole ring and elaboration from the indole nitrogen. This elaboration led to several low nanomolar *M. abscessus* TrmD inhibitors demonstrating a significant improvement of *in vitro* affinity from the initial fragments.

The lead compounds reported in this work exhibited bactericidal effects on *M. abscessus*, *M. tuberculosis*, and *M. terrae*. Furthermore, compound **AW7** showed bactericidal intracellular activity against *M. abscessus* and was also a potent inhibitor of intracellular *M. leprae*. The compounds reported here are the first TrmD inhibitors reported in the literature with strong bactericidal activity against mycobacteria. However, we observe a poor correlation between K_d_/ IC_50_ and MIC for the lead compounds, with **AW6**, the weakest of the lead compounds, presenting similar MIC values to the other compounds against most of the species tested. Furthermore, whilst the compounds have nanomolar affinities against TrmD from *M. abscessus and M. tuberculosis*, and likely similar affinities for other mycobacterial TrmD given the very high sequence identity shared between the TrmD orthologues (**Figure S1**), their MICs are ∼35-3000 fold worse across the NTMs tested. This poor correlation and variation across different mycobacterial species suggests differential effects of compound permeability, retention, and/or metabolism on *in vivo* activity. Similar results were found by others in a recent phenotypic screen of 129 TB active and 271 non-TB active compounds against *M. abscessus* which revealed that only a small subset (12 compounds) from the TB-active group and just one compound from the non-TB active group were effective against *M. abscessus* (Low et al., 2017). Nevertheless, our results with macrophage infection models, both with *M. leprae* and *M. abscessus*, show the potential of these molecules to be further optimized. The preliminary toxicity study of the lead molecules **AW6** and **AW7** using a lactate dehydrogenase (LDH) assay further showed no significant cytotoxicity on primary human macrophages.

Most currently-used antibiotics that target microbial protein synthesis act either by interacting with ribosomal sub-units (aminoglycosides, tetracyclines, macrolides etc) or *via* inhibiting mRNA synthesis (rifamycins) and elongation (actinomycin) (Kohanski et al., 2010). This study represents a proof of concept for the development of a new class of antibiotics targeting bacterial tRNA modification with potent bactericidal activities. Furthermore, the results presented in this work suggest the potential to develop novel mycobacterial drugs targeting bacterial tRNA methylation by TrmD.

## Materials & Methods

### Expression and purification of full length *M. abscessus* TrmD

*E. coli* BL21 (DE3) strain containing AVA0421 plasmid with an N-His-3C Protease site-TrmD full-length insert, kindly provided by the Seattle Structural Genomics Consortium, (Baugh et al., 2015) was grown overnight at 37 °C in LB-media containing Ampicillin (100 μg/mL). This seed stage culture was used to inoculate 6 shake flasks containing 1 L each of 2XYT media with Ampicillin (100 μg/mL) until optical density (A_600nm_) reached 0.6. The expression of recombinant construct was induced by the addition of Isopropyl β-D-1-thiogalactopyranoside (IPTG) to a final concentration of 0.5 mM and further allowed to grow at 18 °C for 16 h.

#### Isolation of cells & lysis

Cells were harvested by centrifugation at 4 °C for 20 min at 5000 g and the pellet was re-suspended in buffer A (25 mM HEPES pH 7.5, 500 mM NaCl, 5% Glycerol, 10 mM MgCl_2_, 1 mM TCEP, 20 mM Imidazole). 0.1% Triton (Sigma), 10 μg/mL DNaseI, 5 mM MgCl_2,_ and 3 protease inhibitor cocktail tablets (New England Biolabs) were added to the cell suspension. The cells were lysed in an Emulsiflex (Glen Creston) and clarified the lysate by centrifugation at 4 °C for 40 min at 25,568 g.

#### Immobilized Metal Affinity Chromatography

The clarified lysate was filtered using a 0.45 μm syringe filter and passed through a pre-equilibrated (with buffer A), 10 mL pre-packed nickel-sepharose column (HiTrap IMAC FF, GE Healthcare). The column was washed with 5 column volumes of buffer A and the bound protein was eluted as 4x 10 mL elutes using buffer B (25 mM HEPES pH 7.5, 500 mM NaCl, 5% Glycerol, 1 mM TCEP, 500 mM Imidazole). The protein was analyzed on a 15% SDS-PAGE gel.

#### Dialysis

Elutes from Hi-Trap IMAC column were pooled, added 3C Protease in the ratio of 1: 50 mg (protease: protein) and subjected to dialysis against 2 L of buffer C (25 mM HEPES pH 7.5, 500 mM NaCl, 5% Glycerol, 1 mM TCEP) overnight at 4 °C.

Protein, after overnight dialysis and cleavage of N-His tag, was passed through a pre-equilibrated (buffer A) 5 mL HiTrap IMAC FF Nickel column (GE Healthcare).

#### Size Exclusion Chromatography

The flow through from the above column was concentrated to 3 mL using a 10 kDa centrifugal concentrator (Sartorius Stedim) and loaded onto a pre-equilibrated (with buffer D: 25 mM HEPES pH 7.5, 500 mM NaCl, 5% Glycerol) 120 mL Superdex200 16/600 column (GE Healthcare). 2 mL fractions were collected and analyzed on a 15% SDS-PAGE gel. Fractions corresponding to pure TrmD protein were pooled and concentrated to 25 mg/mL, flash frozen in liquid nitrogen and stored at −80 °C. Identity of the purified protein was further confirmed by MALDI fingerprinting.

### Crystallization of apo form of full length *M. abscessus* TrmD

*M. abscessus* TrmD apo crystals were grown in 48-well sitting drop plates (Swiss CDI) in the following condition: 0.08 mM Sodium cacodylate pH 5.8 to 6.8, 1–2 M Ammonium sulfate. 24 mg/mL of the protein in storage buffer (25 mM HEPES pH 7.5, 500 mM NaCl, 5% Glycerol) at drop ratio 1 μL: 1 μL (protein: reservoir respectively) were set up and equilibrated against 70 μL reservoir.

### Soaking of TrmD native crystals with fragments and ligands

Crystals for this experiment were grown at 19 °C in 48-well sitting drop plates (Swiss CDI) in the following condition: 0.08 mM Sodium cacodylate pH 6.5 to 7.0, 1–2 M Ammonium sulfate, 20 mg/mL of the protein in storage buffer (25 mM HEPES pH 7.5, 500 mM NaCl, 5% Glycerol) at drop ratio 1 μL: 1 μL were set up and equilibrated against 250 μL reservoir. Further, the crystals were picked and allowed to soak in a 4 μL drop containing reservoir solution and 10 mM fragments/compound (in DMSO) which was then equilibrated against 700 μL of the corresponding reservoir solution overnight at 19 °C in 24-well hanging drop vapor diffusion set up.

### Co-crystallization of TrmD protein with SAM/ SAH/ AW6/ AW7

2-5 mM final concentration of compound in DMSO/water was added to 20 mg/mL of TrmD protein, mixed and incubated for 2 h on ice. Crystals were grown in the following condition: 0.08mM Sodium cacodylate pH 6.5 to 7.0, 1–2 M Ammonium sulfate or in sparse matrix screens: Wizard 1&2 (Molecular Dimensions), Wizard 3&4 (Molecular Dimensions), JCSG +Suite (Molecular Dimensions). The crystallization drops were set up at a protein to reservoir drop ratio of 0.3 μL: 0.3 μL, in 96-well (MRC2) sitting drop plate, using Mosquito crystallization robot (TTP labtech) and the drops were equilibrated against 70 μL of reservoir at 19 °C.

### X-ray Data Collection and Processing

The TrmD apo/ligand-bound crystals were cryo-cooled in mother liquor containing 27.5% ethylene glycol. X-ray data sets were collected on I04, I02, I03, I04-1 or I24 beamlines at the Diamond Light Source in the UK, using the rotation method at wavelength of 0.979 Å, Omega start: 0°, Omega Oscillation: 0.1-0.2°, Total oscillation: 210-240 ^o^, Total images: 2100-2400, Exposure time: 0.05-0.08 s. The diffraction images were processed using AutoPROC (Vonrhein et al., 2011), utilizing XDS (Kabsch, 2010) for indexing, integration, followed by POINTLESS (Evans, 2011), AIMLESS (Evans and Murshudov, 2013) and TRUNCATE (French, 1978) programs from CCP4 Suite (Winn et al., 2011) for data reduction, scaling and calculation of structure factor amplitudes and intensity statistics. All TrmD crystals belonged to space group P2_1_2_1_2_1_ and consisted of two protomers in the asymmetric unit.

### Structure Solution and refinement

The *Mycobacterium abscessus* TrmD Apo structure was solved by molecular replacement using PHASER (McCoy et al., 2007) with the atomic coordinates of *Mycobacterium abscessus* TrmD at 1.7 Å (PDB entry: 3QUV Seattle Structural Genomics Consortium for Infectious Diseases) as search model and TrmD ligand bound structures were solved by molecular replacement with the atomic coordinates of the solved *Mycobacterium abscessus* TrmD Apo structure (PDB entry: 6NVR) as search model. Structure refinement was carried out using REFMAC (Murshudov et al., 2011) and PHENIX (Adams et al., 2010).

The models obtained were manually re-built using COOT interactive graphics program (Emsley and Cowtan, 2004) and electron density maps were calculated with 2|F_o_|-|F_c_| and |F_o_| - |F_c_| coefficients. Positions of ligands and water molecules were located in difference electron density maps and OMIT difference maps |mFo − DFc| (Hodel, 1992) were calculated and analysed to further verify positions of fragments and ligands. The corresponding statistics and omit maps are presented in supplementary data **Table S1** and **Figure S8**.

### Differential scanning fluorimetry (DSF)

DSF were carried out in a 96-well format with each well containing 25 μL of reaction mixture of 10 μM TrmD protein in buffer (50 mM HEPES pH 7.5, 500 mM NaCl, 5% glycerol), 5 mM compound, 5% DMSO and 5X Sypro orange dye. Appropriate positive (Protein, DMSO and SAM) and negative (Protein, DMSO only) controls were also included. The measurements were performed in a Biorad-CFX connect thermal cycler using the following program: 25 °C for 10 mins followed by a linear increment of 0.5 °C every 30 sec to reach a final temperature of 95 °C. The results were analyzed using Microsoft excel.

### Isothermal Titration Calorimetry (ITC)

ITC experiments to quantify binding of ligands to TrmD were done as described in (Whitehouse *et al*. 2019) using Malvern MicroCal iTC200 or Auto-iTC200 systems at 25 °C. Titrations consisted of an initial injection (0.2 µL), discarded during data processing, followed by either 19 (2 µL) or 39 (1 µL) injections separated by intervals of 60 – 150 seconds duration. Protein was dialysed overnight at 4 °C in storage buffer (*M. abscessus* TrmD: 50 mM HEPES pH 7.5, 500 mM NaCl, 5% glycerol; *M. tuberculosis* TrmD: 25 mM HEPES pH 7.5, 500 mM NaCl). Sample cell and syringe solutions were prepared using the same storage buffer, with a final DMSO concentration of 2 – 10% according to ligand solubility in the buffer. TrmD concentrations of either 33 or 100 µM were used, with ligand to protein concentration ratios ranging from 10-20:1. Control titrations without protein were also performed and subtracted from ligand to protein titrations. Titrations were fitted with Origin software (OriginLab, Northampton, MA, USA), using a one-site binding model with N fixed to 1 only for weakly binding ligands. Titrations were typically performed once (n = 1), with multiple isotherms obtained (n > 1) for key compounds of interest. K_d_ values are reported to 2 significant figures. Error provided by Origin software due to model fit is reported when n = 1, whereas standard deviation is reported when n > 1.

### Biochemical activity assays

Assays for quantifying TrmD methylation reactions were carried out in 20 uL reactions consisting of 6.25 μM SAM, 0.1 μM TrmD and 6.25 μM tRNA in the presence of 0-500 μM compounds in serial dilutions using assay buffer containing 50 mM Tris-HCl pH 7.5, 10 mM MgCl_2_, 24 mM NH_4_Cl, 5% DMSO and 1 mM DTT in nuclease free water. tRNA sequences were identified from the *M. abscessus* genome sequence using tRNAscan-SE algorithm, (Lowe and Chan, 2016; Lowe and Eddy, 1997). The substrate tRNA^Pro^ for the assay was purchased commercially from Integrated DNA technologies (USA). The reactions were carried out for 1 h at room temperature followed by addition of 20 mM EDTA to stop the reactions. Each of the 20 uL samples were diluted ten-fold with the UPLC mobile phase solvent A (0.1% formic acid in water), centrifuged for 10 min at 13,000 g, to remove any precipitates, and the supernatant was aliquoted into 96-well plates. 40 uL samples were then injected into Acquity UPLC (Waters) T3 1.8 μM column and eluted using a gradient elution consisting of Mobile Phase A: 0.1% formic acid in water and mobile phase B: 0.1% formic acid in 100% methanol for 4 min. The absorbance was monitored using a photodiode array (PDA) detector (Waters) at wavelength range of ƛ: 220– 500 nm. All reactions were carried out in triplicate. The blank corrected data were analysed using Microsoft excel and non-linear regression analysis for IC_50_ determination were done using Graph Pad prism version 7.00, GraphPad Software, La Jolla California USA.

### Mycobacterial strains used and MIC measurements

The following mycobacterial strains were used: *Mycobacterium abscessus subspecies abscessus* (ATCC 19977) transformed with pmv310 plasmid expressing Lux ABDCE operon, grown in Middlebrook 7H9 broth supplemented with ADC (Sigma, UK) and *M. tuberculosis* ΔleuD ΔpanCD (Bleupan) (Sampson et al., 2004) transformed with pSMT1 expressing Lux AB and GFP, grown in Middlebrook 7H9 broth supplemented with 0.5% glycerol, 0.05% Tween 80 (removed for 24 h prior to experiments), 10% OADC (BD), 0.05 mg/ml L-leucine, and 0.024 mg/ml calcium pantothenate, Hygromycin and Zeocin (removed for 24 h prior to experiments). All the other NTM strains are clinical isolates. Minimum Inhibitory Concentrations (MIC) were determined for mycobacteria according to the Clinical and Laboratory Standards Institute (CLSI) method M07-A9. Briefly, mycobacteria were grown to optical density (A_600nm_) of 0.2-0.3 in liquid culture and 1×10^5^ bacteria were added to each well of 96-well plates containing serial dilutions of compound (400, 200, 100, 50, 25, 12.5, 6.3, 3.1, 1.6, 0.8, 0.4, 0 µM), in triplicate wells per condition, and incubated at 37 ^o^C until growth was seen in the control wells. MIC measurements using *M. tuberculosis* H37Rv were performed as reported in. *M. tuberculosis* H37Rv was grown in Middlebrook 7H9 base containing 14 mg/L dipalmitoyl phosphatidylcholine (DPPC), 0.81 g/L NaCl, 0.3 g/L casitone, and 0.05% Tyloxapol. H37Rv was grown and diluted to a similar inoculum size as mentioned above prior to exposure to serial dilutions of compounds (starting at 100 µM), and the plates were incubated at 37°C for two weeks. The MIC value was determined as the last well which showed no bacterial growth.

### Expression and purification of *M. tuberculosis* TrmD

A colony of *E. coli* strain ANG3685 (XL1 Blue pET23b-His6-trmDTB) kindly provided by the research group of Professor Angelika Gründling at Imperial College London (Zhang et al., 2017), was transferred to LB media (5 mL) with ampicillin (100 µg mL^−1^) and incubated overnight (37 °C, 160 rpm). The resultant material was processed with a GeneJET^TM^ Plasmid Miniprep Kit (Thermo Scientific™) to obtain plasmid (30 ng µL^−1^, A_260nm_/A_280nm_ 1.87), with identity confirmed by Sanger sequencing (DNA Sequencing Facility, Department of Biochemistry, University of Cambridge).

The isolated plasmid was used to transform *E. coli* strain BL21 (DE3), with a colony transferred to LB media (20 mL) with ampicillin (100 µg mL^−1^) and incubated overnight (37 °C, 160 rpm). The starter culture was used to inoculate 2 flasks, each containing LB media (1 L) with ampicillin (100 µg mL^−1^), with incubation (37 °C, 200 rpm) until an optical density (A_600nm_) of 0.5 was reached. Protein expression was induced by the addition of isopropyl β-D-1-thiogalactopyranoside (0.5 mM), followed by overnight incubation (20 °C, 200 rpm). Cells were harvested by centrifugation (4 °C, 4000 g, 20 minutes), then frozen.

The cells were resuspended in 50 mL lysis buffer (50 mM HEPES pH 7.4, 1 M NaCl, 25 mM imidazole, 5 mM mercaptoethanol) with a tablet of cOmplete™ Protease Inhibitor Cocktail (Roche). The suspension was sonicated (10 minutes: 10 seconds on/ 20 seconds off), centrifuged (4 °C, 30000 g, 20 minutes) and filtered (0.45 µm). The resultant lysate was loaded onto a 7.5 mL nickel Sepharose^TM^ fast flow column (GE Healthcare), pre-equilibrated with lysis buffer. The column was washed with 5 column volumes of buffer A (25 mM HEPES pH 7.5, 500 mM NaCl, 20 mM imidazole, 5 mM mercaptoethanol) and eluted with buffer B (25 mM HEPES pH 7.5, 500 mM NaCl, 500 mM imidazole, 5 mM mercaptoethanol) in 7×5 mL aliquots. Protein-containing aliquots, as determined by SDS-PAGE, were combined and concentrated (10 kDa cutoff) to a volume of 7 mL, then loaded onto a Superdex 75 Hiload^TM^ 16/60 column (GE Healthcare) pre-equilibrated with filtration buffer (25 mM HEPES pH 7.5, 500 mM NaCl, 5 mM mercaptoethanol). Protein-containing fractions, as determined by SDS-PAGE, were combined and concentrated (10 kDa cutoff) to 14.4 mg mL^−1^ (5.0 mg L^−1^ yield), then flash-frozen in liquid nitrogen and stored at −80 °C. The identity of the protein was confirmed by LCMS analysis.

### Macrophage infection study

Blood samples were donated by healthy volunteers who had undertaken informed consent in accordance with local Research Ethics Committee approval. Peripheral blood mononuclear cells were isolated from citrated peripheral blood samples by density gradient separation using Lympholyte (Cedarlane Labs), and subsequent CD14^+^ positive selection using the MACS Miltenyi Biotec Human CD14 microbead protocol (Miltenyi Biotec). CD14^+^ cells were differentiated in to macrophages using recombinant human granulocyte-macrophage colony-stimulating factor (200 ng/mL GM-CSF) and recombinant human interferon gamma (50 ng/ml IFNγ) (Peprotech) in standard tissue culture DMEM media containing fetal calf serum, penicillin and streptomycin. Following removal of antibiotics, macrophages were infected at a multiplicity of infection of 10:1 with *M abscessus* 19977 for 2 h, washed in sterile phosphate buffered saline, and then incubated in DMEM media with FCS and 25 µM of compound for 24 and 48 h. At the given time points, supernatant was saved for cell cytotoxicity studies, and *M abscessus* survival within the macrophages calculated by macrophage lysis in sterile water, and colony forming unit calculation on Columbia Blood Agar plates (VWR BDH).

### Cytotoxicity

Lactate dehydrogenase (LDH) was measured as a biomarker for cellular cytotoxicity using the Pierce LDH Cytotoxicity Assay Kit. Cell supernatant was measured at 2, 24 and 48 hours post infection according to the kit protocol.

### Nude mouse derived *M. leprae*

*Mycobacterium leprae* (isolate Thai-53) was maintained in serial passage in the foot pads of athymic nude mice (Envigo, US.). Mice were inoculated in the plantar surface of both hind feet with 5×10^7^ fresh, viable nude mice derived *M. leprae*. When the mouse foot pads became moderately enlarged (at ∼ 5 - 6 months), they were harvested for intracellular *M. leprae* as described previously (Truman and Krahenbuhl, 2001), washed by centrifugation, re-suspended in medium, enumerated by direct count of acid fast bacilli according to Shepard’s method (Shepard and McRae, 1968), held at 4 ^o^C pending quality control tests for contamination and viability (Truman and Krahenbuhl, 2001). Freshly harvested bacilli were always employed in experiments within 24 h of harvest.

### *M. leprae* axenic culture

Freshly harvested nude mouse foot pad derived *M. leprae* were suspended in modified 7H12 medium, **AW7** was added at different concentrations (100 μM – 6.25 μM) and were incubated for 7 days at 33 ^o^C. Media only and rifampin (Sigma, USA) at 2.4 μM were used as negative and positive controls. Following incubation aliquots of **AW7** treated and control *M. leprae* were processed for radiorespirometry (RR) as described previously (Lahiri et al., 2005).

### *M. leprae* macrophage culture

Bone marrow cells were obtained aseptically from both femurs of female BALB/c mice and cultured on plastic cover slips in Dulbecco modified Eagle media (DMEM, Life Technologies, USA) supplemented with 10% v/v fetal calf serum (Life Technologies), 25 mM/L HEPES (Sigma, USA), 2 mM/L glutamine (Sigma, USA), 50 μg/mL ampicillin (Sigma, USA) and 10 ng/mL of recombinant murine macrophages colony stimulating factor (R &D Systems, USA) for 6 - 7 days at 37°C and 5% CO_2_. The cells were infected with freshly harvested nude mice foot pad derived live *M. leprae* at a multiplicity of infection (MOI) of 20:1 overnight at 33 ^o^C and then washed to remove extracellular bacteria. **AW7** was added at different concentrations (100 μM – 6.25 μM) and the cells were incubated for 7 days at 33°C. Media only and rifampicin at 2.4 μM were used as negative and positive controls. **AW7** treated and control cells were lysed with sodium dodecyl sulfate (SDS, 0.1% w/v, Sigma, USA) and the intracellular *M. leprae* processed for radiorespirometry (Lahiri et al., 2010).

### Radiorespirometry

Metabolism of a suspension of *M. leprae* was measured by evaluating the oxidation of ^14^C-palmitic acid to ^14^CO_2_ by radiorespirometry as described previously (Franzblau, 1988). Levels of captured ^14^CO_2_ is proportional to the rate of ^14^C-palmitic acid oxidation and used as an indicator of *M. leprae* viability. In the present study the 7^th^ day cumulative counts per minute (CPM) were recorded and percentage inhibition of metabolism determined as compared to no drug control. Statistical significance between treatment groups and no drug control were determined by Student’s t-test and *P* <0.05 is considered as significant.

## Abbreviations

DSF: differential scanning fluorimetry
FBDD: fragment-based drug discovery
LE: ligand efficiency
MIC: minimum inhibitory concentration
NTM: nontuberculous mycobacteria
SAH: S-adenosyl-L-homocysteine
SAM: S-adenosyl-L-methionine

## Author Contributions

RAF, TLB, VM, AGC and CA conceived and managed the project. SET, AJW and VM wrote the manuscript and designed the experiments. SET and PG performed the molecular biology and expression, protein purification, characterization, crystallography and fragment library screening. AJW designed, synthesized and characterised the compounds. SET and AJW performed the biophysical and biochemical assays. KPB performed the microbiological experiments on *M. abscessus, M. tuberculosis Bluepan* and NTMs. MDJL and HIMB performed the microbiological experiments on *M. tuberculosis* H37Rv. MJ and JMB designed and carried out the *trmD* knockout studies. RL designed and performed the experiments on *M. leprae.* SM performed the bioinformatics studies and t-RNA sequence searches.

## Acknowledgements

This work is funded by the Cystic Fibrosis Trust (Registered as a charity in England and Wales (1079049) and in Scotland (SC040196)) Strategic research consortium and in part by the Intramural Research Program of the NIH, NIAID. The authors would like to thank the Diamond Light Source for beam-time (proposals mx9537, mx14043, mx18548) and the staff of beamlines I03, I02, I04, I04-1 and I24 for assistance with data collection, the Seattle Structural Genomics Consortium for kindly providing the *M. abscessus trmD* AVA0421 plasmid, Prof. Ben Luisi and Dr. Kasia J. Bandrya (Department of Biochemistry, University of Cambridge) for insights on experimental design using tRNA, Prof. Angelika Gründling (MRC Centre for Molecular Bacteriology and Infection, Imperial College London) for kindly providing an *E. coli* strain for the expression of *M. tuberculosis* TrmD, Dr. Sundeep Chaitanya (Department of Biochemistry, University of Cambridge) for discussions on *M. leprae* inhibition studies. SET is funded by the Cystic Fibrosis Trust (RG 70975) and Foundation Botnar (RG91317); VM is funded by the Bill and Melinda Gates Foundation SHORTEN-TB (RG 86546); TLB is funded by the Wellcome Trust (Wellcome Trust Investigator Award 200814_Z_16_Z: RG83114); AJW is funded through the EPSRC. RAF, TLB and CA would like to thank the Foundation Botnar for funding (Grant No. RG91317).

## Accession Numbers

Coordinates and structure factors related to this work been deposited in the PDB with accession numbers 6NVR, 6NW6, 6NW7, 6QO2, 6QO3,6QO4, 6QO6, 6QOA, 6QOC, 6QOD, 6QOE, 6QOF, 6QOG, 6QOH, 6QOI, 6QOJ, 6QOK, 6QOL, 6QOM, 6QON, 6QOO, 6QOP, 6QOQ, 6QOR, 6QOS, 6QOT, 6QOU, 6QOV, 6QOW, 6QOX, 6QQS, 6QQX, 6QQY, 6QR6, 6QR5 and 6QR8.

