## Supplemental data for "Fragment-based discovery of a new class of inhibitors targeting mycobacterial tRNA modification"

### Results

#### Conservation of catalytic residues in mycobacterial TrmD

A multiple sequence alignment of 5 mycobacterial TrmD enzyme sequences with 10 TrmD ortholog amino acid sequences (Figure S1) shows conservation of important catalytic residues and residues involved in tRNA recognition and methyl transfer reactions. The catalytic residues Asp169 and Arg154 are highly conserved across TrmD orthologs. Key residues such as Asp50, Gly59, His46 involved in tRNA G36 and G37 base recognition, interactions with bases at positions 38, 32 and anticodon branch of wild type tRNA respectively, are also broadly conserved. In mycobacterial TrmD, a Histidine replaces Phe171 which participates in recognition by the AdoMet methionine moiety initiating the catalytic cascade leading to methyl transfer reaction (Ito et al., 2015). Further, the C-terminal motif S-G-H/D-H involved in tRNA minor groove recognition is highly conserved across TrmD orthologs.

The amino-acid sequence of *M. abscessus* TrmD (MAB-3226c) was obtained from the National Centre for Biotechnology Information (NCBI) database. Orthologous proteins from *M.abscessus ; M. chelonae; M. fortuitumM.terrae, M.leprae; M.gordonae; M.tuberculosis; M.avium; Pseudomonas aeruginosa; Haemophilus influenzae; Escherichia coli* were identified by performing a protein-BLAST (Altschul et al., 1990; States and Gish, 1994) search with *M. abscessus* protein sequence. The sequences were aligned using Clustal Omega (Sievers et al., 2011) and analysed using Jalview (Waterhouse et al., 2009).

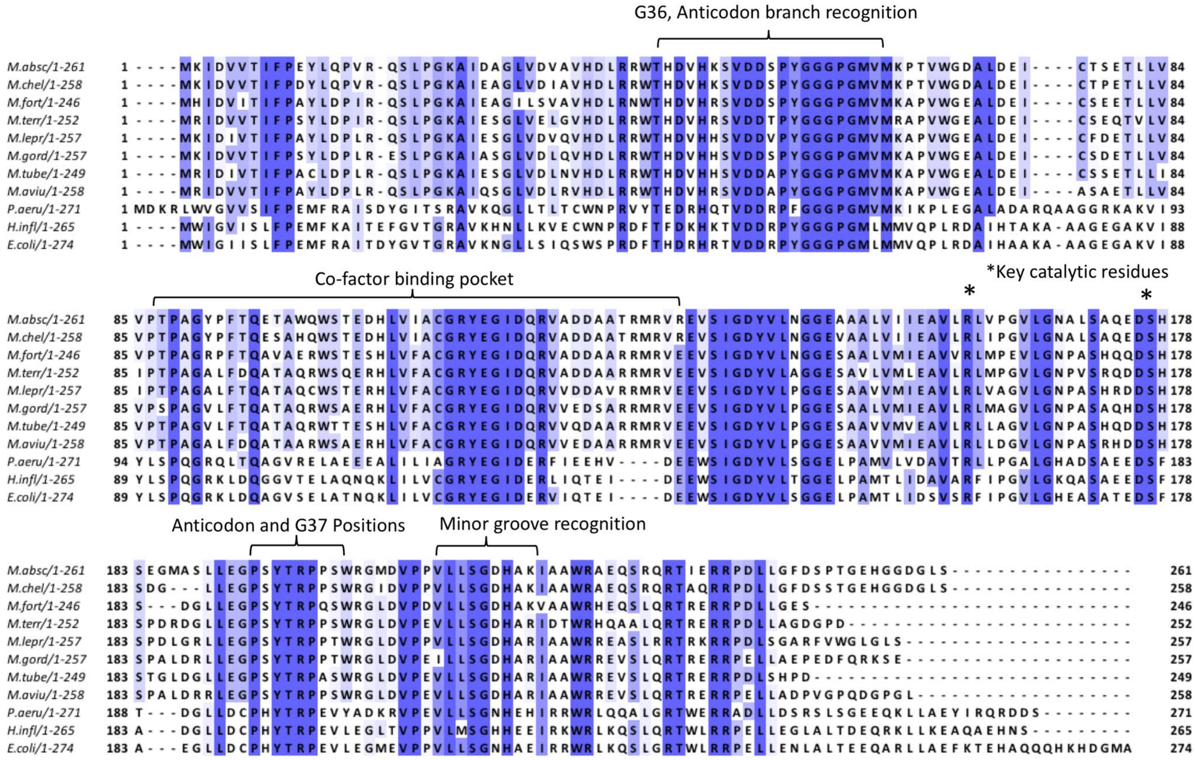


***Figure S1:*** *Multiple sequence alignment of 8 Mycobacterial TrmDs (M.absc- M.abscessus ;M.chel- M. chelonae; M.fort- M. fortuitum, M. terr- M.terrae, M.lepr- M.leprae; M. gord- M.gordonae; M.tube- M.tuberculosis; M.aviu- M.avium) with 3 other bacterial TrmD orthologs (P.aeru- Pseudomonas aeruginosa; H.influ- Haemophilus influenzae; E.coli- Escherichia coli) coloured from white to dark purple indicating increasing percentage conservation of amino acid residues. The important catalytic residues and their corresponding functions are also illustrated in the alignment.*

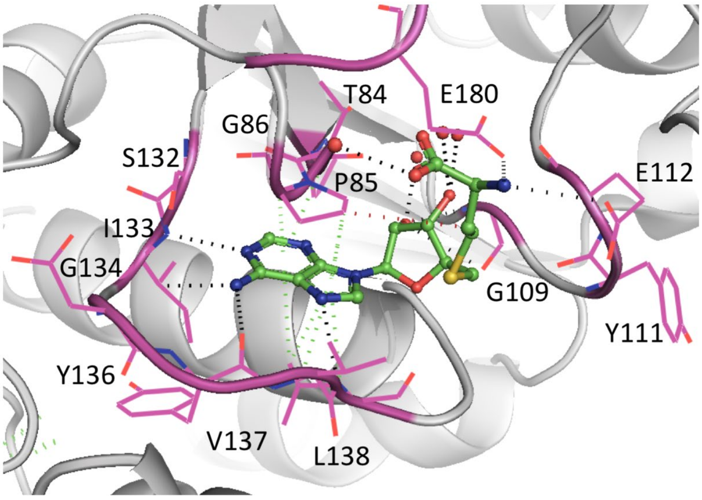


***Figure S2:*** *Interaction map of TrmD (pink) in complex S-adenosyl homocysteine (**SAH**), PDB code **6NW7**, shown as green stick representation. Hydrogen bonds, hydrophobic contacts, π-interactions are depicted in black, green and red dotted lines respectively.*

**X-ray crystal structures and clustering for all fragment hits reported in this work**

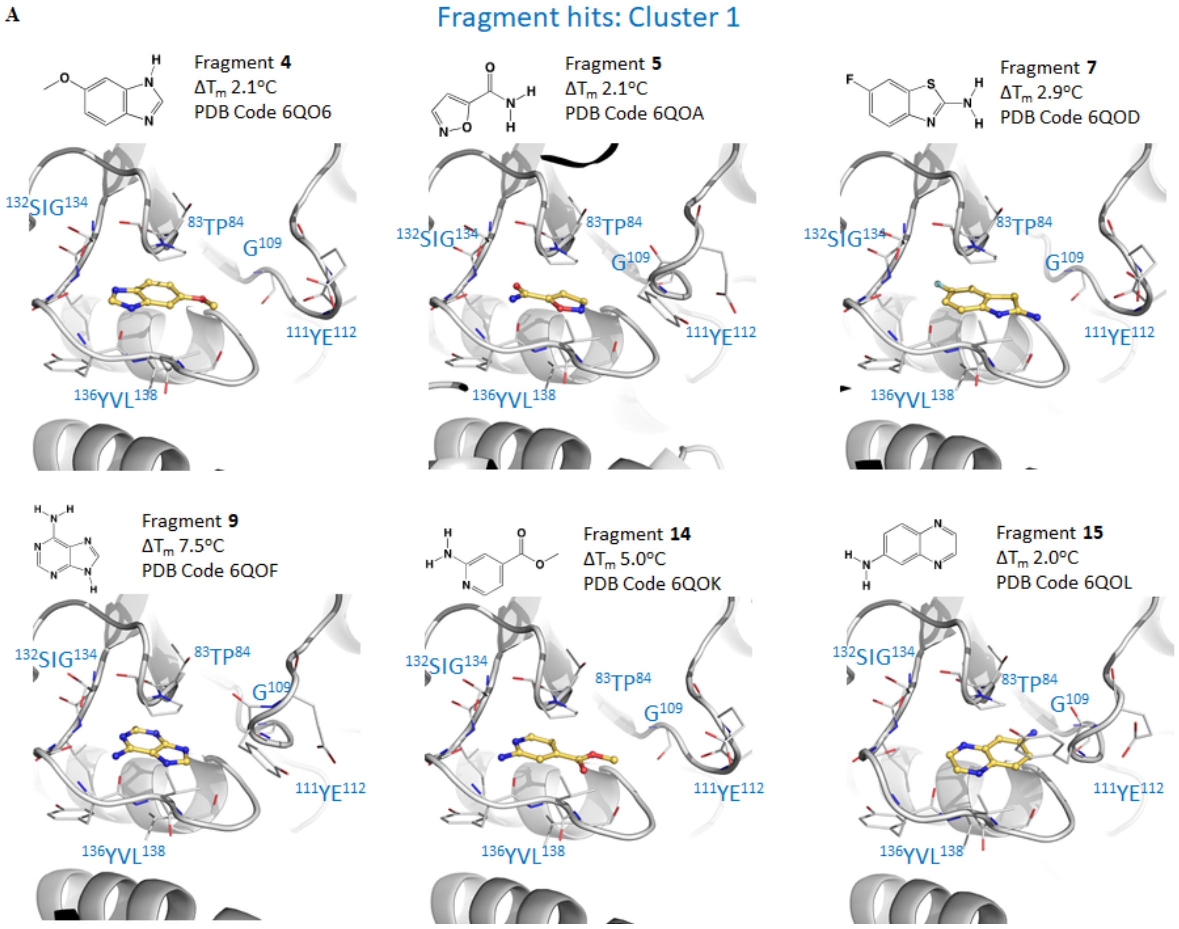

**B**

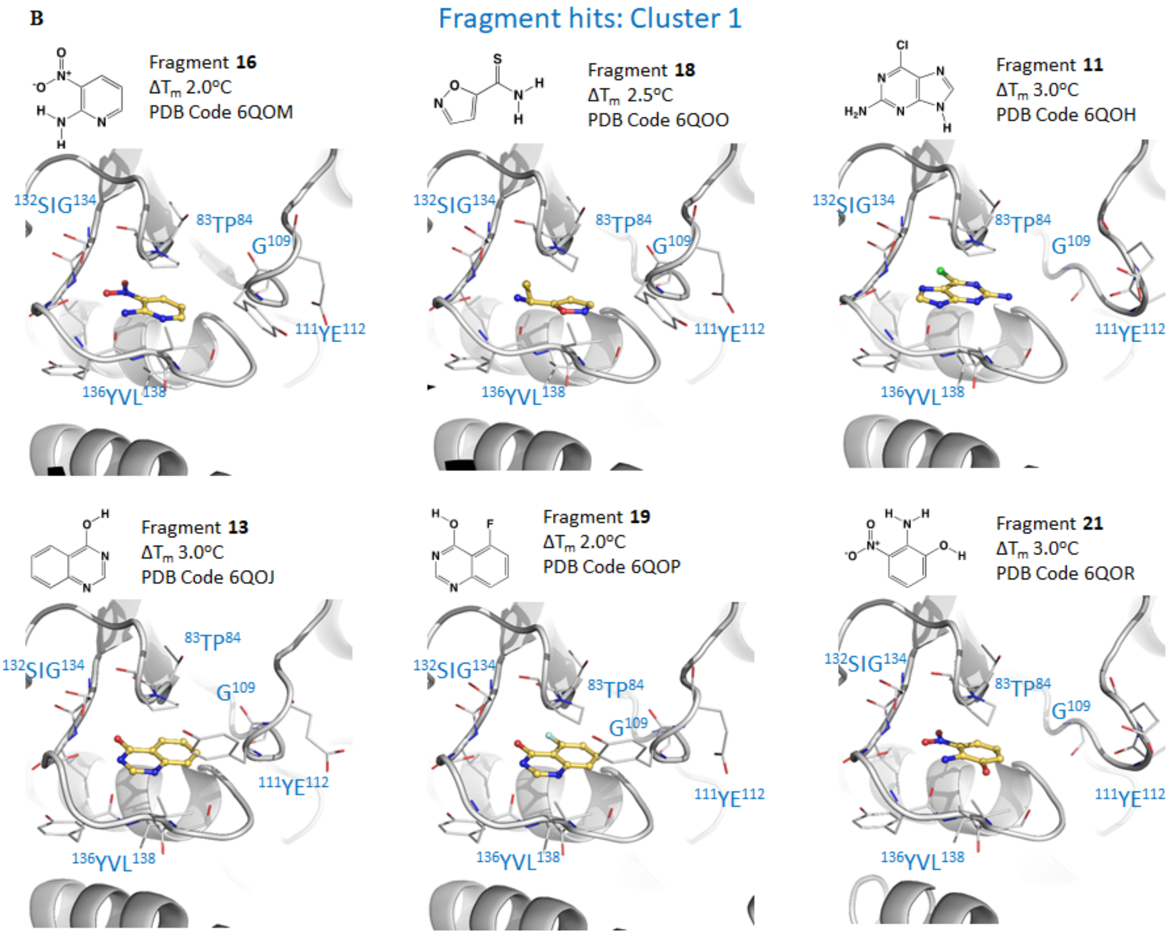

Fragment hits: Cluster 1
Fragment 16
ΔTm 2.0°C
PDB Code 6QOM
Fragment 18
ΔTm 2.5°C
PDB Code 6QOO
Fragment 11
ΔTm 3.0°C
PDB Code 6QOH
Fragment 13
ΔTm 3.0°C
PDB Code 6QOJ
Fragment 19
ΔTm 2.0°C
PDB Code 6QOP
Fragment 21
ΔTm 3.0°C
PDB Code 6QOR
132SIG134
83TP84
G109
111YE112
136YVL138

**C**

Fragment hits: Cluster 2

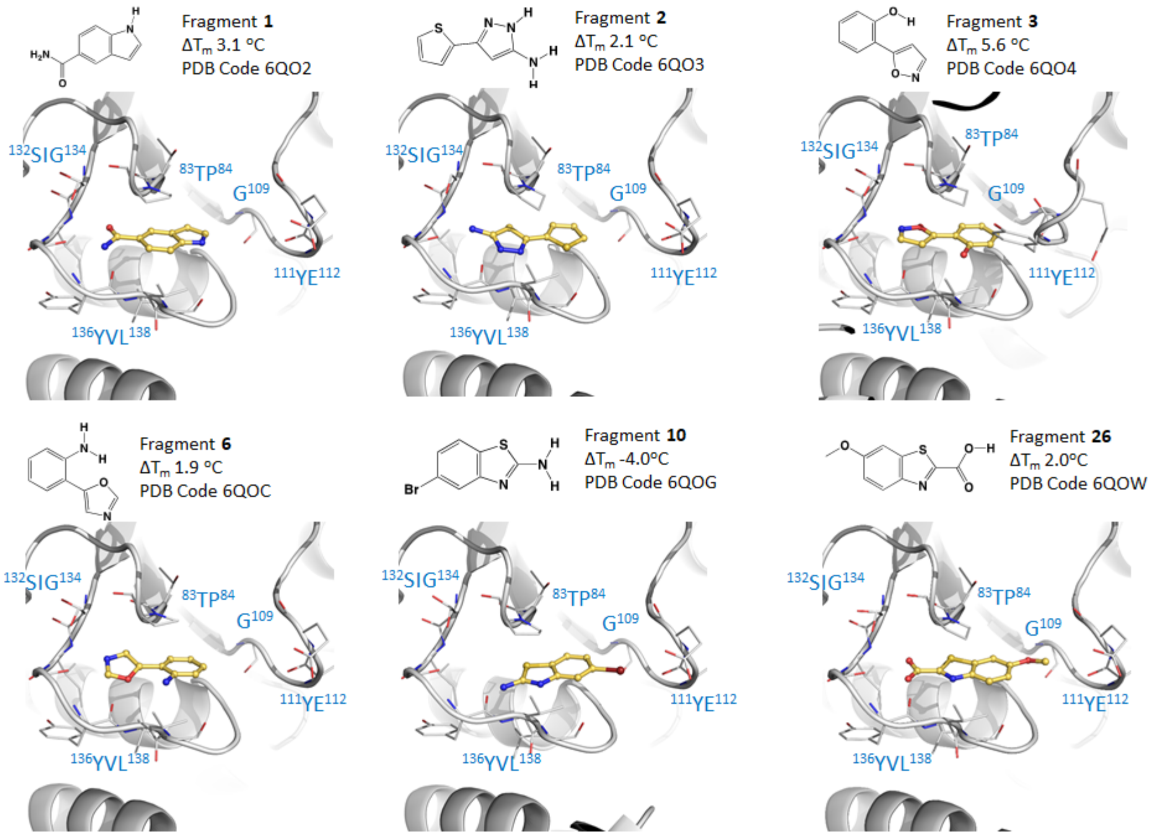


**D**

Fragment hits: Cluster 2

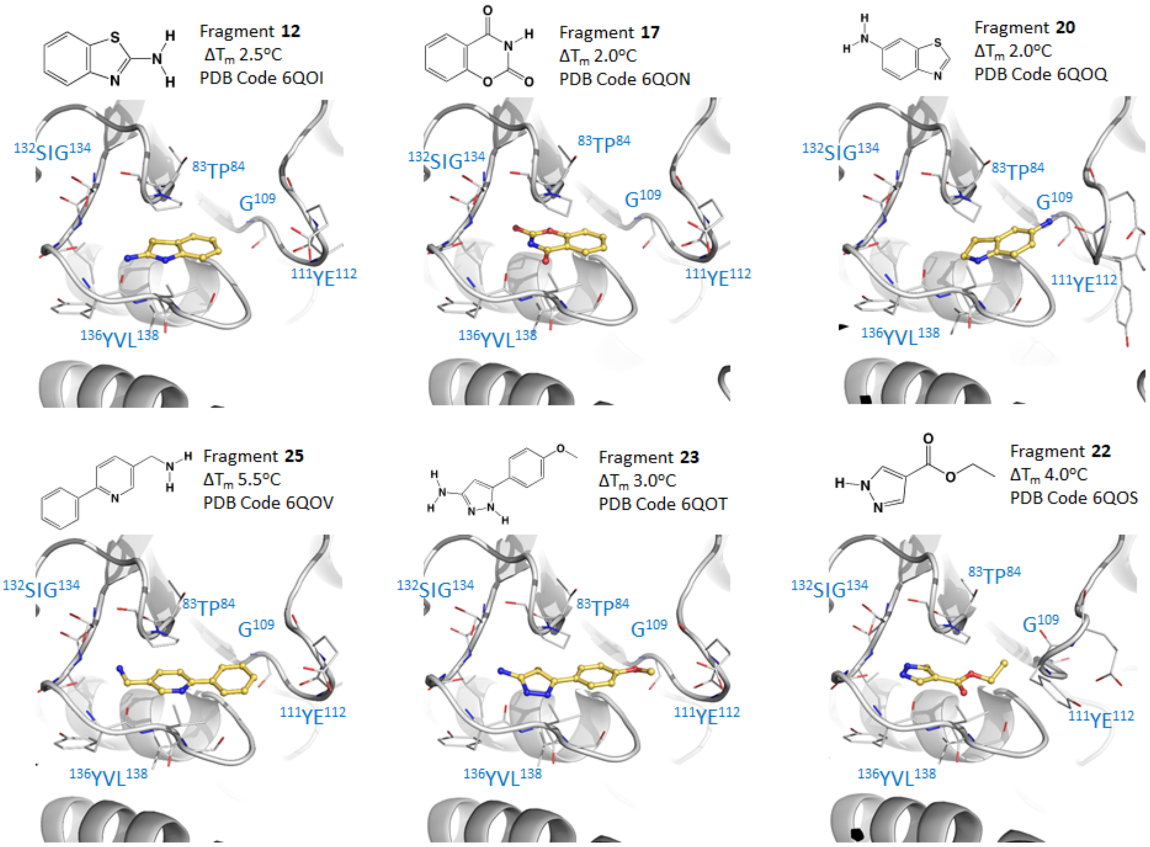

E

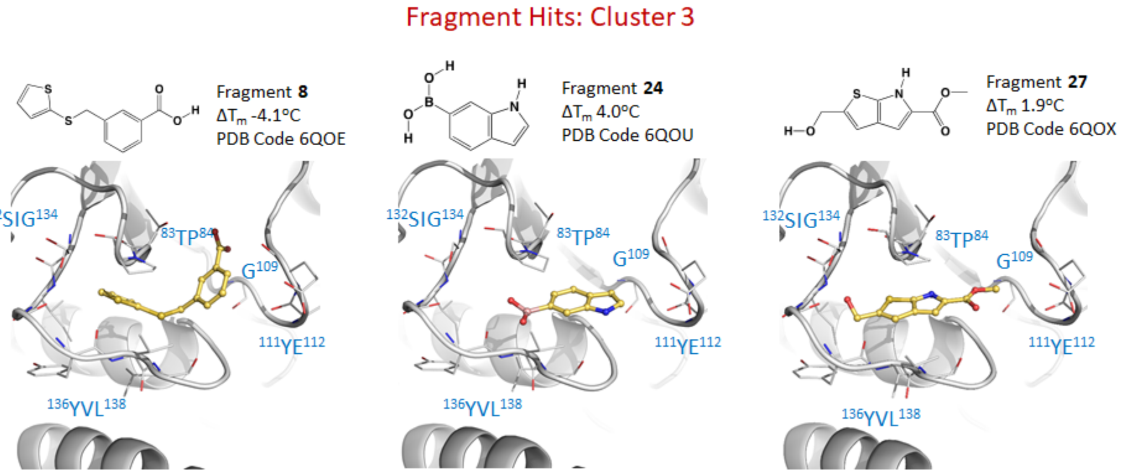


***Figure S3 A-E: Hits obtained from fragment library screening of TrmD***. *Fragment hits identified from DSF and X-ray crystallography, clustered into three groups based on the mode of binding at the M. abscessus TrmD active site.*

#### Enzymatic assays

A

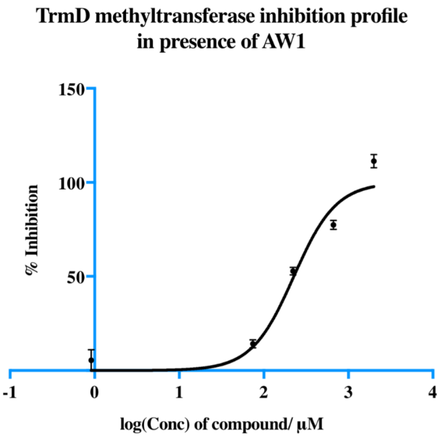


B

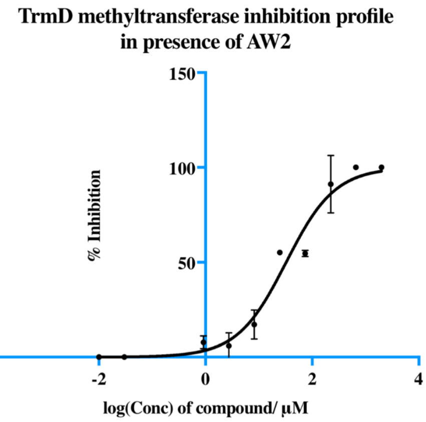

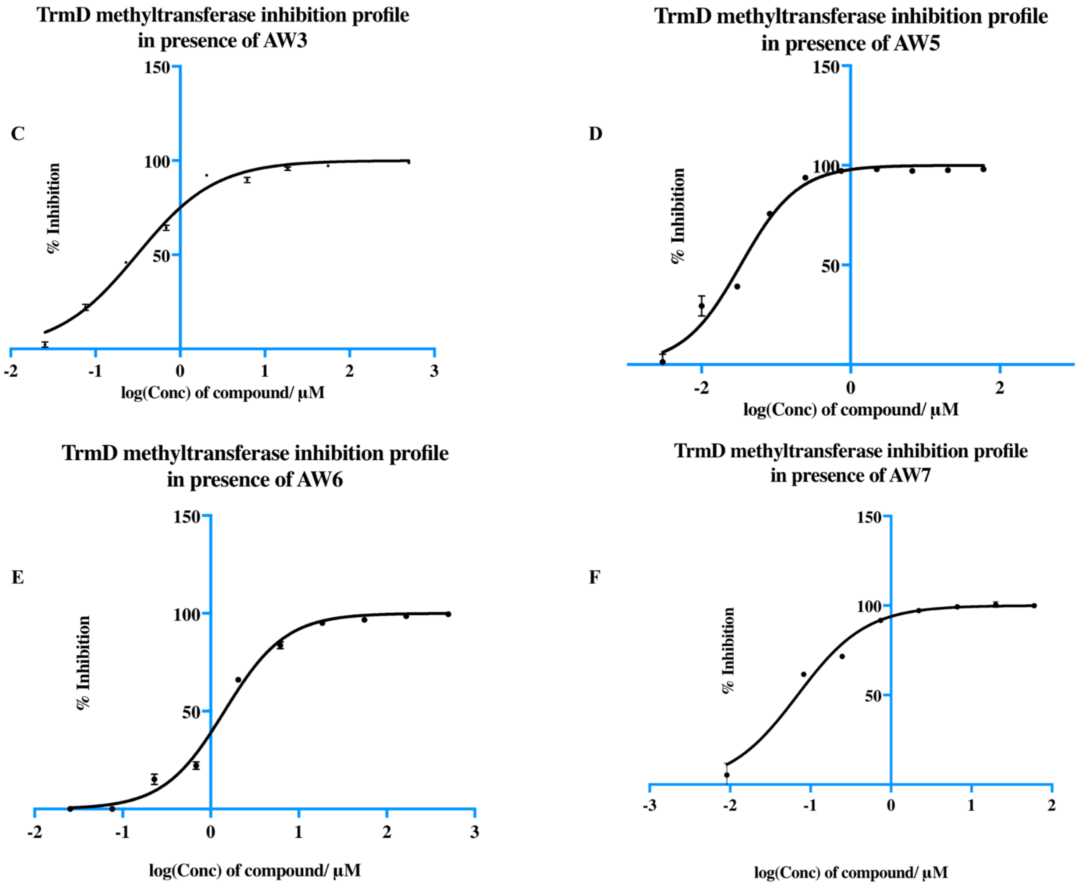


***Figure S4 A-F:*** *M. abscessus TrmD Methyltransferase inhibition profiles in the presence of TrmD lead compounds AW1-7.*

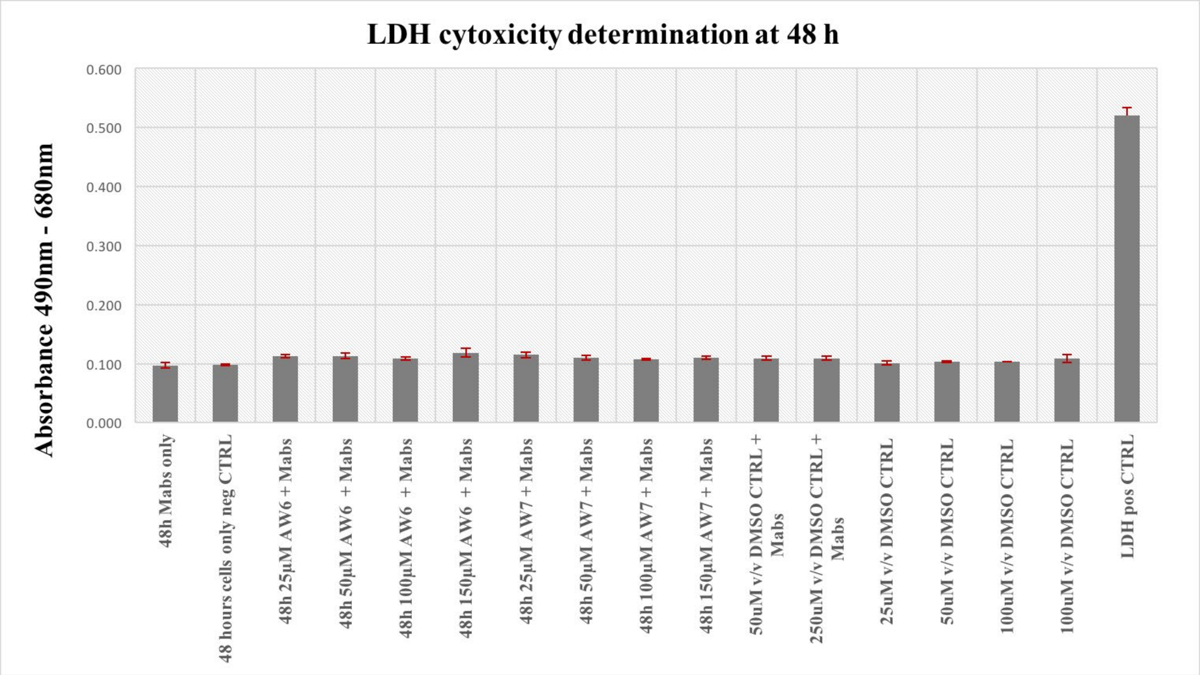


***Figure S5:*** *Lactate dehydrogenase* **(***LDH) based cytotoxicity profiles of TrmD lead compounds* ***AW6*** *&* ***AW7*** *in M. abscessus infected primary human macrophages.*

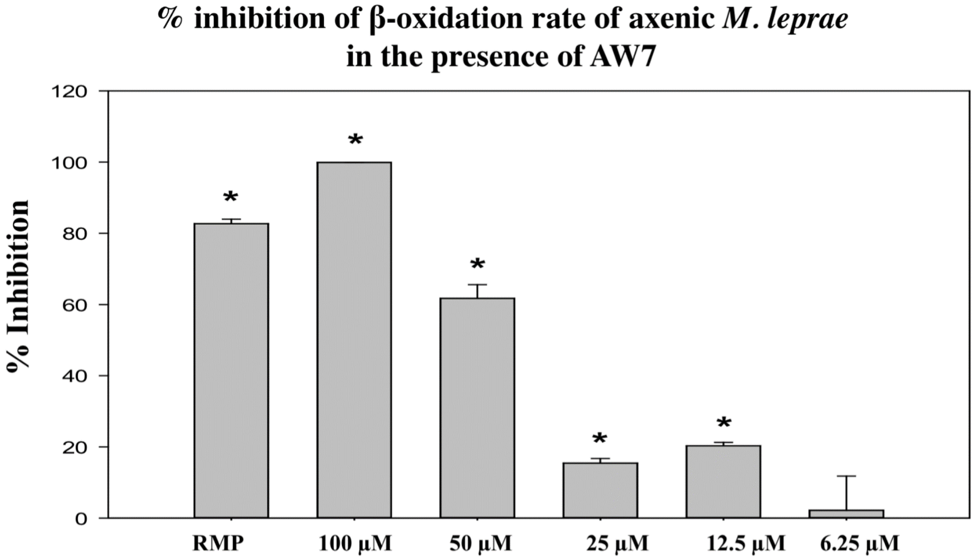


***Figure S6****: Percentage inhibition of M. leprae palmitic acid oxidation rate (radiorespirometry) in the presence of different concentrations of AW241 for 7 days. 7th day cumulative counts per minute (CPM) were recorded and percentage inhibition of metabolism determined as compared to no drug control. AW241 concentrations, in mM, are shown in parenthesis and rifampin (RMP) was used at 2mg/mL. * - Inhibition is statistically significant (P<0.05) compared to no drug control. Data representative of three separate experiments.*

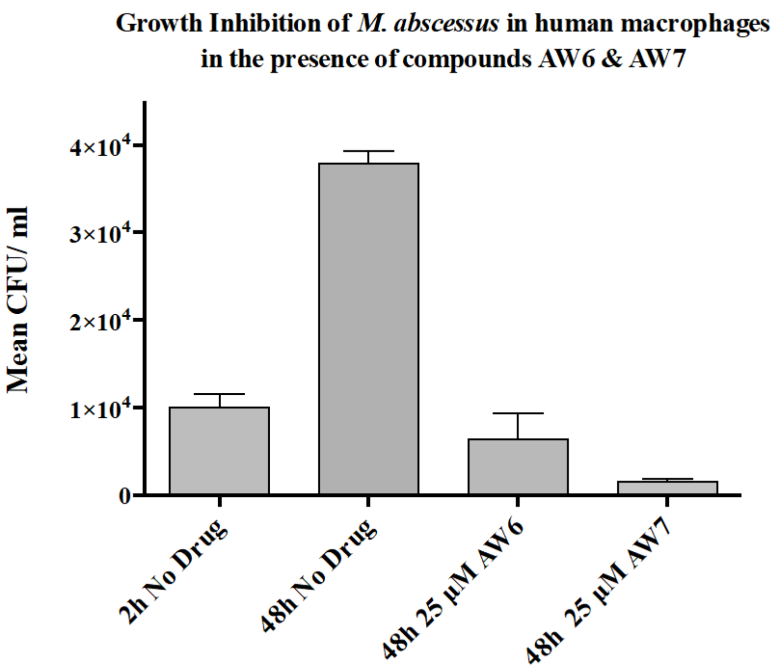


***Figure S7****: Growth inhibition study of lead compounds* ***AW6*** *&* ***AW7*** *in M. abscessus infected human macrophages over a 48 h period. Data representative of three separate experiments.*

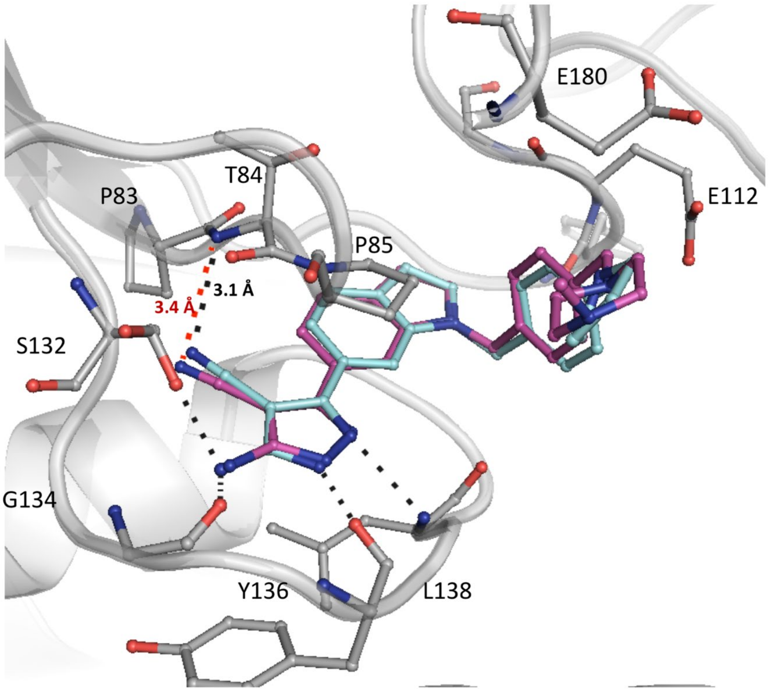


***Figure S8****: Superposition of the crystal structures of TrmD in complex with AW5 (blue stick) and AW7 (pink stick) showing changes in positon of the nitrile group leading to increased distance of 3.4 Å from Thr84 backbone amide there by diminishing H-bond interaction as compared to AW239 (3.1 Å) nitrile.*

*Table S1: X-ray Crystallographic Data Collection & Refinement Statistics for crystal structures described in this study*

| Ligand | Apo | S-adenosyl methionine (SAM) | S-adenosyl homocysteine (SAH) |
|---|---|---|---|
| **PDB Codes** | **6NVR** | **6NW6** | **6NW7** |
| **Resolution range (Å)** | 58.3 - 1.562 (1.618 - 1.562) | 56.91 - 1.667 (1.727 - 1.667) | 58.48 - 1.481 (1.534 - 1.481) |
| **Space group** | P 21 21 21 | P 21 21 21 | P 21 21 21 |
| **Unit cell** | 73.724 79.128 86.235 90 90 90 | 75.034 79.24 87.313 90 90 90 | 75.334 78.254 88.005 90 90 90 |
| **Total reflections** | 555563 (51454) | 476241 (48544) | 672385 (65190) |
| **Unique reflections** | 72163 (7142) | 61337 (6066) | 87015 (8620) |
| **Multiplicity** | 7.7 (7.2) | 7.8 (8.0) | 7.7 (7.6) |
| **Completeness (%)** | 99.98 (99.97) | 99.92 (99.95) | 99.91 (99.92) |
| **Mean I/sigma(I)** | 20.51 (2.30) | 16.52 (2.23) | 28.57 (2.33) |
| **Wilson B-factor** | 27.07 | 26.21 | 22.69 |
| **R-merge** | 0.04883 (0.7835) | 0.06756 (0.984) | 0.03225 (0.783) |
| **R-meas** | 0.05237 | 0.07266 | 0.03457 |
| **CC1/2** | 0.999 (0.801) | 0.998 (0.711) | 1 (0.782) |
| **CC*** | 1 (0.943) | 0.999 (0.911) | 1 (0.937) |
| **R-work** | 0.1927 (0.2480) | 0.1794 (0.2486) | 0.1876 (0.2505) |
| **R-free** | 0.2056 (0.2892) | 0.2033 (0.2799) | 0.2088 (0.2662) |
| **Number of non-hydrogen atoms** | 3692 | 3677 | 3675 |
| **macromolecules** | 3337 | 3371 | 3321 |
| **ligands** |  | 5 | 5 |
| **water** | 355 | 301 | 349 |
| **Protein residues** | 432 | 427 | 422 |
| **RMS(bonds)** | 0.006 | 0.012 | 0.008 |
| **RMS(angles)** | 1.04 | 1.22 | 1.11 |
| **Ramachandran favored (%)** | 99 | 98 | 99 |
| **Ramachandran outliers (%)** | 0.23 | 0.23 | 0 |
| **Clashscore** | 2.86 | 2.99 | 1.52 |
| **Average B-factor** | 33.7 | 34.3 | 29.7 |
| **macromolecules** | 32.9 | 33.3 | 28.8 |
| **ligands** |  | 57 | 32.2 |
| **solvent** | 41.3 | 44.6 | 38.4 |

| Ligand | Fragment 1 | Fragment 2 | Fragment 3 |
|---|---|---|---|
| **PDB Codes** | **6QO2** | **6QO3** | **6QO4** |
| **Resolution range (Å)** | 56.58 - 1.676 (1.736 - 1.676) | 56.27 - 1.645 (1.704 - 1.645) | 58.22 - 1.778 (1.841 - 1.778) |
| **Space group** | P 21 21 21 | P 21 21 21 | P 21 21 21 |
| **Unit cell** | 74.416 77.893 87.113 90 90 90 | 74.304 79.397 86.148 90 90 90 | 74.487 78.902 86.272 90 90 90 |
| **Total reflections** | 116261 (11471) | 124732 (12311) | 99055 (9736) |
| **Unique reflections** | 58320 (5759) | 62492 (6159) | 49601 (4879) |
| **Multiplicity** | 2.0 (2.0) | 2.0 (2.0) | 2.0 (2.0) |
| **Completeness (%)** | 99.20 (98.51) | 99.96 (99.98) | 99.99 (99.98) |
| **Mean I/sigma(I)** | 20.53 (2.19) | 14.66 (2.23) | 18.19 (2.19) |
| **Wilson B-factor** | 27.4 | 26.22 | 31.86 |
| **R-merge** | 0.01642 (0.3223) | 0.02281 (0.2933) | 0.01874 (0.353) |
| **R-meas** | 0.02322 | 0.03226 | 0.0265 |
| **CC1/2** | 1 (0.738) | 0.999 (0.795) | 0.999 (0.716) |
| **CC*** | 1 (0.922) | 1 (0.941) | 1 (0.913) |
| **R-work** | 0.1711 (0.2335) | 0.1745 (0.2533) | 0.1800 (0.2729) |
| **R-free** | 0.1981 (0.2688) | 0.2072 (0.3030) | 0.2134 (0.3163) |
| **Number of non-hydrogen atoms** | 3563 | 3614 | 3498 |
| **macromolecules** | 3264 | 3278 | 3248 |
| **ligands** | 24 | 22 | 24 |
| **water** | 275 | 314 | 226 |
| **Protein residues** | 427 | 429 | 424 |
| **RMS(bonds)** | 0.007 | 0.006 | 0.007 |
| **RMS(angles)** | 1 | 1.03 | 1.07 |
| **Ramachandran favored (%)** | 99 | 99 | 98 |
| **Ramachandran outliers (%)** | 0 | 0.24 | 0.24 |
| **Clashscore** | 1.24 | 0.77 | 3.57 |
| **Average B-factor** | 34.9 | 31.6 | 39.7 |
| **macromolecules** | 34.2 | 30.6 | 39.4 |
| **ligands** | 29.2 | 23.3 | 31.7 |
| **solvent** | 44.4 | 43 | 45.4 |

| Ligand | Fragment 4 | Fragment 5 | Fragment 6 |
|---|---|---|---|
| **PDB Codes** | **6QO6** | **6QOA** | **6QOC** |
| **Resolution range (Å)** | 56.22 - 1.745 (1.807 - 1.745) | 58.21 - 1.927 (1.996 - 1.927) | 45.82 - 1.838 (1.904 - 1.838) |
| **Space group** | P 21 21 21 | P 21 21 21 | P 21 21 21 |
| **Unit cell** | 74.077 78.772 86.324 90 90 90 | 73.763 80.272 84.542 90 90 90 | 74.205 78.712 86.601 90 90 90 |
| **Total reflections** | 103964 (10265) | 76955 (7619) | 89476 (8818) |
| **Unique reflections** | 52072 (5136) | 38594 (3817) | 44802 (4414) |
| **Multiplicity** | 2.0 (2.0) | 2.0 (2.0) | 2.0 (2.0) |
| **Completeness (%)** | 99.97 (99.96) | 99.93 (99.95) | 99.97 (100.00) |
| **Mean I/sigma(I)** | 19.41 (2.32) | 18.01 (2.42) | 20.68 (2.41) |
| **Wilson B-factor** | 31.55 | 36.97 | 35.4 |
| **R-merge** | 0.01638 (0.3367) | 0.01996 (0.3006) | 0.01579 (0.3125) |
| **R-meas** | 0.02316 | 0.02822 | 0.02233 |
| **CC1/2** | 0.999 (0.731) | 0.999 (0.827) | 0.999 (0.772) |
| **CC*** | 1 (0.919) | 1 (0.951) | 1 (0.934) |
| **R-work** | 0.1812 (0.2545) | 0.1749 (0.2488) | 0.1854 (0.2550) |
| **R-free** | 0.2088 (0.2988) | 0.2198 (0.3437) | 0.2096 (0.3125) |
| **Number of non-hydrogen atoms** | 3496 | 3477 | 3438 |
| **macromolecules** | 3223 | 3276 | 3229 |
| **ligands** | 22 | 16 | 24 |
| **water** | 251 | 185 | 185 |
| **Protein residues** | 422 | 426 | 420 |
| **RMS(bonds)** | 0.007 | 0.007 | 0.008 |
| **RMS(angles)** | 1.01 | 1.04 | 1.08 |
| **Ramachandran favored (%)** | 98 | 99 | 98 |
| **Ramachandran outliers (%)** | 0.72 | 0.23 | 1.2 |
| **Clashscore** | 2.19 | 2 | 2.66 |
| **Average B-factor** | 39.1 | 44.5 | 43 |
| **macromolecules** | 38.5 | 44.2 | 42.7 |
| **ligands** | 35.4 | 47.7 | 39.6 |
| **solvent** | 46.6 | 49 | 48.3 |

| Ligand | Fragment 7 | Fragment 8 | Fragment 9 |
|---|---|---|---|
| **PDB Codes** | **6QOD** | **6QOE** | **6QOF** |
| **Resolution range (Å)** | 56.3 - 1.852 (1.918 - 1.852) | 58.71 - 3.057 (3.166 - 3.057) | 45.51 - 1.756 (1.819 - 1.756) |
| **Space group** | P 21 21 21 | P 21 21 21 | P 21 21 21 |
| **Unit cell** | 74.143 78.379 86.54 90 90 90 | 74.596 79.739 86.758 90 90 90 | 73.688 80.749 82.971 90 90 90 |
| **Total reflections** | 87002 (8551) | 20438 (2020) | 99719 (9802) |
| **Unique reflections** | 43567 (4287) | 10229 (1011) | 50043 (4903) |
| **Multiplicity** | 2.0 (2.0) | 2.0 (2.0) | 2.0 (2.0) |
| **Completeness (%)** | 99.89 (99.81) | 99.71 (99.90) | 99.93 (100.00) |
| **Mean I/sigma(I)** | 16.39 (2.24) | 13.13 (2.60) | 13.29 (2.26) |
| **Wilson B-factor** | 29.65 | 57.07 | 28.61 |
| **R-merge** | 0.02446 (0.4081) | 0.04665 (0.2577) | 0.02784 (0.2886) |
| **R-meas** | 0.03459 | 0.06597 | 0.03937 |
| **CC1/2** | 0.999 (0.642) | 0.998 (0.909) | 0.999 (0.835) |
| **CC*** | 1 (0.884) | 0.999 (0.976) | 1 (0.954) |
| **R-work** | 0.1843 (0.2990) | 0.1891 (0.2725) | 0.1802 (0.2419) |
| **R-free** | 0.2116 (0.3414) | 0.2808 (0.4267) | 0.2135 (0.3003) |
| **Number of non-hydrogen atoms** | 3418 | 3249 | 3527 |
| **macromolecules** | 3211 | 3232 | 3292 |
| **ligands** | 22 | 16 | 28 |
| **water** | 185 | 1 | 207 |
| **Protein residues** | 422 | 424 | 429 |
| **RMS(bonds)** | 0.008 | 0.01 | 0.007 |
| **RMS(angles)** | 1.12 | 1.28 | 1.06 |
| **Ramachandran favored (%)** | 98 | 95 | 99 |
| **Ramachandran outliers (%)** | 0.24 | 0.24 | 0 |
| **Clashscore** | 2.05 | 5.79 | 1.52 |
| **Average B-factor** | 37.7 | 51.2 | 35.1 |
| **macromolecules** | 37.4 | 51.1 | 34.8 |
| **ligands** | 51.9 | 78.9 | 26.8 |
| **solvent** | 42.4 | 16.1 | 42.4 |

| Ligand | Fragment 10 | Fragment 11 | Fragment 12 |
|---|---|---|---|
| **PDB Codes** | **6QOG** | **6QOH** | **6QOI** |
| **Resolution range (Å)** | 53.75 - 1.547 (1.603 - 1.547) | 58.32 - 1.657 (1.717 - 1.657) | 45.78 - 1.857 (1.923 - 1.857) |
| **Space group** | P 21 21 21 | P 21 21 21 | P 21 21 21 |
| **Unit cell** | 73.75 78.513 86.707 90 90 90 | 73.652 79.108 86.318 90 90 90 | 74.107 79.509 85.468 90 90 90 |
| **Total reflections** | 147636 (14608) | 120782 (11932) | 86344 (8474) |
| **Unique reflections** | 73971 (7327) | 60509 (5971) | 43267 (4241) |
| **Multiplicity** | 2.0 (2.0) | 2.0 (2.0) | 2.0 (2.0) |
| **Completeness (%)** | 99.93 (99.92) | 99.96 (99.97) | 99.96 (100.00) |
| **Mean I/sigma(I)** | 21.56 (2.35) | 15.01 (2.24) | 17.45 (2.31) |
| **Wilson B-factor** | 24.52 | 26.86 | 36.31 |
| **R-merge** | 0.01437 (0.316) | 0.02302 (0.333) | 0.01907 (0.3199) |
| **R-meas** | 0.02032 | 0.03255 | 0.02696 |
| **CC1/2** | 1 (0.783) | 0.999 (0.775) | 0.999 (0.777) |
| **CC*** | 1 (0.937) | 1 (0.934) | 1 (0.935) |
| **R-work** | 0.1786 (0.2394) | 0.1889 (0.2515) | 0.1947 (0.2644) |
| **R-free** | 0.2010 (0.2474) | 0.2246 (0.2902) | 0.2251 (0.3098) |
| **Number of non-hydrogen atoms** | 3608 | 3528 | 3317 |
| **macromolecules** | 3279 | 3224 | 3152 |
| **ligands** | 11 | 27 | 20 |
| **water** | 318 | 277 | 145 |
| **Protein residues** | 428 | 419 | 417 |
| **RMS(bonds)** | 0.006 | 0.006 | 0.007 |
| **RMS(angles)** | 1.04 | 1.04 | 1.07 |
| **Ramachandran favored (%)** | 100 | 98 | 98 |
| **Ramachandran outliers (%)** | 0 | 0.72 | 0.73 |
| **Clashscore** | 0.92 | 3.44 | 2.09 |
| **Average B-factor** | 32.4 | 34.6 | 44.5 |
| **macromolecules** | 31.3 | 33.9 | 44.4 |
| **ligands** | 36.4 | 38.1 | 50.5 |
| **solvent** | 43 | 41.6 | 47.7 |

| Ligand | Fragment 13 | Fragment 14 | Fragment 15 |
|---|---|---|---|
| **PDB Codes** | **6QOJ** | **6QOK** | **6QOL** |
| **Resolution range (Å)** | 45.76 - 2.023 (2.095 - 2.023) | 58.18 - 1.476 (1.528 - 1.476) | 56.3 - 1.903 (1.97 - 1.903) |
| **Space group** | P 21 21 21 | P 21 21 21 | P 21 21 21 |
| **Unit cell** | 74.01 80.498 84.284 90 90 90 | 73.702 78.625 86.484 90 90 90 | 74.008 78.608 86.729 90 90 90 |
| **Total reflections** | 66991 (6581) | 169607 (16630) | 80642 (7953) |
| **Unique reflections** | 33536 (3293) | 85013 (8354) | 40390 (3982) |
| **Multiplicity** | 2.0 (2.0) | 2.0 (2.0) | 2.0 (2.0) |
| **Completeness (%)** | 99.90 (99.88) | 99.96 (99.89) | 99.98 (100.00) |
| **Mean I/sigma(I)** | 17.29 (2.17) | 21.66 (2.27) | 18.24 (2.38) |
| **Wilson B-factor** | 46.57 | 21.06 | 36.27 |
| **R-merge** | 0.01689 (0.368) | 0.01465 (0.3515) | 0.01889 (0.3324) |
| **R-meas** | 0.02389 | 0.02072 | 0.02671 |
| **CC1/2** | 1 (0.812) | 1 (0.72) | 1 (0.745) |
| **CC*** | 1 (0.947) | 1 (0.915) | 1 (0.924) |
| **R-work** | 0.1959 (0.3020) | 0.1700 (0.2838) | 0.1855 (0.2632) |
| **R-free** | 0.2366 (0.3695) | 0.1987 (0.3133) | 0.2151 (0.3458) |
| **Number of non-hydrogen atoms** | 3359 | 3689 | 3395 |
| **macromolecules** | 3270 | 3323 | 3236 |
| **ligands** | 22 | 32 | 22 |
| **water** | 67 | 334 | 137 |
| **Protein residues** | 429 | 431 | 422 |
| **RMS(bonds)** | 0.007 | 0.006 | 0.007 |
| **RMS(angles)** | 1.05 | 1.05 | 1.02 |
| **Ramachandran favored (%)** | 100 | 99 | 99 |
| **Ramachandran outliers (%)** | 0 | 0 | 0.24 |
| **Clashscore** | 2.32 | 1.21 | 2.66 |
| **Average B-factor** | 60.1 | 30.4 | 45 |
| **macromolecules** | 60.2 | 29.1 | 45 |
| **ligands** | 47.8 | 45.6 | 37.5 |
| **solvent** | 55.9 | 41.5 | 45.6 |

| Ligand | Fragment 16 | Fragment 17 | Fragment 18 |
|---|---|---|---|
| **PDB Codes** | **6QOM** | **6QON** | **6QOO** |
| **Resolution range (Å)** | 54.18 - 1.897 (1.965 - 1.897) | 56.33 - 1.81 (1.875 - 1.81) | 58.19 - 1.939 (2.008 - 1.939) |
| **Space group** | P 21 21 21 | P 21 21 21 | P 21 21 21 |
| **Unit cell** | 73.955 79.592 85.853 90 90 90 | 74.02 78.919 86.836 90 90 90 | 73.574 80.037 84.737 90 90 90 |
| **Total reflections** | 81353 (7977) | 93844 (9248) | 75366 (7415) |
| **Unique reflections** | 40737 (3993) | 47009 (4630) | 37761 (3709) |
| **Multiplicity** | 2.0 (2.0) | 2.0 (2.0) | 2.0 (2.0) |
| **Completeness (%)** | 99.90 (99.90) | 99.93 (99.96) | 99.95 (100.00) |
| **Mean I/sigma(I)** | 17.55 (2.39) | 17.12 (2.33) | 15.39 (2.19) |
| **Wilson B-factor** | 37.82 | 28.85 | 37.61 |
| **R-merge** | 0.01689 (0.34) | 0.02202 (0.3552) | 0.02256 (0.3031) |
| **R-meas** | 0.02389 | 0.03114 | 0.03191 |
| **CC1/2** | 1 (0.748) | 0.999 (0.708) | 0.999 (0.834) |
| **CC*** | 1 (0.925) | 1 (0.91) | 1 (0.954) |
| **R-work** | 0.1924 (0.2564) | 0.1745 (0.2703) | 0.1815 (0.2483) |
| **R-free** | 0.2296 (0.2935) | 0.2061 (0.3146) | 0.2263 (0.3209) |
| **Number of non-hydrogen atoms** | 3354 | 3587 | 3399 |
| **macromolecules** | 3212 | 3326 | 3245 |
| **ligands** | 10 | 29 | 21 |
| **water** | 132 | 232 | 133 |
| **Protein residues** | 419 | 431 | 424 |
| **RMS(bonds)** | 0.009 | 0.008 | 0.007 |
| **RMS(angles)** | 1.21 | 1.13 | 1.03 |
| **Ramachandran favored (%)** | 97 | 99 | 98 |
| **Ramachandran outliers (%)** | 1.2 | 0 | 0.24 |
| **Clashscore** | 2.37 | 2.41 | 1.86 |
| **Average B-factor** | 47 | 36 | 46.5 |
| **macromolecules** | 46.9 | 35.3 | 46.2 |
| **ligands** | 66.6 | 62.9 | 65.7 |
| **solvent** | 47.7 | 43.4 | 49.4 |

| Ligand | Fragment 19 | Fragment 20 | Fragment 21 |
|---|---|---|---|
| PDB Codes | 6QOP | 6QOQ | 6QOR |
| Resolution range (Å) | 58.21 - 1.912 (1.98 - 1.912) | 56.24 - 1.666 (1.726 - 1.666) | 45.75 - 1.671 (1.73 - 1.671) |
| Space group | P 21 21 21 | P 21 21 21 | P 21 21 21 |
| Unit cell | 74.133 78.896 86.245 90 90 90 | 74.048 78.489 86.472 90 90 90 | 74.915 77.088 87.278 90 90 90 |
| Total reflections | 79389 (7856) | 118831 (11748) | 118266 (11615) |
| Unique reflections | 39796 (3935) | 59496 (5884) | 59237 (5823) |
| Multiplicity | 2.0 (2.0) | 2.0 (2.0) | 2.0 (2.0) |
| Completeness (%) | 99.94 (99.97) | 99.97 (99.98) | 99.97 (99.98) |
| Mean I/sigma(I) | 16.88 (2.40) | 17.71 (2.30) | 16.47 (2.35) |
| Wilson B-factor | 38.79 | 27.16 | 25.31 |
| R-merge | 0.02026 (0.3011) | 0.01919 (0.3273) | 0.01969 (0.293) |
| R-meas | 0.02865 | 0.02714 | 0.02784 |
| CC1/2 | 0.999 (0.759) | 0.999 (0.722) | 0.999 (0.769) |
| CC* | 1 (0.929) | 1 (0.916) | 1 (0.932) |
| R-work | 0.1899 (0.2702) | 0.1723 (0.2409) | 0.1727 (0.2384) |
| R-free | 0.2227 (0.3457) | 0.1934 (0.2747) | 0.2064 (0.2593) |
| Number of non-hydrogen atoms | 3368 | 3573 | 3619 |
| macromolecules | 3217 | 3290 | 3263 |
| ligands | 29 | 20 | 27 |
| water | 122 | 263 | 329 |
| Protein residues | 419 | 426 | 421 |
| RMS(bonds) | 0.007 | 0.006 | 0.009 |
| RMS(angles) | 1.02 | 1.01 | 1.2 |
| Ramachandran favored (%) | 98 | 99 | 97 |
| Ramachandran outliers (%) | 0.72 | 0 | 0 |
| Clashscore | 1.72 | 1.99 | 2 |
| Average B-factor | 48.7 | 34.6 | 33.5 |
| macromolecules | 48.5 | 33.9 | 32.6 |
| ligands | 62.9 | 39.3 | 31.5 |
| solvent | 50.8 | 43.3 | 43 |

| Ligand | Fragment 22 | Fragment 23 | Fragment 24 |
|---|---|---|---|
| PDB Codes | 6QOS | 6QOT | 6QOU |
| Resolution range (Å) | 58.69 - 2.052 (2.126 - 2.052) | 57.06 - 1.618 (1.676 - 1.618) | 58.63 - 1.556 (1.611 - 1.556) |
| Space group | P 21 21 21 | P 21 21 21 | P 21 21 21 |
| Unit cell | 74.913 81.357 84.754 90 90 90 | 75.291 78.925 87.458 90 90 90 | 75.188 79.07 87.396 90 90 90 |
| Total reflections | 253108 (24437) | 133947 (13225) | 150513 (14876) |
| Unique reflections | 33037 (3253) | 67062 (6632) | 75337 (7445) |
| Multiplicity | 7.7 (7.5) | 2.0 (2.0) | 2.0 (2.0) |
| Completeness (%) | 99.89 (99.94) | 99.91 (99.83) | 99.95 (99.95) |
| Mean I/sigma(I) | 15.59 (2.41) | 26.32 (2.33) | 31.53 (2.13) |
| Wilson B-factor | 39.66 | 26.25 | 25.28 |
| R-merge | 0.06667 (0.7334) | 0.01205 (0.3141) | 0.009729 (0.3395) |
| R-meas | 0.07153 | 0.01704 | 0.01376 |
| CC1/2 | 0.999 (0.857) | 1 (0.784) | 1 (0.753) |
| CC* | 1 (0.961) | 1 (0.937) | 1 (0.927) |
| R-work | 0.1911 (0.2634) | 0.1831 (0.2301) | 0.1835 (0.2416) |
| R-free | 0.2289 (0.2944) | 0.2143 (0.2640) | 0.2060 (0.2643) |
| Number of non-hydrogen atoms | 3432 | 3538 | 3574 |
| macromolecules | 3267 | 3245 | 3240 |
| ligands | 25 | 32 | 24 |
| water | 140 | 261 | 310 |
| Protein residues | 427 | 424 | 422 |
| RMS(bonds) | 0.008 | 0.009 | 0.009 |
| RMS(angles) | 1.03 | 1.03 | 1.04 |
| Ramachandran favored (%) | 99 | 99 | 98 |
| Ramachandran outliers (%) | 0 | 0 | 0 |
| Clashscore | 1.85 | 0.62 | 1.4 |
| Average B-factor | 50 | 33.8 | 33.8 |
| macromolecules | 49.9 | 33.1 | 32.9 |
| ligands | 60.2 | 25.5 | 27.1 |
| solvent | 50.2 | 43.4 | 44.3 |

| Ligand | Fragment 25 | Fragment 26 | Fragment 27 |
|---|---|---|---|
| **PDB Codes** | **6QOV** | **6QOW** | **6QOX** |
| **Resolution range (Å)** | 58.68 - 1.548 (1.603 - 1.548) | 58.48 - 1.525 (1.58 - 1.525) | 58.7 - 1.739 (1.801 - 1.739) |
| **Space group** | P 21 21 21 | P 21 21 21 | P 21 21 21 |
| **Unit cell** | 75.033 79.041 87.583 90 90 90 | 75.693 78.16 88.152 90 90 90 | 74.963 78.46 88.483 90 90 90 |
| **Total reflections** | 152376 (15078) | 159811 (15805) | 108501 (10712) |
| **Unique reflections** | 76391 (7557) | 80070 (7918) | 54302 (5361) |
| **Multiplicity** | 2.0 (2.0) | 2.0 (2.0) | 2.0 (2.0) |
| **Completeness (%)** | 99.86 (99.88) | 99.81 (99.60) | 99.96 (100.00) |
| **Mean I/sigma(I)** | 25.49 (2.23) | 25.95 (2.36) | 24.28 (2.26) |
| **Wilson B-factor** | 22.28 | 22.17 | 29.04 |
| **R-merge** | 0.01377 (0.3227) | 0.01348 (0.3093) | 0.01492 (0.3145) |
| **R-meas** | 0.01947 | 0.01906 | 0.02109 |
| **CC1/2** | 1 (0.778) | 1 (0.782) | 1 (0.778) |
| **CC*** | 1 (0.935) | 1 (0.937) | 1 (0.935) |
| **R-work** | 0.1822 (0.2402) | 0.1757 (0.2441) | 0.1841 (0.2644) |
| **R-free** | 0.2066 (0.2733) | 0.1968 (0.2812) | 0.2135 (0.3040) |
| **Number of non-hydrogen atoms** | 3544 | 3685 | 3563 |
| **macromolecules** | 3230 | 3269 | 3262 |
| **ligands** | 28 | 33 | 14 |
| **water** | 286 | 383 | 287 |
| **Protein residues** | 419 | 422 | 425 |
| **RMS(bonds)** | 0.009 | 0.011 | 0.009 |
| **RMS(angles)** | 1.02 | 1.19 | 1.02 |
| **Ramachandran favored (%)** | 98 | 98 | 98 |
| **Ramachandran outliers (%)** | 0 | 0 | 0 |
| **Clashscore** | 1.25 | 2.61 | 0.93 |
| **Average B-factor** | 30.5 | 30.3 | 36.9 |
| **macromolecules** | 29.7 | 29.1 | 36.2 |
| **ligands** | 21 | 30.1 | 44 |
| **solvent** | 40.3 | 40.4 | 45.2 |

| Ligand | AW1 | AW2 | AW3 |
|---|---|---|---|
| **PDB Codes** | **6QQS** | **6QQX** | **6QQY** |
| **Resolution range (Å)** | 54.15 - 1.758 (1.821 - 1.758) | 56.49 - 2.693 (2.789 - 2.693) | 54.03 - 1.49 (1.543 - 1.49) |
| **Space group** | P 21 21 21 | P 21 21 21 | P 21 21 21 |
| **Unit cell** | 74.357 79.028 85.537 90 90 90 | 74.614 78.22 86.471 90 90 90 | 74.51 78.45 86.04 90 90 90 |
| **Total reflections** | 386872 (38846) | 125554 (13048) | 165480 (16281) |
| **Unique reflections** | 50846 (5047) | 14535 (1440) | 82869 (8156) |
| **Multiplicity** | 7.6 (7.7) | 8.6 (9.1) | 2.0 (2.0) |
| **Completeness (%)** | 99.98 (100.00) | 99.92 (99.93) | 99.95 (99.96) |
| **Mean I/sigma(I)** | 18.44 (2.21) | 8.33 (2.19) | 22.56 (1.42) |
| **Wilson B-factor** | 30.86 | 36.88 | 24.83 |
| **R-merge** | 0.05835 (1.035) | 0.2334 (1.087) | 0.01375 (0.5068) |
| **R-meas** | 0.06277 | 0.2489 | 0.01945 |
| **CC1/2** | 0.999 (0.716) | 0.984 (0.77) | 1 (0.59) |
| **CC*** | 1 (0.914) | 0.996 (0.933) | 1 (0.861) |
| **R-work** | 0.1829 (0.2675) | 0.1778 (0.2386) | 0.1853 (0.2846) |
| **R-free** | 0.2081 (0.2996) | 0.2698 (0.3798) | 0.1997 (0.3104) |
| **Number of non-hydrogen atoms** | 3534 | 3372 | 3701 |
| **macromolecules** | 3282 | 3263 | 3275 |
| **ligands** | 30 | 49 | 48 |
| **water** | 222 | 60 | 378 |
| **Protein residues** | 423 | 422 | 422 |
| **RMS(bonds)** | 0.007 | 0.01 | 0.008 |
| **RMS(angles)** | 1.13 | 1.16 | 1.11 |
| **Ramachandran favored (%)** | 99 | 97 | 99 |
| **Ramachandran outliers (%)** | 0.24 | 0.24 | 0 |
| **Clashscore** | 3.21 | 3.69 | 4.13 |
| **Average B-factor** | 39.9 | 34 | 31.4 |
| **macromolecules** | 39.5 | 34.1 | 30.4 |
| **ligands** | 27.2 | 25.7 | 24 |
| **solvent** | 47.1 | 32.8 | 41 |

| Ligand | AW5 | AW6 | AW7 |
|---|---|---|---|
| **PDB Codes** | **6QR6** | **6QR5** | **6QR8** |
| **Resolution range (Å)** | 45.85 - 1.707 (1.768 - 1.707) | 56.27 - 1.81 (1.875 - 1.81) | 57.2 - 2.149 (2.225 - 2.149) |
| **Space group** | P 21 21 21 | P 21 21 21 | P 21 21 21 |
| **Unit cell** | 74.95 77.483 87.347 90 90 90 | 74.183 77.985 86.363 90 90 90 | 74.85 76.329 86.381 90 90 90 |
| **Total reflections** | 484104 (49526) | 92506 (9074) | 260137 (26060) |
| **Unique reflections** | 55948 (5512) | 46297 (4542) | 27601 (2722) |
| **Multiplicity** | 8.7 (9.0) | 2.0 (2.0) | 9.4 (9.6) |
| **Completeness (%)** | 99.97 (100.00) | 99.96 (100.00) | 99.94 (100.00) |
| **Mean I/sigma(I)** | 20.79 (2.35) | 26.01 (2.49) | 10.58 (2.37) |
| **Wilson B-factor** | 31.78 | 35.93 | 40.03 |
| **R-merge** | 0.0484 (0.8462) | 0.01154 (0.2436) | 0.1318 (1.18) |
| **R-meas** | 0.05156 | 0.01632 | 0.1395 |
| **CC1/2** | 0.999 (0.781) | 1 (0.873) | 0.998 (0.727) |
| **CC*** | 1 (0.936) | 1 (0.966) | 0.999 (0.918) |
| **R-work** | 0.1807 (0.2398) | 0.1922 (0.2614) | 0.1848 (0.2478) |
| **R-free** | 0.2014 (0.2654) | 0.2123 (0.3004) | 0.2262 (0.2750) |
| **Number of non-hydrogen atoms** | 3559 | 3504 | 3408 |
| **macromolecules** | 3277 | 3248 | 3245 |
| **ligands** | 65 | 56 | 64 |
| **water** | 217 | 200 | 99 |
| **Protein residues** | 422 | 424 | 423 |
| **RMS(bonds)** | 0.008 | 0.007 | 0.009 |
| **RMS(angles)** | 1.08 | 1.23 | 1.13 |
| **Ramachandran favored (%)** | 99 | 99 | 99 |
| **Ramachandran outliers (%)** | 0 | 0 | 0 |
| **Clashscore** | 2.44 | 2.78 | 2 |
| **Average B-factor** | 37.6 | 41 | 46.7 |
| **macromolecules** | 37.2 | 40.7 | 46.6 |
| **ligands** | 35.2 | 39.1 | 45.7 |
| **solvent** | 44.2 | 45.4 | 48.4 |

**Difference electron density maps (omit maps) for ligands illustrated in this study**

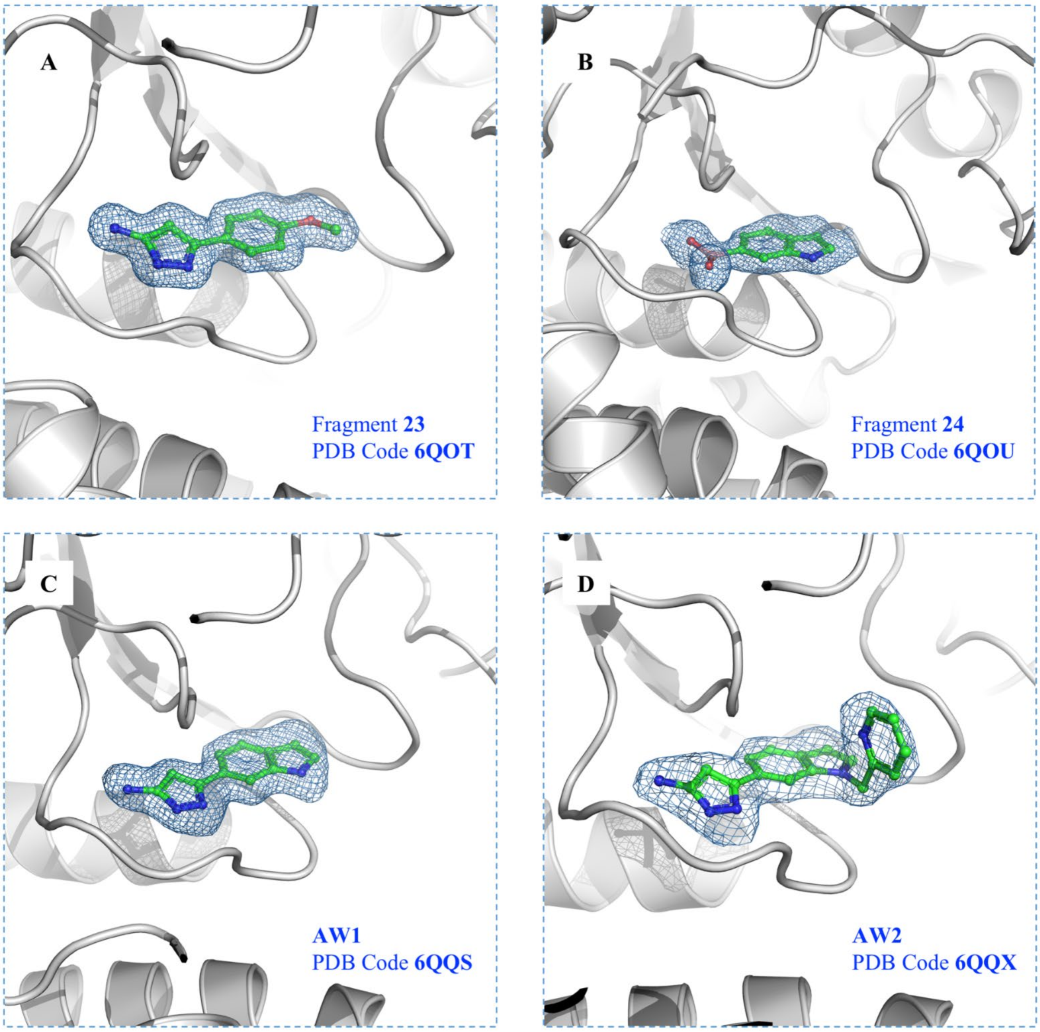

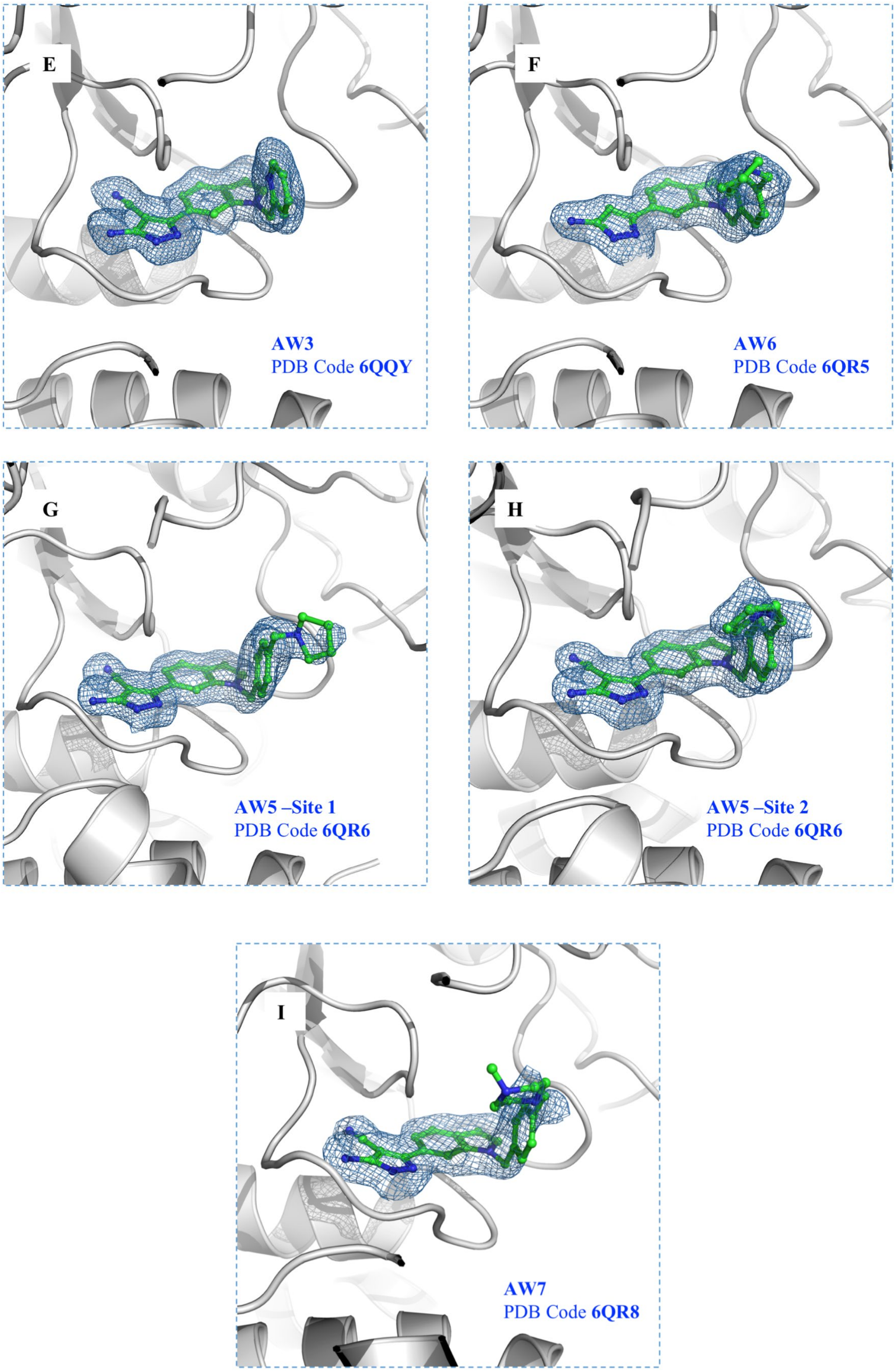


***Figure S9***: *Sigma A weighted Fo-Fc Omit maps corresponding to fragments and compounds described in this study*

### Isothermal Titration Calorimetry (ITC)

***Figure S10:*** *ITC traces with TrmD from Mycobacterium abscessus for a) fragment 23 (n = 1), b) fragment 24 (n = 1), c) AW1 (n = 2), and d) AW2 (n = 1).*

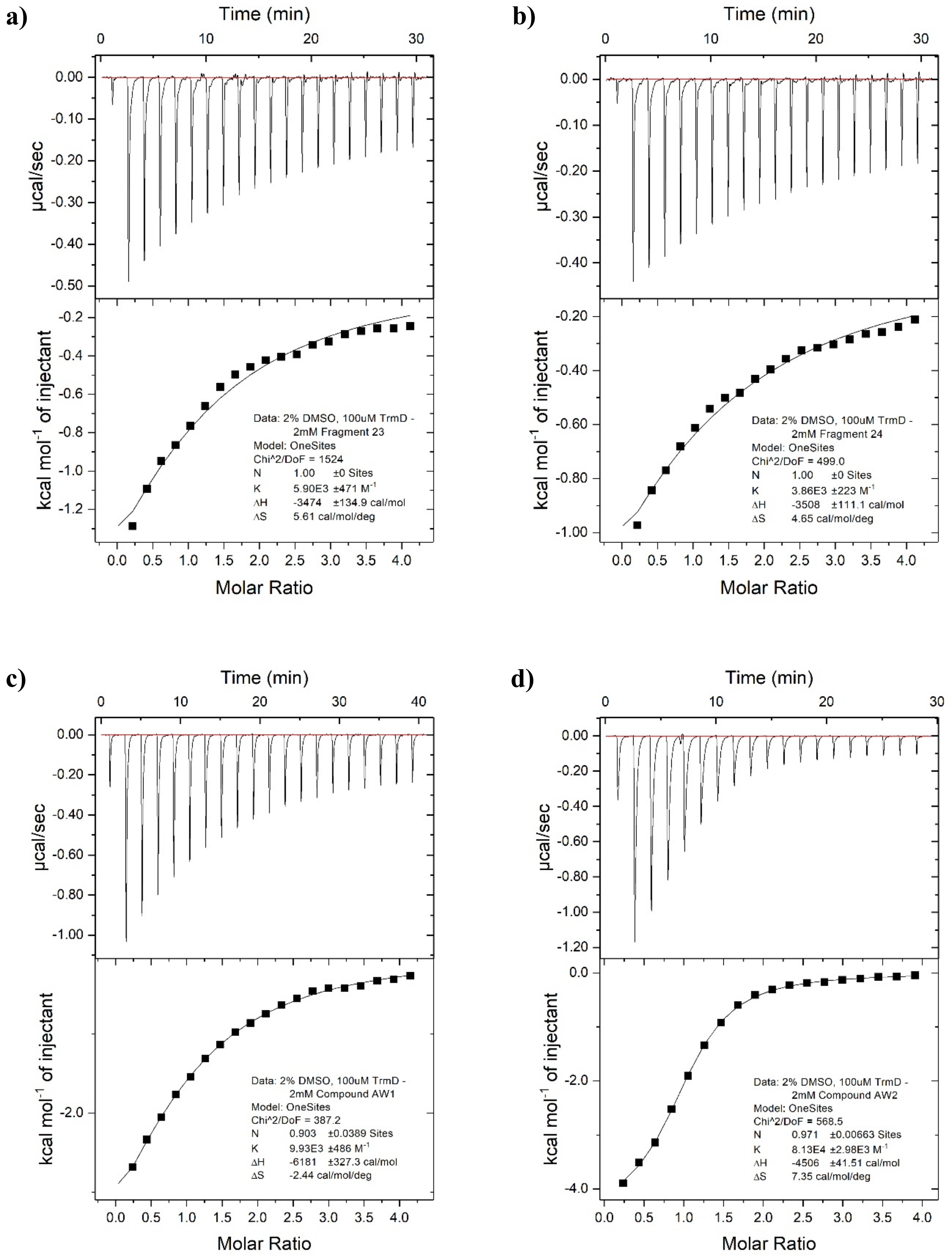

***Figure S11:*** *ITC traces with TrmD from Mycobacterium abscessus for a) AW3 (n = 3), b) AW4 (n = 1), c) AW5 (n = 3), and d) AW6 (n = 2).*

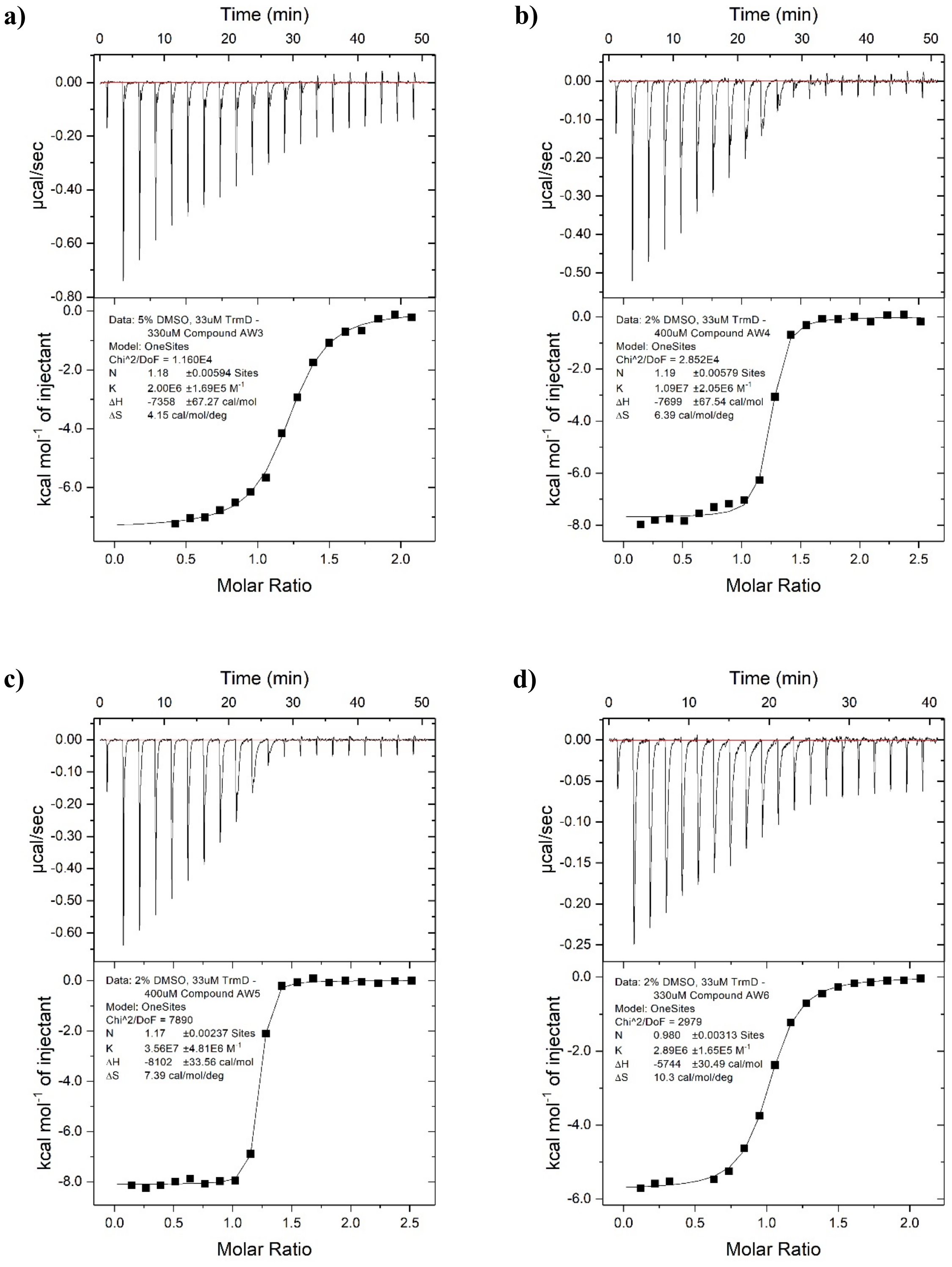

***Figure S12:*** *ITC trace with TrmD from Mycobacterium abscessus for a) AW7 (n = 3), and ITC traces with TrmD from Mycobacterium tuberculosis (n = 1) for b) AW6, and c) AW7.*

**a)**

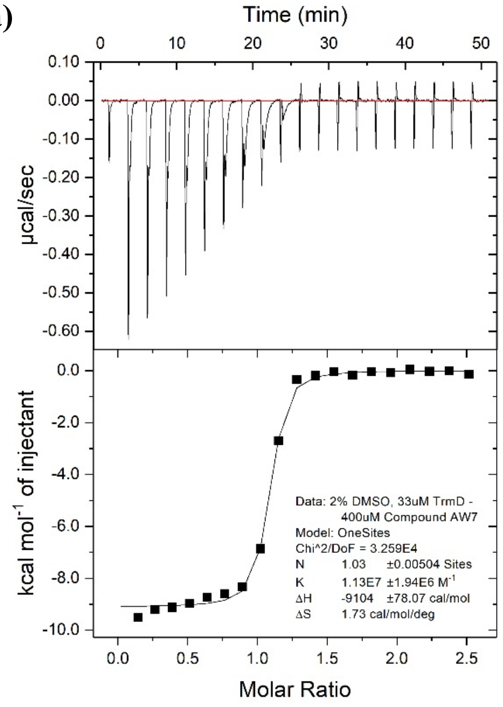


**b)**

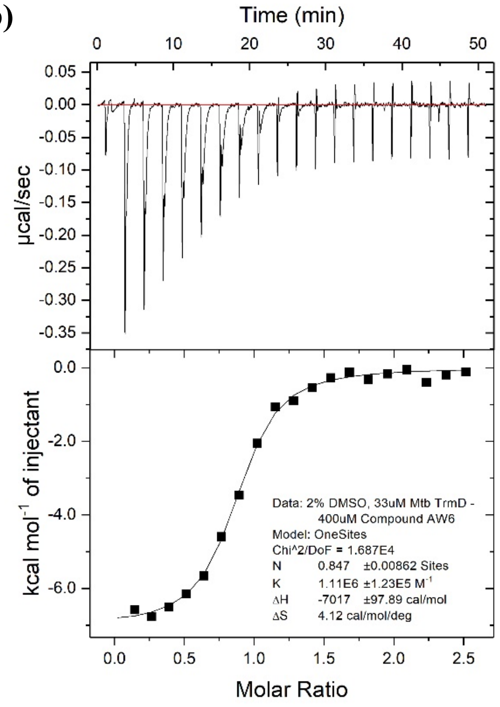


**c)**

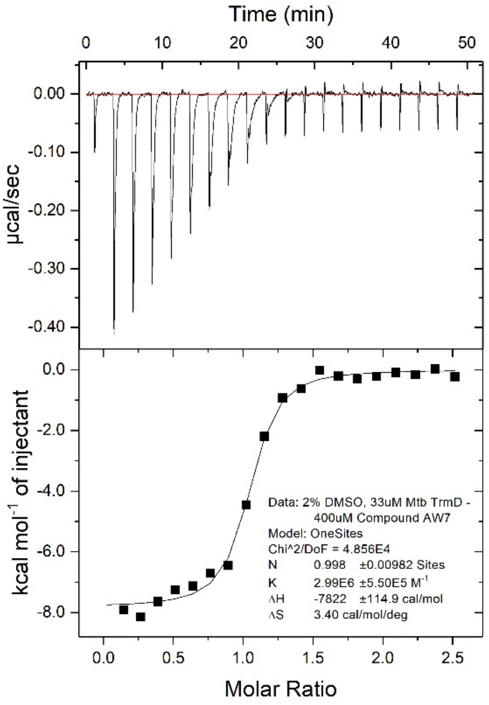

### Synthetic Chemistry

#### General Chemistry

All reactions were carried out in oven-dried glassware under a positive pressure of dry nitrogen atmosphere. Temperatures of 0 and -78 °C were obtained by submerging the reaction vessel in a bath containing either ice or a mixture of solid $CO_2$ pellets and acetone respectively. The solvents DCM, ethyl acetate, acetonitrile, methanol, petroleum ether and toluene were distilled over calcium hydride under a dry nitrogen atmosphere prior to use, with THF distilled over a mixture of lithium aluminium hydride, calcium hydride and triphenylphosphine. DMF was purchased as anhydrous from commercial suppliers, with ethanol and acetic acid obtained in the absolute and glacial forms respectively. All purchased chemicals were used as received. Solutions of $Na_2CO_3$, $NaHCO_3$, NaCl (brine) and $NH_4Cl$ were aqueous and saturated.

Flash column chromatography was performed using automated Biotage® Isolera™ Spektra purification systems with appropriately sized Biotage® SNAP cartridges, containing either KP 50 μm silica in 'normal phase' purification or HP-sphere 25 μm C18 silica in 'reverse phase' purification. Microwave-heating was performed using a Biotage® Initiator+ system with sealed Biotage® microwave reaction vials. Analytical thin layer chromatography (TLC) was performed using Merck glass-backed silica plates, with visualization by 254 or 365 nm ultraviolet light.

Liquid chromatography mass spectrometry (LCMS) was carried out using a Waters® Acquity UPLC® H-Class system, with samples run on a solvent gradient from 0 to 95% acetonitrile in water (+ 0.1% formic acid) over 4 minutes. Peaks corresponding to desired product are described, including the retention time (rt) and % purity by integration. High resolution mass spectrometry (HRMS) was mainly performed using ThermoFinnigan Orbitrap Classic, Waters® LCT Premier™ or Waters® Vion™ IMS QTof systems. A Perkin-Elmer® Spectrum One FT-IR spectrometer fitted with a universal attenuated total reflectance accessory was used to record infrared spectra, with wavelengths of maximum absorbance ($\nu_{max}$) quoted in wavenumbers ($cm^{-1}$) for signals outside of the fingerprint region (br = broad). Only peaks corresponding to key functional groups were characterized. Nuclear magnetic resonance (NMR) spectra were recorded in the indicated deuterated solvents with Avance™ III HD (400 MHz), QNP Cryoprobe (400 MHz) or DCH Cryoprobe (500 MHz) Bruker spectrometers. $^{1}H$ NMR data are presented in the following order: chemical shift (in ppm on a δ scale relative to the residual solvent resonance peak), integration, multiplicity (s = singlet, d = doublet, t = triplet, q = quartet, m = multiplet) and coupling constant (J, in Hz). $^{13}C$ NMR spectra were proton-decoupled, with chemical shifts recorded and further description provided for certain peaks (br = broad).

A combination of TLC and LCMS analysis was used to monitor reactions. All tested compounds possessed a purity of at least 95% as determined by LCMS analysis.

**Synthesis of AW1**

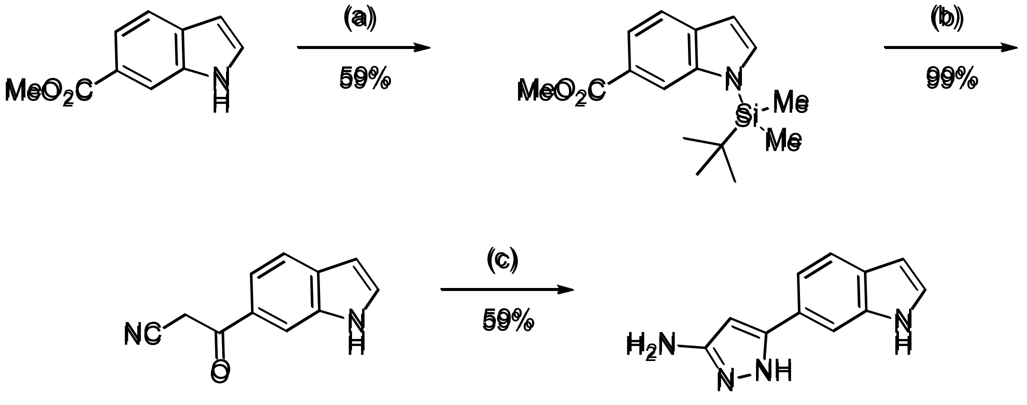


***Scheme S1:*** *Reagents and conditions: (a) tert-butyldimethylsilyl chloride (1 M in DCM), sodium hydride, THF, 0 °C to room temperature; (b) (i) acetonitrile, n-butyllithium (1.6 M in hexanes), toluene, -78 °C to room temperature (ii) tetra-n-butylammonium fluoride (1 M in THF), THF; (c) hydrazine monohydrate, ethanol, reflux.*

**methyl 1-(*tert*-butyldimethylsilyl)-1H-indole-6-carboxylate**

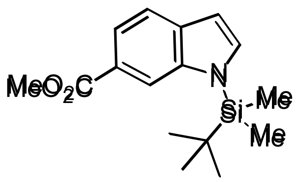


A solution of methyl 1H-indole-6-carboxylate (0.500 g, 2.71 mmol) in THF (10 mL) was added dropwise at 0 °C to a stirred suspension of sodium hydride (60% in mineral oil, 0.141 g, 3.52 mmol) in THF (10 mL). The reaction mixture was warmed to room temperature and stirred over 30 minutes. *tert*-Butyldimethylsilyl chloride (1 M in DCM, 4.1 mL, 4.1 mmol) was added dropwise at 0 °C to the reaction mixture. The reaction mixture was warmed to room temperature and stirred over 10 hours. $NH_4Cl$ solution (20 mL) was added dropwise at 0 °C. The product was extracted into diethyl ether (2 x 50 mL). The combined organic extracts were washed (brine), dried ($MgSO_4$) and concentrated *in vacuo*. Purification by flash column chromatography (0 – 20% ethyl acetate in petroleum ether) gave methyl 1-(*tert*-butyldimethylsilyl)-1H-indole-6-carboxylate (0.462 g, 59% yield).

LCMS (ESI+): m/z 290.2 $[M + H]^+$, rt 3.05 minutes, 100%; $^1H$ NMR (500 MHz, $CDCl_3$) 8.29-8.27 (1H, m), 7.80 (1H, dd, J = 8.3, 1.4 Hz), 7.63 (1H, d, J = 8.4 Hz), 7.34 (1H, d, J = 3.2 Hz),

6.65 (1H, dd, J = 3.1, 0.9 Hz), 3.93 (3H, s), 0.93 (9H, s), 0.65 (6H, s); $^{13}$C NMR (125 MHz, $CDCl_3$) 168.5, 140.5, 135.3, 134.6, 123.2, 121.0, 120.2, 116.2, 105.2, 52.1, 26.4, 19.5, -3.8; spectroscopic data consistent with literature. [1]

**3-(1H-indol-6-yl)-3-oxopropanenitrile**

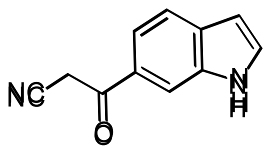

*n*-Butyllithium (1.6 M in hexanes, 3.0 mL, 4.8 mmol) was added dropwise at -78 °C to a mixture of acetonitrile (0.180 mL, 3.45 mmol) and toluene (3 mL). The reaction mixture was stirred at -78 °C over 30 minutes. A solution of methyl 1-(*tert*-butyldimethylsilyl)-1H-indole-6-carboxylate (0.200 g, 0.691 mmol) in toluene (2 mL) was added dropwise at -78 °C over 30 minutes. The reaction mixture was stirred at -78 °C over 1 hour, then warmed to room temperature over 1 hour. Aqueous HCl (3 M, 20 mL) was added dropwise at 0 °C. The intermediate was extracted into ethyl acetate (2 x 25 mL). The combined organic extracts were washed (brine), dried ($MgSO_4$) and concentrated *in vacuo*. The crude residue was dissolved in THF (2 mL) and tetra-*n*-butylammonium fluoride (1 M in THF, 0.760 mL, 0.760 mmol) was added dropwise. The reaction mixture was stirred over 20 minutes, then $NaHCO_3$ solution (30 mL) was added dropwise. The product was extracted into DCM (3 x 25 mL). The combined organic extracts were washed (brine), dried ($MgSO_4$) and concentrated *in vacuo*. Purification by flash column chromatography (0 – 20% methanol in DCM) gave 3-(1H-indol-6-yl)-3-oxopropanenitrile (0.131 g, 99% yield).

LCMS (ESI-): m/z 183.1 $[M - H]^-$, rt 1.91 minutes, 100%; $^1$H NMR (500 MHz, $CD_3CN$) 9.74 (1H, br s), 8.07 (1H, s), 7.69 (1H, d, J = 8.4 Hz), 7.62 (1H, dd, J = 8.4, 1.5 Hz), 7.53 (1H, t, J = 2.8 Hz), 6.61-6.57 (1H, m), 4.36 (2H, s); $^{13}$C NMR (125 MHz, $CD_3CN$) 189.5, 136.2, 133.6, 131.0, 129.3, 121.3, 120.1, 116.4, 113.9, 103.2, 30.6; $\nu_{max}$/cm$^{-1}$ 3326 (N-H), 1675 (C=O), 1611, 1501; HRMS (ESI)+: m/z calculated for $[C_{11}H_8N_2O + H]^+$ = 185.0709, observed 185.0708.

**5-(1H-indol-6-yl)-1H-pyrazol-3-amine (AW1)**

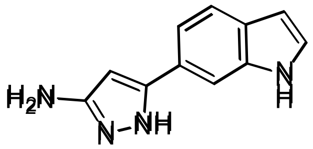


Hydrazine monohydrate (0.101 mL, 0.814 mmol) was added to a solution of 3-(1H-indol-6-yl)-3-oxopropanenitrile (50 mg, 0.27 mmol) in ethanol (5 mL). The reaction mixture was heated under reflux for 11 hours. Further hydrazine monohydrate (0.101 mL, 0.814 mmol) was added to the reaction mixture. The reaction mixture was heated under reflux for 1 hour, then concentrated *in vacuo*. Purification by flash column chromatography (0 – 10% methanol in DCM) gave 5-(1H-indol-6-yl)-1H-pyrazol-3-amine (32 mg, 59% yield).

LCMS (ESI+): m/z 199.1 $[M + H]^+$, (ESI-): m/z 197.0 $[M - H]^-$, rt 1.98 minutes, 100%; $^1H$ NMR (400 MHz, $CD_3OD$) 7.65 (1H, s), 7.56 (1H, d, J = 8.3 Hz), 7.30 (1H, dd, J = 8.3, 1.5 Hz), 7.26 (1H, d, J = 3.1 Hz), 6.44 (1H, dd, J = 3.1, 0.7 Hz), 5.93 (1H, s); $^{13}C$ NMR (125 MHz, $CD_3CN$) 155.2, 147.7, 137.1, 128.8, 126.9, 125.5, 121.4, 108.8, 102.5, 89.3 (1 peak missing); $\nu_{max}/cm^{-1}$ 3389 (N-H), 3250 (br, N-H), 1616, 1584, 1561, 1516; HRMS (ESI+): m/z calculated for $[C_{11}H_{10}N_4 + H]^+$ = 199.0978, observed 199.0973.

**Synthesis of AW2/ AW3**

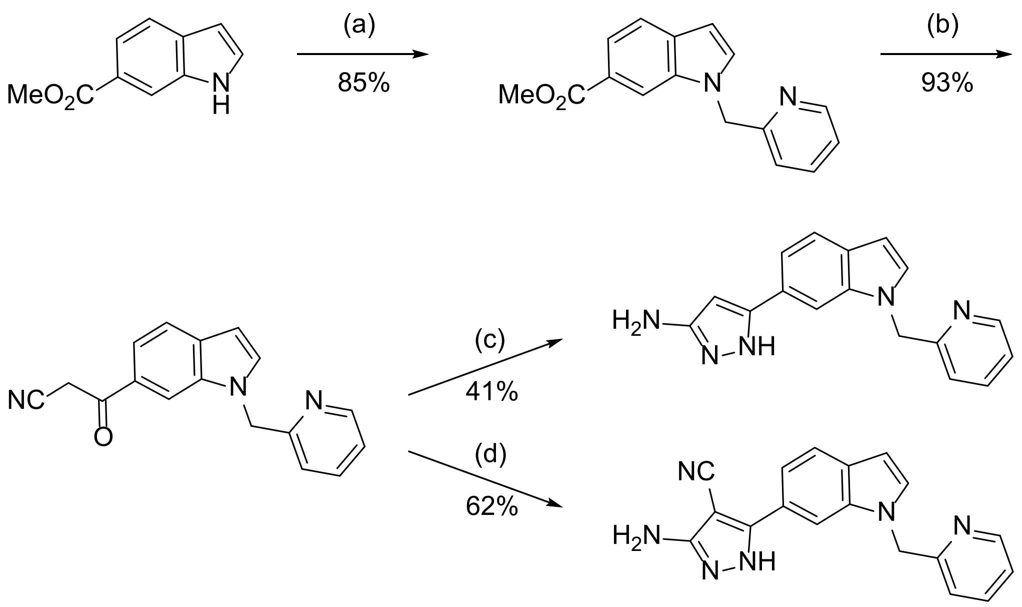


***Scheme S2:*** *Reagents and conditions: (a) 2-(bromomethyl)pyridine hydrobromide, caesium carbonate, acetonitrile, reflux; (b) acetonitrile, n-butyllithium (1.6 M in hexanes), THF, -78 °C; (c) hydrazine monohydrate, ethanol, reflux; (d) (i) trichloroacetonitrile, sodium acetate, ethanol (ii) hydrazine monohydrate, ethanol, reflux.*

**methyl 1-(pyridin-2-ylmethyl)-1H-indole-6-carboxylate**

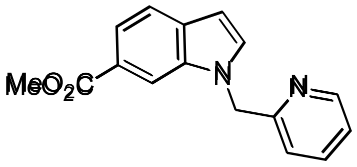

2-(Bromomethyl)pyridine hydrobromide (0.476 mg, 1.88 mmol) was added to a suspension of methyl 1H-indole-6-carboxylate (0.300 g, 1.71 mmol) and caesium carbonate (1.39 g, 4.28 mmol) in acetonitrile (15 mL). The reaction mixture was heated under reflux for 15 hours, then concentrated *in vacuo*. Water (20 mL) was added and the product extracted into DCM (3 x 25 mL). The combined organic extracts were washed (brine), dried ($MgSO_4$) and concentrated *in vacuo*. Purification by flash column chromatography (0 – 50% ethyl acetate in petroleum ether) gave methyl 1-(pyridin-2-ylmethyl)-1H-indole-6-carboxylate (0.386 g, 85% yield).

LCMS (ESI+): m/z 267.3 $[M + H]^+$, rt 1.82 minutes, 100%; $^1H$ NMR (400 MHz, $CDCl_3$) 8.60 (1H, dq, J = 4.9, 0.8 Hz), 8.09-8.05 (1H, m), 7.82 (1H, dd, J = 8.3, 1.4 Hz), 7.67 (1H, dd, J =

8.3, 0.5 Hz), 7.54 (1H, td, J = 7.7, 1.8 Hz), 7.37 (1H, d, J = 3.2 Hz), 7.21-7.14 (1H, m), 6.70 (1H, d, J = 7.8 Hz), 6.63 (1H, dd, J = 3.1, 0.8 Hz), 5.52 (2H, s), 3.90 (3H, s); $^{13}$C NMR (100 MHz, $CDCl_3$) 168.2, 157.1, 149.7, 137.3, 135.8, 132.5, 131.8, 123.8, 122.8, 120.9, 120.8, 112.1, 102.7, 52.10, 52.08 (1 peak missing); $\nu_{max}$/cm$^{-1}$ 1717 (C=O), 1593, 1505; HRMS (ESI)+: m/z calculated for $[C_{16}H_{14}N_2O_2 + H]^+$ = 267.1128, observed 267.1136.

**3-oxo-3-(1-(pyridin-2-ylmethyl)-1H-indol-6-yl)propanenitrile**

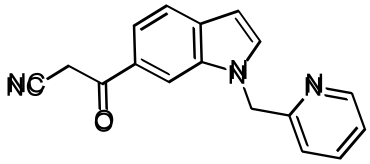


*n*-Butyllithium (1.6 M in hexanes, 2.6 mL, 4.2 mmol) was added dropwise at -78 °C to a mixture of acetonitrile (0.369 mL, 7.06 mmol) and THF (5 mL). The reaction mixture was stirred at -78 °C over 30 minutes. A solution of methyl 1-(pyridin-2-ylmethyl)-1H-indole-6-carboxylate (0.376 g, 1.41 mmol) in THF (5 mL) was added dropwise at -78 °C over 30 minutes. The reaction mixture was stirred at -78 °C over 30 minutes. Aqueous HCl (1 M, 15 mL) was added dropwise at 0 °C. The product was extracted into DCM (3 x 50 mL). The combined organic extracts were washed (brine), dried ($MgSO_4$) and concentrated *in vacuo* to afford 3-oxo-3-(1-(pyridin-2-ylmethyl)-1H-indol-6-yl)propanenitrile (0.363 g, 93% yield). LCMS (ESI+): m/z 276.2 $[M + H]^+$, (ESI-): m/z 274.2 $[M - H]^-$, rt 1.65 minutes, 100%; $^1$H NMR (400 MHz, $(CD_3)_2SO$) 8.55 (1H, d, J = 4.8 Hz), 8.19 (1H, s), 7.81 (1H, d, J = 3.1 Hz), 7.76 (1H, td, J = 7.6, 1.7 Hz), 7.70 (1H, d, J = 8.4 Hz), 7.61 (1H, dd, J = 8.4, 1.4 Hz), 7.31 (1H, dd, J = 7.5, 4.9 Hz), 7.06 (1H, d, J = 7.8 Hz), 6.64 (1H, d, J = 3.1 Hz), 5.66 (2H, s), 4.74 (2H, s); $^{13}$C NMR (100 MHz, $(CD_3)_2SO$) 189.2, 156.8, 149.1, 137.6, 135.3, 134.3, 132.7, 128.1, 122.9, 121.3, 120.5, 119.3, 116.3, 111.8, 102.0, 50.8, 29.9; $\nu_{max}$/cm$^{-1}$ 1673 (C=O), 1606, 1590, 1502; HRMS (ESI)+: m/z calculated for $[C_{17}H_{13}N_3O + H]^+$ = 276.1131, observed 276.1126.

**5-(1-(pyridin-2-ylmethyl)-1H-indol-6-yl)-1H-pyrazol-3-amine (AW2)**

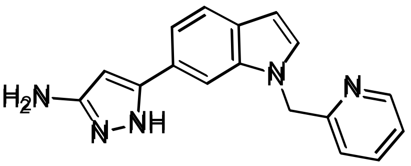


Hydrazine monohydrate (0.300 mL, 6.17 mmol) was added to a solution of 3-oxo-3-(1-(pyridin-2-ylmethyl)-1H-indol-6-yl)propanenitrile (0.170 g, 0.617 mmol) in ethanol (15 mL). The reaction mixture was heated under reflux for 13 hours, then concentrated *in vacuo*. Purification by flash column chromatography (0 – 20% methanol in DCM) gave 5-(1-(pyridin-2-ylmethyl)-1H-indol-6-yl)-1H-pyrazol-3-amine (73 mg, 41% yield).

LCMS (ESI+): m/z 290.3 $[M + H]^+$, rt 1.43 minutes, 100%; $^1$H NMR (400 MHz, $CD_3OD$) 8.56-8.50 (1H, m), 7.68 (1H, td, J = 7.7, 1.8 Hz), 7.63-7.58 (2H, m), 7.38-7.33 (2H, m), 7.28 (1H, dd, J = 7.6, 5.0 Hz), 6.89 (1H, d, J = 7.9 Hz), 6.55 (1H, d, J = 3.1 Hz), 5.90 (1H, s), 5.52 (2H, s); $^{13}$C NMR (125 MHz, $CD_3OD$) 158.8, 155.7 (br), 150.0, 148.4 (br), 139.2, 137.9, 130.9, 130.4, 125.8, 124.1, 122.7, 122.2, 118.7, 107.5, 103.1, 90.2, 52.2; $\nu_{max}/cm^{-1}$ 3150 (br, N-H), 2916, 1597, 1585, 1504; HRMS (ESI+): m/z calculated for $[C_{17}H_{15}N_5 + H]^+$ = 290.1400, observed 290.1405.

**3-amino-5-(1-(pyridin-2-ylmethyl)-1H-indol-6-yl)-1H-pyrazole-4-carbonitrile (AW3)**

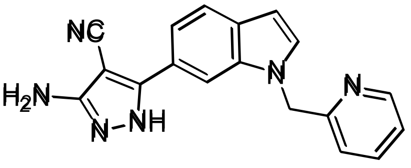


Trichloroacetonitrile (0.186 mL, 1.85 mmol) was added to a suspension of 3-oxo-3-(1-(pyridin-2-ylmethyl)-1H-indol-6-yl) propanenitrile (0.170 g, 0.617 mmol) and sodium acetate (0.253 g, 3.09 mmol) in ethanol (5 mL). The reaction mixture was stirred over 90 minutes. The reaction mixture was concentrated *in vacuo*, then water (20 mL) was added. The intermediate was extracted into DCM (3 x 25 mL). The combined organic extracts were washed (brine), dried ($MgSO_4$) and concentrated *in vacuo*. The crude residue was dissolved in ethanol (10 mL) and hydrazine monohydrate (0.300 mL, 6.17 mmol) was added. The reaction mixture was heated under reflux for 15 hours, then concentrated *in vacuo*. Purification by flash column chromatography (0 - 100% ethyl acetate in petroleum ether, 0 – 10% methanol in DCM) gave 3-amino-5-(1-(pyridin-2-ylmethyl)-1H-indol-6-yl)-1H-pyrazole-4-carbonitrile (0.121 g, 62% yield).

LCMS (ESI+): m/z 315.2 $[M + H]^+$, (ESI-): m/z 313.1 $[M - H]^-$, rt 1.53 minutes, 100%; $^1H$ NMR (400 MHz, $CDCl_3$) 8.46 (1H, d, J = 5.0 Hz), 7.70 (1H, s), 7.67 (1H, d, J = 8.3 Hz), 7.62-7.53 (2H, m), 7.31 (1H, d, J = 3.1 Hz), 7.14 (1H, dd, J = 7.6, 4.9 Hz), 6.91 (1H, d, J = 7.8 Hz), 6.58 (1H, d, J = 3.2 Hz), 5.38 (2H, s); $^{13}C$ NMR (125 MHz, $CDCl_3$) 157.8, 156.7, 149.2, 149.1, 137.8, 136.1, 130.9, 130.4, 123.3, 122.1, 121.8, 121.0, 118.1, 115.6, 107.6, 102.7, 76.2, 52.3; $\nu_{max}/cm^{-1}$ 3200 (br, N-H), 2211 (C≡N), 1621, 1592, 1503; HRMS (ESI+): m/z calculated for $[C_{18}H_{14}N_6 + H]^+$ = 315.1353, observed 315.1338.

#### Synthesis of AW4

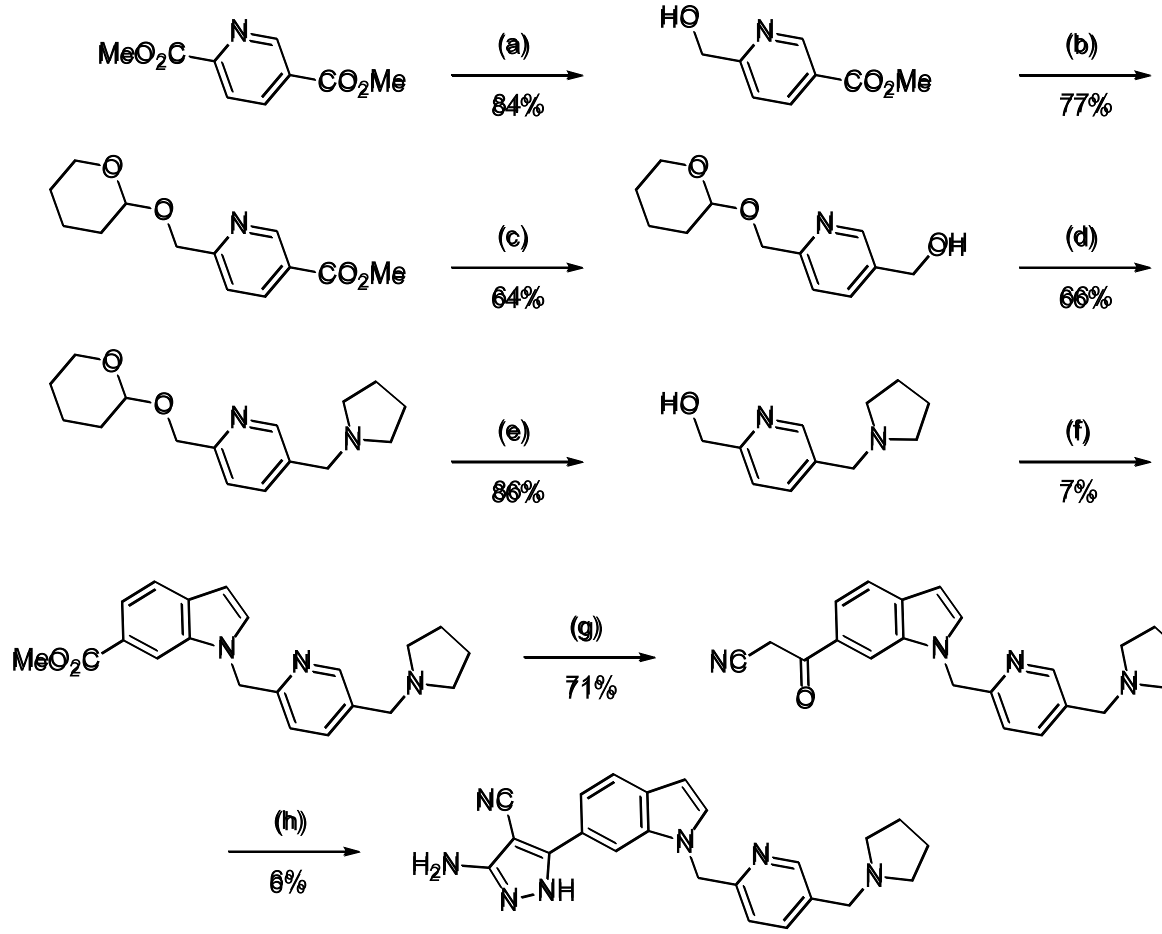

***Scheme S3:*** *Reagents and conditions: (a) sodium borohydride, $CaCl_2$, methanol, THF, 0 °C; (b) methanesulfonic acid, 3,4-dihydro-2H-pyran, DCM; (c) lithium aluminium hydride (2.4 M in THF), THF, 0 °C; (d) (i) methanesulfonyl chloride, triethylamine, DCM, 0 °C to room temperature (ii) pyrrolidine, caesium carbonate, DMF; (e) p-toluenesulfonic acid monohydrate, ethanol, 50 °C; (f) (i) methanesulfonyl chloride, triethylamine, DCM, 0 °C to room temperature (ii) methyl 1H-indole-6-carboxylate, sodium hydride, NaI, DMF, 0 to 60 °C (iii) methanol, sulfuric acid, reflux; (g) acetonitrile, n-butyllithium (1.6 M in hexanes), THF, -78 °C; (h) (i) trichloroacetonitrile, sodium acetate, ethanol (ii) hydrazine monohydrate, ethanol, reflux.*

##### methyl 6-(hydroxymethyl)nicotinate [2]

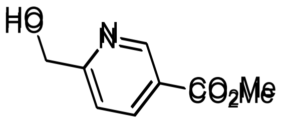

Sodium borohydride (0.727 g, 19.2 mmol) was added portionwise at 0 °C to a suspension of dimethyl pyridine-2,5-dicarboxylate (2.50 g, 12.8 mmol) and $CaCl_2$ (5.69 g, 51.2 mmol) in methanol/THF (2:1, 90 mL). The reaction mixture was stirred at 0 °C over 90 minutes, then diluted with water (50 mL) dropwise at 0 °C. The product was extracted into chloroform (3 x

50 mL). The combined organic extracts were washed (brine), dried ($MgSO_4$) and concentrated *in vacuo* to afford methyl 6-(hydroxymethyl)nicotinate (1.80 g, 84% yield).

$^1$H NMR (400 MHz, $CDCl_3$) 9.15 (1H, d, J = 1.4 Hz), 8.29 (1H, dd, J = 8.2, 2.1 Hz), 7.36 (1H, d, J = 8.2 Hz), 4.83 (2H, d, J = 3.6 Hz), 3.95 (3H, s), 3.75-3.65 (1H, m); $^{13}$C NMR (100 MHz, $CDCl_3$) 165.8, 163.7, 150.1, 137.9, 125.1, 120.1, 64.4, 52.6; spectroscopic data consistent with literature. [2]

**methyl 6-(((tetrahydro-2H-pyran-2-yl)oxy)methyl)nicotinate**

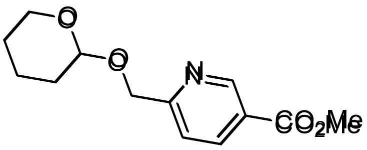


Methanesulfonic acid (0.763 mL, 11.8 mmol) was added to a solution of methyl 6-(hydroxymethyl)nicotinate (1.79 g, 10.7 mmol) and 3,4-dihydro-2H-pyran (1.95 mL, 21.4 mmol) in DCM (20 mL). The reaction mixture was stirred over 150 minutes, then washed with $NaHCO_3$ solution (2 x 15 mL) and brine (15 mL), dried ($MgSO_4$) and concentrated *in vacuo*. Purification by flash column chromatography (0 – 20% ethyl acetate in petroleum ether) gave methyl 6-(((tetrahydro-2H-pyran-2-yl)oxy)methyl)nicotinate (2.07 g, 77% yield).

$^1$H NMR (400 MHz, $CDCl_3$) 9.14 (1H, d, J = 2.1 Hz), 8.29 (1H, dd, J = 8.1, 2.2 Hz), 7.58 (1H, dd, J = 8.2, 0.7 Hz), 4.95 (1H, d, J = 14.6 Hz), 4.78 (1H, t, J = 3.5 Hz), 4.70 (1H, d, J = 14.5 Hz), 3.94 (3H, s), 3.93-3.84 (1H, m), 3.60-3.52 (1H, m), 1.97-1.47 (6H, m); $^{13}$C NMR (100 MHz, $CDCl_3$) 166.0, 163.5, 150.5, 137.8, 124.7, 120.7, 98.8, 69.7, 62.5, 52.5, 30.6, 25.5, 19.5; $\nu_{max}$/cm$^{-1}$ 2946, 2926, 2876, 2848, 1714 (C=O), 1596; HRMS (ESI)+: m/z calculated for $[C_{13}H_{17}NO_4 + Na]^+$ = 274.1050, observed 274.1045.

**(6-(((tetrahydro-2H-pyran-2-yl)oxy)methyl)pyridin-3-yl)methanol**

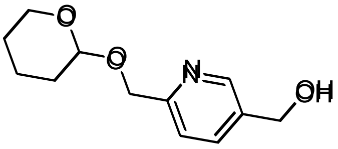

Lithium aluminium hydride (2.4 M in THF, 5.1 mL, 12 mmol) was added dropwise at 0 °C to a solution of methyl 6-(((tetrahydro-2H-pyran-2-yl)oxy)methyl)nicotinate (2.06 g, 8.20 mmol) in THF (20 mL). The reaction mixture was stirred at 0 °C for 45 minutes. Isopropanol (5 mL) was added dropwise at 0 °C to the reaction mixture, followed by water (50 mL). The product was extracted into DCM/methanol (9:1, 3 x 100 mL). The combined organic extracts were washed (brine), dried ($MgSO_4$) and concentrated *in vacuo*. Purification by flash column chromatography (60 – 90% ethyl acetate in petroleum ether) gave (6-(((tetrahydro-2H-pyran-2-yl)oxy)methyl)pyridin-3-yl)methanol (1.16 g, 64% yield).

$^1$H NMR (400 MHz, $CDCl_3$) 8.50 (1H, d, J = 1.7 Hz), 7.74 (1H, dd, J = 7.9, 2.2 Hz), 7.48 (1H, d, J = 8.1 Hz), 4.90 (1H, d, J = 13.3 Hz), 4.78 (1H, t, J = 3.6 Hz), 4.73 (2H, s), 4.65 (1H, d, J = 13.3 Hz), 3.97-3.88 (1H, m), 3.61-3.53 (1H, m), 2.60 (1H, br s), 1.91-1.49 (6H, m); $^{13}$C NMR (100 MHz, $CDCl_3$) 158.0, 148.0, 135.7, 135.0, 121.5, 98.7, 69.8, 62.7, 62.4, 30.7, 25.5, 19.5; $\nu_{max}$/cm$^{-1}$ 3300 (br, O-H), 2939, 2869, 1604, 1574; HRMS (ESI)+: m/z calculated for $[C_{12}H_{17}NO_3 + H]^+$ = 224.1281, observed 224.1284.

**5-(pyrrolidin-1-ylmethyl)-2-(((tetrahydro-2H-pyran-2-yl)oxy)methyl)pyridine**

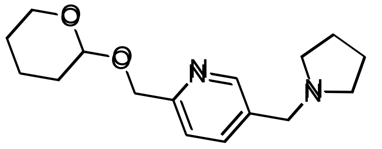

Methanesulfonyl chloride (0.478 mL, 6.18 mmol) was added dropwise at 0 °C to a solution of (6-(((tetrahydro-2H-pyran-2-yl)oxy)methyl)pyridin-3-yl)methanol (1.15 g, 5.15 mmol) and triethylamine (0.933 mL, 6.70 mmol) in DCM (10 mL). The reaction mixture was warmed to room temperature and stirred over 1 hour. The reaction was diluted with $NaHCO_3$ solution (25 mL) and extracted into DCM (3 x 50 mL). The combined organic extracts were washed (brine), dried ($MgSO_4$) and concentrated *in vacuo*. The crude residue was dissolved in DMF (10 mL), then pyrrolidine (0.516 mL, 6.18 mmol) and caesium carbonate (2.18 g, 6.70 mmol) were added. The reaction mixture was stirred over 14 hours, then diluted with ethyl acetate (100 mL), washed with $Na_2CO_3$ solution (2 x 100 mL) and brine (100 mL), dried ($MgSO_4$) and concentrated *in vacuo*. Purification by flash column chromatography (0 – 8% methanol in DCM) gave 5-(pyrrolidin-1-ylmethyl)-2-(((tetrahydro-2H-pyran-2-yl)oxy)methyl)pyridine (0.970 g, 66% yield).

LCMS (ESI+): m/z 277.3 $[M + H]^+$, rt 0.65 minutes, 97%; $^1H$ NMR (400 MHz, $CDCl_3$) 8.48 (1H, s), 7.68 (1H, dd, J = 7.9, 1.7 Hz), 7.42 (1H, d, J = 7.9 Hz), 4.88 (1H, d, J = 13.2 Hz), 4.77 (1H, t, J = 3.4 Hz), 4.62 (1H, d, J = 13.3 Hz), 3.96-3.85 (1H, m), 3.61 (2H, s), 3.58-3.51 (1H, m), 2.55-2.43 (4H, m), 1.95-1.48 (10H, m); $^{13}C$ NMR (100 MHz, $CDCl_3$) 157.3, 149.6, 137.3, 133.4, 121.3, 98.6, 69.9, 62.3, 57.8, 54.2, 30.7, 25.5, 23.5, 19.5; $\nu_{max}/cm^{-1}$ 2939, 2783, 1602, 1572; HRMS (ESI)+: m/z calculated for $[C_{16}H_{24}N_2O_2 + H]^+$ = 277.1911, observed 277.1906.

**(5-(pyrrolidin-1-ylmethyl)pyridin-2-yl)methanol**

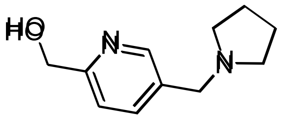


*p*-Toluenesulfonic acid monohydrate (1.28 g, 6.74 mmol) was added to a solution of 5-(pyrrolidin-1-ylmethyl)-2-(((tetrahydro-2H-pyran-2-yl)oxy)methyl)pyridine (0.960 g, 3.37 mmol) in ethanol (10 mL). The reaction mixture was heated to 50 °C over 30 minutes, then diluted with water (15 mL), adjusted to pH 14 using aqueous NaOH (10% w/v) and extracted into DCM (6 x 25 mL). The combined organic extracts were washed (brine), dried ($MgSO_4$) and concentrated *in vacuo*. Purification by flash column chromatography (0 – 15% methanol (+ 0.1% $NH_3$) in DCM) gave (5-(pyrrolidin-1-ylmethyl)pyridin-2-yl)methanol (0.560 g, 86% yield).

$^1H$ NMR (400 MHz, $CDCl_3$) 8.48 (1H, d, J = 1.5 Hz), 7.69 (1H, dd, J = 7.9, 2.0 Hz), 7.21 (1H, d, J = 8.1 Hz), 4.74 (2H, s), 3.62 (2H, s), 2.57-2.44 (4H, m), 1.85-1.73 (4H, m); $^{13}C$ NMR (100 MHz, $CDCl_3$) 158.0, 148.9, 137.5, 133.6, 120.3, 64.2, 57.7, 54.2, 23.6; $\nu_{max}/cm^{-1}$ 3200 (br, O-

H), 2963, 2794, 1603, 1572; HRMS (ESI)+: m/z calculated for $[C_{11}H_{16}N_2O + H]^+$ = 193.1335, observed 193.1335.

**methyl 1-((5-(pyrrolidin-1-ylmethyl)pyridin-2-yl)methyl)-1H-indole-6-carboxylate**

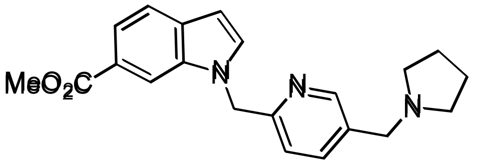


Methanesulfonyl chloride (0.242 mL, 3.12 mmol) was added dropwise at 0 °C to a solution of (5-(pyrrolidin-1-ylmethyl)pyridin-2-yl)methanol (0.500 g, 2.60 mmol) and triethylamine (0.471 mL, 3.38 mmol) in DCM (10 mL). The reaction mixture was warmed to room temperature and stirred over 150 minutes. $Na_2CO_3$ solution (15 mL) was added dropwise at 0 °C to the reaction mixture. The intermediate was extracted into DCM (4 x 25 mL). The combined organic extracts were washed (brine), dried ($MgSO_4$) and concentrated *in vacuo* to afford the intermediate as a crude residue. A solution of methyl 1H-indole-6-carboxylate (0.547 g, 3.12 mmol) in DMF (3 mL) was added dropwise at 0 °C to a suspension of sodium hydride (60% in mineral oil, 0.520 g, 13.0 mmol) in DMF (2 mL). The reaction mixture was warmed to room temperature over 30 minutes. The crude residue was dissolved in DMF (5 mL) and added dropwise at 0 °C to the reaction mixture. The reaction mixture was warmed to room temperature and stirred over 45 minutes. Sodium iodide (39 mg, 0.26 mmol) was added to the reaction mixture, and it was heated to 60 °C over 30 minutes. Methanol (150 mL) was added dropwise at 0 °C to the reaction mixture followed by sulfuric acid (10 mL). The reaction mixture was heated under reflux for 14 hours, then concentrated *in vacuo*. $Na_2CO_3$ solution (200 mL) was added dropwise to the reaction mixture, followed by water (50 mL). The product was extracted into ethyl acetate (2 x 200 mL), followed by DCM/methanol (9:1, 3 x 200 mL). The combined organic extracts were washed (brine), dried ($MgSO_4$) and concentrated *in vacuo*. Purification by flash column chromatography (0 – 100% ethyl acetate in petroleum ether, 0 –

10% methanol in DCM) gave methyl 1-((5-(pyrrolidin-1-ylmethyl)pyridin-2-yl)methyl)-1H-indole-6-carboxylate (95 mg, 7% yield).

LCMS (ESI+): m/z 350.3 $[M + H]^+$, rt 1.39 minutes, 84%; $^1$H NMR (400 MHz, $CDCl_3$) 8.46 (1H, d, J = 1.6 Hz), 8.07 (1H, s), 7.82 (1H, dd, J = 8.4, 1.4 Hz), 7.66 (1H, d, J = 8.4 Hz), 7.36 (1H, d, J = 7.9 Hz), 7.31 (1H, dd, J = 8.0, 2.2 Hz), 7.27-7.24 (1H, m), 6.61 (1H, dd, J = 3.1, 0.8 Hz), 5.39 (2H, s), 3.92 (3H, s), 3.79 (2H, s), 2.61 (4H, br s), 1.87-1.75 (4H, m); $^{13}$C NMR (100 MHz, $CDCl_3$) 168.0, 158.4, 147.6, 135.5, 135.1, 132.4, 131.10, 131.08, 123.6, 123.4, 120.9, 120.7, 111.7, 102.7, 61.5, 54.1, 52.0, 47.4, 23.5; $\nu_{max}/cm^{-1}$ 2951, 2799, 1705 (C=O), 1615, 1602, 1569, 1505; HRMS (ESI)+: m/z calculated for $[C_{21}H_{23}N_3O_2 + H]^+$ = 350.1863, observed 350.1863.

**3-oxo-3-(1-((5-(pyrrolidin-1-ylmethyl)pyridin-2-yl)methyl)-1H-indol-6-yl)propanenitrile**

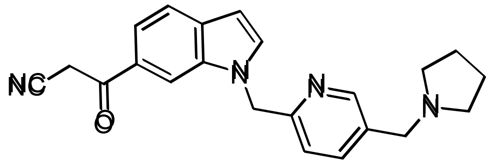

*n*-Butyllithium (1.6 M in hexanes, 0.42 mL, 0.67 mmol) was added dropwise at -78 °C to a mixture of acetonitrile (58 µL, 1.1 mmol) and THF (2 mL). The reaction mixture was stirred at -78 °C over 30 minutes. A solution of methyl 1-((5-(pyrrolidin-1-ylmethyl)pyridin-2-yl)methyl)-1H-indole-6-carboxylate (92 mg, 0.22 mmol) in THF (3 mL) was added dropwise at -78 °C over 30 minutes. The reaction mixture was stirred at -78 °C over 1 hour. Water (15 mL) was added dropwise at 0 °C, and the reaction mixture adjusted to pH 8. The product was extracted into DCM (3 x 25 mL), followed by DCM/methanol (9:1, 3 x 25 mL). The combined organic extracts were washed (brine), dried ($MgSO_4$) and concentrated *in vacuo*. Purification by flash column chromatography (0 – 20% methanol in DCM) gave 3-oxo-3-(1-((5-(pyrrolidin-1-ylmethyl)pyridin-2-yl)methyl)-1H-indol-6-yl)propanenitrile (69 mg, 71% yield).

LCMS (ESI+): m/z 359.2 $[M + H]^+$, (ESI-): m/z 357.1 $[M - H]^-$, rt 1.30 minutes, 81%; $^1H$ NMR (400 MHz, $CDCl_3$) 8.38 (1H, s), 7.96 (1H, s), 7.66 (1H, d, J = 8.4 Hz), 7.56 (1H, d, J = 8.3 Hz), 7.42-7.29 (3H, m), 6.60 (1H, d, J = 2.8 Hz), 5.37 (2H, s), 4.13 (2H, br s), 3.79 (2H, s), 2.70-2.55 (4H, m), 1.85-1.70 (4H, m); $^{13}C$ NMR (100 MHz, $CDCl_3$) 187.1, 158.3, 147.8, 135.7, 135.2, 133.7, 132.9, 130.8, 128.3, 123.5, 121.4, 119.8, 114.6, 110.9, 103.1, 61.4, 54.2, 47.7, 29.8, 23.5; $\nu_{max}/cm^{-1}$ 2919, 2799, 2168 (C≡N), 1675 (C=O), 1605; HRMS (ESI)+: m/z calculated for $[C_{22}H_{22}N_4O + H]^+$ = 359.1866, observed 359.1866.

**3-amino-5-(1-((5-(pyrrolidin-1-ylmethyl)pyridin-2-yl)methyl)-1H-indol-6-yl)-1H-pyrazole-4-carbonitrile (AW4)**

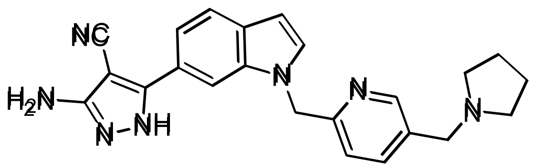


Trichloroacetonitrile (45 μL, 0.45 mmol) was added to a suspension of 3-oxo-3-(1-((5-(pyrrolidin-1-ylmethyl)pyridin-2-yl)methyl)-1H-indol-6-yl)propanenitrile (66 mg, 0.15 mmol) and sodium acetate (61 mg, 0.75 mmol) in ethanol (4 mL). The reaction mixture was stirred over 36 hours, then $Na_2CO_3$ solution (12.5 mL) and water (12.5 mL) were added. The intermediate was extracted into DCM (2 x 25 mL) and methanol/DCM (9:1, 4 x 25 mL). The combined organic extracts were washed (brine), dried ($MgSO_4$) and concentrated *in vacuo*. The crude residue was dissolved in ethanol (5 mL) and hydrazine monohydrate (73 μL, 1.5 mmol) was added. The reaction mixture was heated under reflux for 24 hours, then quenched with excess acetone at room temperature and concentrated *in vacuo*. Purification by flash column chromatography (0 – 20% methanol (+ 0.1% $NH_3$) in DCM) gave 3-amino-5-(1-((5-(pyrrolidin-1-ylmethyl)pyridin-2-yl)methyl)-1H-indol-6-yl)-1H-pyrazole-4-carbonitrile (3.6 mg, 6% yield).

LCMS (ESI+): m/z 398.3 $[M + H]^+$, (ESI-): m/z 396.2 $[M - H]^-$, rt 1.32 minutes, 100%; $^1H$ NMR (500 MHz, $CD_3OD$) 8.41 (1H, d, J = 1.9 Hz), 7.83 (1H, s), 7.66 (1H, d, J = 8.4 Hz), 7.64

(1H, dd, J = 8.1, 2.3 Hz), 7.51 (1H, d, J = 7.9 Hz), 7.47 (1H, d, J = 3.2 Hz), 7.42 (1H, d, J = 8.2 Hz), 6.58 (1H, d, J = 3.1 Hz), 5.49 (2H, s), 3.79 (2H, s), 2.67-2.59 (4H, m), 1.83-1.76 (4H, m); $^{13}C$ NMR (125 MHz, $CD_3OD$) 158.3, 148.5, 137.5, 137.2, 134.2, 131.5, 125.0, 122.4, 119.0, 117.0, 109.0, 103.2, 61.9, 55.1, 48.2, 24.2 (5 peaks missing); $\nu_{max}/cm^{-1}$ 3150 (br, N-H), 2918, 2209 (C≡N), 1602, 1501; HRMS (ESI)+: m/z calculated for $[C_{23}H_{23}N_7 + H]^+$ = 398.2088, observed 398.2082.

**Synthesis of AW5-AW7**

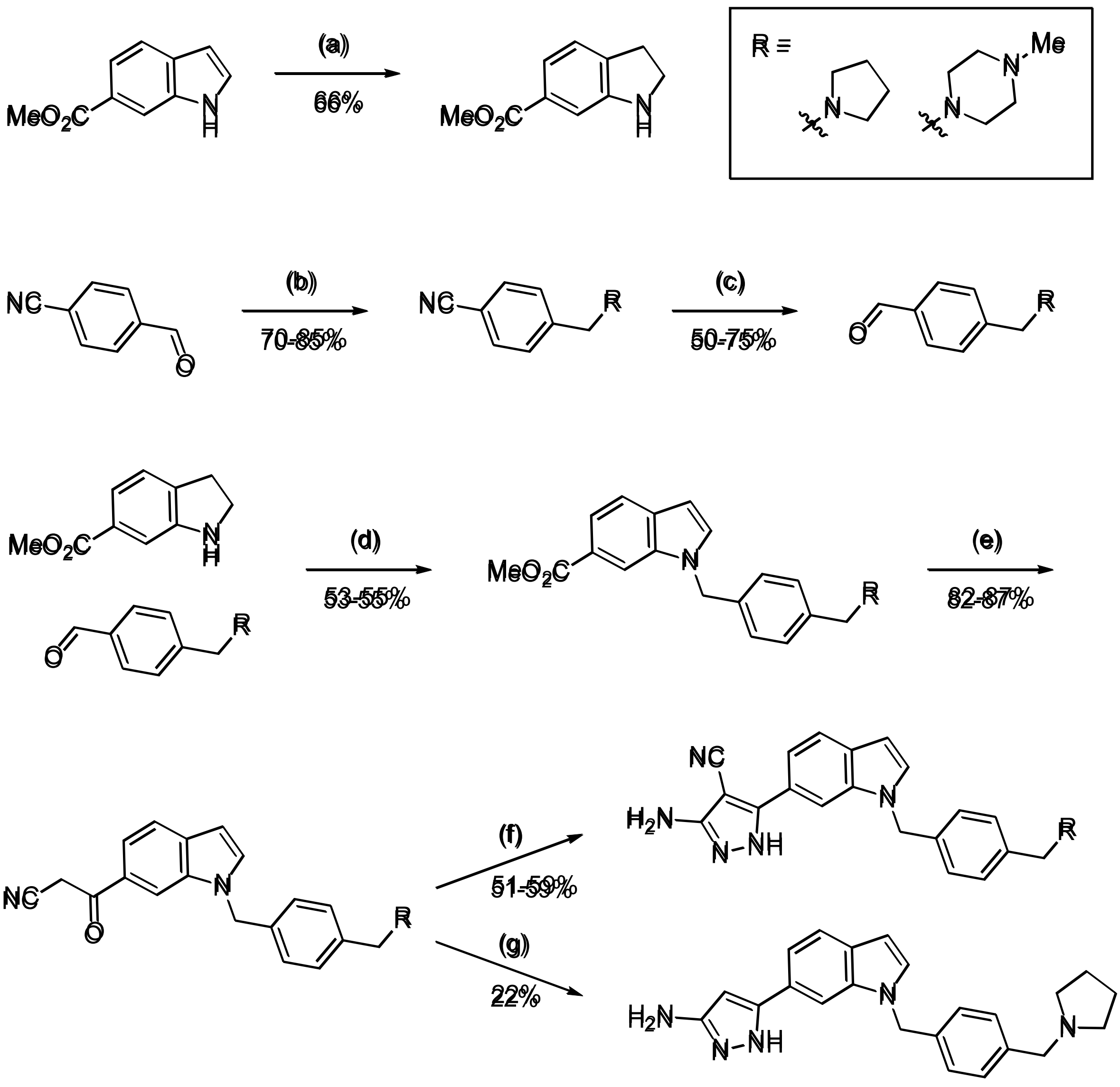


*Scheme S4: Reagents and conditions: (a) sodium cyanoborohydride, acetic acid, 0 °C to room temperature; (b) sodium triacetoxyborohydride, RH, acetic acid, DCM; (c) diisobutylaluminium hydride (1 M in THF), THF, 0 °C to room temperature; (d) benzoic acid, toluene, 200 °C µW; (e) acetonitrile, n-butyllithium (1.6 M in hexanes), THF, -78 °C; (f) (i) trichloroacetonitrile, sodium acetate, ethanol (ii) hydrazine monohydrate, ethanol, reflux; (g) hydrazine monohydrate, ethanol, reflux.*

#### methyl indoline-6-carboxylate [3]

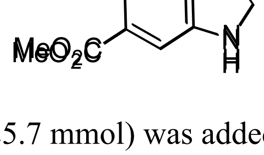


Sodium cyanoborohydride (1.61 g, 25.7 mmol) was added portionwise at 0 °C to a solution of methyl 1H-indole-6-carboxylate (1.50 g, 8.56 mmol) in acetic acid (15 mL) over 30 minutes.

The reaction mixture was warmed to room temperature and stirred over 7 hours. The reaction was diluted with ethyl acetate (100 mL), washed with $NaHCO_3$ solution (3 x 100 mL) and brine (100 mL), dried ($MgSO_4$) and concentrated *in vacuo*. Purification by flash column chromatography (0 – 30% ethyl acetate in petroleum ether) gave methyl indoline-6-carboxylate (1.11 g, 66% yield).

LCMS (ESI+): m/z 178.2 $[M + H]^+$, rt 1.00 minutes, 90%; $^1H$ NMR (400 MHz, $CDCl_3$) 7.40 (1H, dd, J = 7.6, 1.4 Hz), 7.23 (1H, d, J = 1.4 Hz), 7.12 (1H, d, J = 7.6 Hz), 3.90 (1H, br s), 3.86 (3H, s), 3.57 (2H, t, J = 8.4 Hz), 3.04 (2H, t, J = 8.4 Hz); $^{13}C$ NMR (100 MHz, $CDCl_3$) 167.6, 151.9, 135.0, 129.4, 124.3, 120.7, 109.6, 52.0, 47.5, 29.9; $^1H$ NMR spectroscopic data consistent with literature.[3]

**4-(pyrrolidin-1-ylmethyl)benzonitrile**

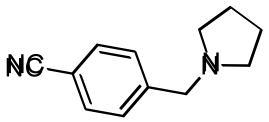


Sodium triacetoxyborohydride (1.29 g, 6.10 mmol) was added to a solution of 4-formylbenzonitrile (0.500 g, 3.81 mmol), pyrrolidine (0.318 mL, 3.81 mmol) and acetic acid (0.262 mL, 4.58 mmol) in DCM (10 mL). The reaction mixture was stirred over 15 hours. The reaction was diluted with water (25 mL), adjusted to pH 14 using aqueous NaOH (6 M) and extracted into DCM (3 x 25 mL). The combined organic extracts were dried ($MgSO_4$) and concentrated *in vacuo*. Purification by flash column chromatography (0 – 5% methanol in DCM) gave 4-(pyrrolidin-1-ylmethyl)benzonitrile (0.645 g, 85% yield).

$^1H$ NMR (400 MHz, $CDCl_3$) 7.62-7.56 (2H, m), 7.47-7.41 (2H, m), 3.65 (2H, s), 2.55-2.44 (4H, m), 1.84-1.73 (4H, m); $^{13}C$ NMR (100 MHz, $CDCl_3$) 145.4, 132.2, 129.4, 119.2, 110.8, 60.3, 54.4, 23.6; spectroscopic data consistent with literature.[4]

**4-(pyrrolidin-1-ylmethyl)benzaldehyde**

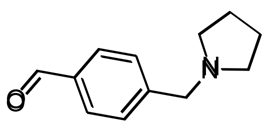

Diisobutylaluminium hydride (1 M in THF, 2.2 mL, 2.2 mmol) was added dropwise at 0 °C to a solution of 4-(pyrrolidin-1-ylmethyl)benzonitrile (0.408 g, 2.19 mmol) in THF (5 mL). The reaction mixture was warmed to room temperature and stirred over 45 minutes. Further diisobutylaluminium hydride (2.2 mL, 2.2 mmol) was added dropwise at 0 °C to the reaction mixture. The reaction mixture was warmed to room temperature and stirred over 15 minutes, then $NaHCO_3$ solution (50 mL) was added dropwise at 0 °C. The product was extracted into DCM/methanol (10:1, 3 x 50 mL). The combined organic extracts were washed (brine), dried ($MgSO_4$) and concentrated *in vacuo*. Purification by flash column chromatography (0 – 4% methanol (+ 0.1% $NH_3$) in DCM) gave 4-(pyrrolidin-1-ylmethyl)benzaldehyde (0.313 g, 75% yield).

$^1$H NMR (400 MHz, $CDCl_3$) 9.99 (1H, s), 7.83 (2H, d, J = 8.2 Hz), 7.50 (2H, d, J = 8.0 Hz), 3.68 (2H, s), 2.56-2.48 (4H, m), 1.84-1.75 (4H, m); $^{13}$C NMR (100 MHz, $CDCl_3$) 192.2, 147.0, 135.5, 129.9, 129.4, 60.6, 54.4, 23.7; spectroscopic data consistent with literature.[5]

**methyl 1-(4-(pyrrolidin-1-ylmethyl)benzyl)-1H-indole-6-carboxylate**

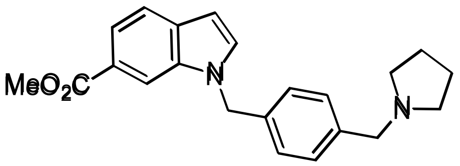


A solution of 4-(pyrrolidin-1-ylmethyl)benzaldehyde (0.302 g, 1.60 mmol) in toluene (3 mL) was added to a mixture of methyl indoline-6-carboxylate (0.377 g, 1.91 mmol) and benzoic acid (39 mg, 0.32 mmol). The reaction mixture was heated to 200 °C by microwave for 20 minutes. The reaction was diluted with ethyl acetate (50 mL), washed with $NaHCO_3$ solution (3 x 50 mL) and brine (50 mL), dried ($MgSO_4$) and concentrated *in vacuo*. Purification by flash column chromatography (40 – 80% ethyl acetate in petroleum ether) gave methyl 1-(4-(pyrrolidin-1-ylmethyl)benzyl)-1H-indole-6-carboxylate (0.297 g, 53% yield).

LCMS (ESI+): m/z 349.3 $[M + H]^+$, rt 1.51 minutes, 100%; $^1H$ NMR (400 MHz, $CDCl_3$) 8.11 (1H, s), 7.81 (1H, dd, J = 8.3, 1.3 Hz), 7.66 (1H, d, J = 8.4 Hz), 7.32-7.23 (3H, m), 7.07 (2H, d, J = 7.9 Hz), 6.59 (1H, dd, J = 3.1, 0.5 Hz), 5.37 (2H, s), 3.92 (3H, s), 3.58 (2H, s), 2.52-2.44 (4H, m), 1.80-1.73 (4H, m); $^{13}C$ NMR (100 MHz, $CDCl_3$) 168.3, 139.3, 135.9, 135.6, 132.4, 131.5, 129.5, 126.9, 123.5, 120.72, 120.65, 112.1, 102.2, 60.4, 54.3, 52.0, 50.0, 23.6; $\nu_{max}/cm^{-1}$ 2951, 2782, 1706 (C=O), 1615, 1505; HRMS (ESI)+: m/z calculated for $[C_{22}H_{24}N_2O_2 + H]^+$ = 349.1911, observed 349.1913.

**3-oxo-3-(1-(4-(pyrrolidin-1-ylmethyl)benzyl)-1H-indol-6-yl)propanenitrile**

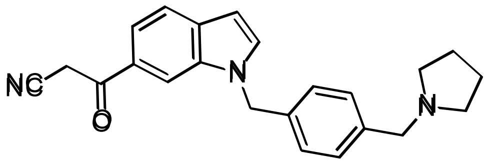


*n*-Butyllithium (1.6 M in hexanes, 2.7 mL, 4.3 mmol) was added dropwise at -78 °C to a mixture of acetonitrile (0.375 mL, 7.17 mmol) and THF (6 mL). The reaction mixture was stirred at -78 °C over 30 minutes. A solution of methyl 1-(4-(pyrrolidin-1-ylmethyl)benzyl)-1H-indole-6-carboxylate (0.500 g, 1.43 mmol) in THF (4 mL) was added dropwise at -78 °C over 30 minutes. The reaction mixture was stirred at -78 °C over 1 hour. Water (15 mL) was added dropwise at 0 °C, and the reaction mixture adjusted to pH 8. The product was extracted into DCM (3 x 25 mL). The combined organic extracts were washed (brine), dried ($MgSO_4$) and concentrated *in vacuo*. Purification by flash column chromatography (50 – 100% ethyl acetate in petroleum ether, 0 – 10% methanol in DCM) gave 3-oxo-3-(1-(4-(pyrrolidin-1-ylmethyl)benzyl)-1H-indol-6-yl)propanenitrile (0.448 g, 87% yield).

LCMS (ESI+): m/z 358.3 $[M + H]^+$, (ESI-): m/z 356.2 $[M - H]^-$, rt 1.38 minutes, 100%; $^1H$ NMR (400 MHz, $CDCl_3$) 7.98 (1H, s), 7.70 (1H, d, J = 8.3 Hz), 7.59 (1H, dd, J = 8.4, 1.3 Hz), 7.36 (1H, d, J = 3.1 Hz), 7.29 (2H, d, J = 8.1 Hz), 7.07 (2H, d, J = 7.9 Hz), 6.61 (1H, d, J = 3.1

Hz), 5.38 (2H, s), 4.09 (2H, br s), 3.59 (2H, s), 2.55-2.44 (4H, m), 1.83-1.71 (4H, m); $^{13}$C NMR (100 MHz, $CDCl_3$) 187.0, 139.4, 136.0, 135.2, 133.9, 133.4, 129.7, 128.1, 127.0, 121.3, 119.7, 114.5, 111.3, 102.6, 60.3, 54.3, 50.3, 29.6, 23.5; $\nu_{max}/cm^{-1}$ 2960, 2786, 2165 (C≡N), 1680 (C=O), 1607, 1503.

**3-amino-5-(1-(4-(pyrrolidin-1-ylmethyl)benzyl)-1H-indol-6-yl)-1H-pyrazole-4-carbonitrile (AW5)**

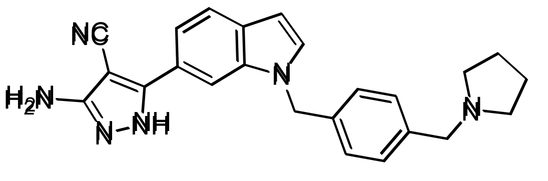


Trichloroacetonitrile (0.201 mL, 2.01 mmol) was added to a suspension of 3-oxo-3-(1-(4-(pyrrolidin-1-ylmethyl)benzyl)-1H-indol-6-yl)propanenitrile (0.239 g, 0.669 mmol) and sodium acetate (0.274 g, 3.34 mmol) in ethanol (5 mL). The reaction mixture was stirred over 4 hours, then $NaHCO_3$ solution (7.5 mL) and water (7.5 mL) were added. The intermediate was extracted into DCM (3 x 25 mL). The combined organic extracts were washed (brine), dried ($MgSO_4$) and concentrated *in vacuo*. The crude residue was dissolved in ethanol (4 mL) and hydrazine monohydrate (0.325 mL, 6.69 mmol) was added. The reaction mixture was heated under reflux for 18 hours, then quenched with excess acetone at room temperature and concentrated *in vacuo*. Purification by flash column chromatography (100% ethyl acetate, 0 – 7% methanol (+ 0.1% $NH_3$) in DCM) gave 3-amino-5-(1-(4-(pyrrolidin-1-ylmethyl)benzyl)-1H-indol-6-yl)-1H-pyrazole-4-carbonitrile (0.135 g, 51% yield).

LCMS (ESI+): m/z 397.3 $[M + H]^+$, (ESI-): m/z 395.3 $[M - H]^-$, rt 1.35 minutes, 100%; $^1$H NMR (500 MHz, $CDCl_3$) 7.70 (1H, s), 7.62 (1H, d, J = 8.3 Hz), 7.42 (1H, dd, J = 8.2, 1.5 Hz), 7.21 (2H, d, J = 8.2 Hz), 7.17 (1H, d, J = 3.0 Hz), 7.04 (2H, d, J = 8.1 Hz), 6.53 (1H, dd, J = 3.1, 0.8 Hz), 5.18 (2H, s), 4.40 (2H, br s), 3.55 (2H, s), 2.55-2.42 (4H, m), 1.75-1.64 (4H, m); $^{13}$C NMR (125 MHz, $CDCl_3$) 156.7, 152.0, 150.4, 138.5, 136.2, 136.0, 130.4, 130.1, 129.7, 127.2, 121.8, 117.9, 115.9, 108.0, 102.2, 75.4, 60.3, 54.2, 50.1, 23.4; $\nu_{max}/cm^{-1}$ 3100 (br, N-H),

2928, 2799, 2208 (C≡N), 1620, 1589, 1501; HRMS (ESI+): m/z calculated for [$C_{24}H_{24}N_6$ + H]$^+$ = 397.2135, observed 397.2132.

**5-(1-(4-(pyrrolidin-1-ylmethyl)benzyl)-1H-indol-6-yl)-1H-pyrazol-3-amine (AW6)**

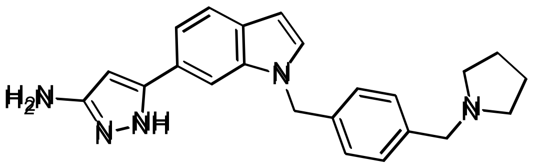

Hydrazine monohydrate (0.157 mL, 3.22 mmol) was added to a solution of 3-oxo-3-(1-(4-(pyrrolidin-1-ylmethyl)benzyl)-1H-indol-6-yl)propanenitrile (0.115 g, 0.322 mmol) in ethanol (15 mL). The reaction mixture was heated under reflux for 12 hours, then concentrated *in vacuo*. Purification by reverse phase column chromatography (0 – 100% acetonitrile in water, 0 – 65% methanol in acetonitrile) gave 5-(1-(4-(pyrrolidin-1-ylmethyl)benzyl)-1H-indol-6-yl)-1H-pyrazol-3-amine (27 mg, 22% yield).

LCMS (ESI+): m/z 372.5 [M + H]$^+$, rt 1.22 minutes, 96%; $^1$H NMR (500 MHz, $CDCl_3$) 7.63 (1H, d, J = 8.2 Hz), 7.43 (1H, s), 7.30-7.25 (1H, m), 7.24 (2H, d, J = 8.2 Hz), 7.13 (1H, d, J = 3.1 Hz), 7.03 (2H, d, J = 8.1 Hz), 6.53 (1H, dd, J = 3.1, 0.5 Hz), 5.87 (1H, s), 5.26 (2H, s), 3.68 (2H, br s), 3.55 (2H, s), 2.53-2.38 (4H, m), 1.81-1.67 (4H, m); $^{13}$C NMR (125 MHz, $CDCl_3$) 155.0 (br), 146.7 (br), 139.1, 136.6, 135.9, 129.6, 129.5, 129.0, 126.9, 124.0, 121.5, 117.7, 106.8, 102.0, 90.5, 60.4, 54.3, 50.0, 23.5; $\nu_{max}$/cm$^{-1}$ 3150 (br, N-H), 2963, 2789, 1587, 1505; HRMS (ESI+): m/z calculated for [$C_{23}H_{25}N_5$ + H]$^+$ = 372.2183, observed 372.2173.

**4-((4-methylpiperazin-1-yl)methyl)benzonitrile**

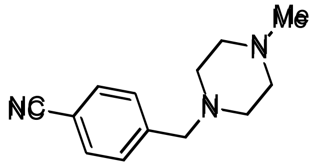

Sodium triacetoxyborohydride (2.59 g, 12.2 mmol) was added to a solution of 4-formylbenzonitrile (1.00 g, 7.63 mmol), 1-methylpiperazine (0.846 mL, 7.63 mmol) and acetic acid (0.524 mL, 9.15 mmol) in DCM (10 mL). The reaction mixture was stirred over

80 minutes. The reaction was diluted with water (10 mL), adjusted to pH 14 using aqueous NaOH (10% w/v) and extracted into DCM (3 x 25 mL). The combined organic extracts were dried ($MgSO_4$) and concentrated *in vacuo*. Purification by flash column chromatography (0 – 4% methanol in DCM) gave 4-((4-methylpiperazin-1-yl)methyl)benzonitrile (1.28 g, 70% yield).

$^1$H NMR (400 MHz, $CDCl_3$) 7.61-7.56 (2H, m), 7.44 (2H, d, J = 8.2 Hz), 3.53 (2H, s), 2.45 (8H, br s), 2.27 (3H, s); $^{13}$C NMR (100 MHz, $CDCl_3$) 144.4, 132.2, 129.6, 119.1, 111.0, 62.5, 55.2, 53.3, 46.1; spectroscopic data consistent with literature.[6]

**4-((4-methylpiperazin-1-yl)methyl)benzaldehyde**

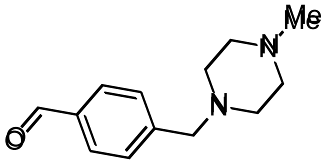


Diisobutylaluminium hydride (1 M in THF, 5.3 mL, 5.3 mmol) was added dropwise at 0 °C to a solution of 4-((4-methylpiperazin-1-yl)methyl)benzonitrile (1.26 g, 5.27 mmol) in THF (10 mL). The reaction mixture was warmed to room temperature and stirred over 30 minutes. Further diisobutylaluminium hydride (5.3 mL, 5.3 mmol) was added dropwise at 0 °C to the reaction mixture. The reaction mixture was warmed to room temperature and stirred over 30 minutes, then $NaHCO_3$ solution (50 mL) was added dropwise at 0 °C. The product was extracted into DCM/methanol (9:1, 3 x 50 mL). The combined organic extracts were washed (brine), dried ($MgSO_4$) and concentrated *in vacuo*. Purification by flash column chromatography (0 – 4% methanol (+ 0.1% $NH_3$) in DCM) gave 4-((4-methylpiperazin-1-yl)methyl)benzaldehyde (0.576 g, 50% yield).

$^1$H NMR (400 MHz, $CDCl_3$) 9.98 (1H, s), 7.85-7.79 (2H, m), 7.50 (2H, d, J = 8.0 Hz), 3.57 (2H, s), 2.47 (8H, br s), 2.28 (3H, s); $^{13}$C NMR (100 MHz, $CDCl_3$) 192.1, 146.0, 135.6, 129.9, 129.6, 62.8, 55.2, 53.3, 46.2; spectroscopic data consistent with literature.[7]

**methyl 1-(4-((4-methylpiperazin-1-yl)methyl)benzyl)-1H-indole-6-carboxylate**

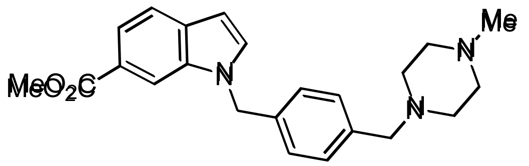

Toluene (3 mL) was added to a mixture of 4-((4-methylpiperazin-1-yl)methyl)benzaldehyde (0.300 g, 1.37 mmol), methyl indoline-6-carboxylate (0.314 g, 1.65 mmol) and benzoic acid (34 mg, 0.28 mmol). The reaction mixture was heated to 200 °C by microwave for 30 minutes. The reaction was diluted with ethyl acetate (25 mL), washed with $NaHCO_3$ solution (3 x 15 mL) and brine (15 mL), dried ($MgSO_4$) and concentrated *in vacuo*. Purification by flash column chromatography (0 – 10% methanol in DCM) gave methyl 1-(4-((4-methylpiperazin-1-yl)methyl)benzyl)-1H-indole-6-carboxylate (0.329 g, 55% yield).

LCMS (ESI+): m/z 378.3 $[M + H]^+$, rt 1.46 minutes, 100%; $^1$H NMR (400 MHz, $CDCl_3$) 8.12 (1H, s), 7.83 (1H, dd, J = 8.4, 1.4 Hz), 7.68 (1H, d, J = 8.4 Hz), 7.30-7.24 (3H, m), 7.08 (2H, d, J = 8.2 Hz), 6.61 (1H, dd, J = 3.1, 0.7 Hz), 5.38 (2H, s), 3.93 (3H, s), 3.49 (2H, s), 2.49 (8H, br s), 2.31 (3H, s); $^{13}$C NMR (100 MHz, $CDCl_3$) 168.1, 137.9, 135.8, 135.7, 132.3, 131.3, 129.6, 126.7, 123.4, 120.6, 120.5, 112.0, 102.1, 62.5, 55.0, 52.8, 51.9, 49.8, 45.8; $\nu_{max}$/cm$^{-1}$ 2937, 2794, 1706 (C=O), 1614, 1504; HRMS (ESI)+: m/z calculated for $[C_{23}H_{27}N_3O_2 + H]^+$ = 378.2176, observed 378.2159.

**3-(1-(4-((4-methylpiperazin-1-yl)methyl)benzyl)-1H-indol-6-yl)-3-oxopropanenitrile**

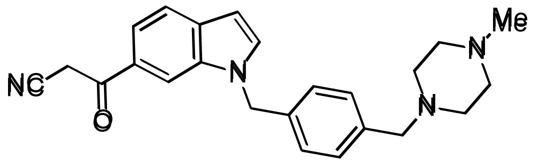

*n*-Butyllithium (1.6 M in hexanes, 1.4 mL, 2.2 mmol) was added dropwise at -78 °C to a mixture of acetonitrile (0.188 mL, 3.61 mmol) and THF (3 mL). The reaction mixture was stirred at -78 °C over 30 minutes. A solution of methyl 1-(4-((4-methylpiperazin-1-yl)methyl)benzyl)-1H-indole-6-carboxylate (0.313 g, 0.721 mmol) in THF (3 mL) was added dropwise at -78 °C over 30 minutes. The reaction mixture was stirred at -78 °C over 30 minutes.

Water (20 mL) was added dropwise at 0 °C, and the reaction mixture adjusted to pH 8. The product was extracted into DCM (3 x 25 mL). The combined organic extracts were washed (brine), dried ($MgSO_4$) and concentrated *in vacuo*. Purification by flash column chromatography (50 – 100% ethyl acetate in petroleum ether, 0 – 10% methanol in DCM) gave 3-(1-(4-((4-methylpiperazin-1-yl)methyl)benzyl)-1H-indol-6-yl)-3-oxopropanenitrile (0.258 g, 82% yield).

LCMS (ESI+): m/z 387.2 $[M + H]^+$, rt 1.38 minutes, 89%; $^1$H NMR (400 MHz, $CDCl_3$) 7.98 (1H, s), 7.69 (1H, d, J = 8.3 Hz), 7.58 (1H, dd, J = 8.4, 1.6 Hz), 7.36 (1H, d, J = 3.2 Hz), 7.26 (2H, d, J = 8.1 Hz), 7.06 (2H, d, J = 8.2 Hz), 6.61 (1H, dd, J = 3.1, 0.7 Hz), 5.37 (2H, s), 4.09 (2H, br s), 3.47 (2H, s), 2.44 (8H, br s), 2.27 (3H, s); $^{13}$C NMR (100 MHz, $CDCl_3$) 187.0, 138.4, 136.0, 135.3, 133.8, 133.4, 129.9, 128.1, 126.9, 121.3, 119.7, 114.5, 111.3, 102.6, 62.6, 55.2, 53.1, 50.3, 46.1, 29.7; $\nu_{max}/cm^{-1}$ 2934, 2790, 2165 (C≡N), 1680 (C=O), 1607, 1502; HRMS (ESI)-: m/z calculated for $[C_{24}H_{26}N_4O - H]^-$ = 385.2034, observed 385.2029.

**3-amino-5-(1-(4-((4-methylpiperazin-1-yl)methyl)benzyl)-1H-indol-6-yl)-1H-pyrazole-4-carbonitrile (AW7)**

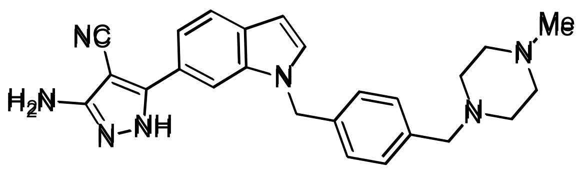


Trichloroacetonitrile (0.167 mL, 1.66 mmol) was added to a suspension of 3-(1-(4-((4-methylpiperazin-1-yl)methyl)benzyl)-1H-indol-6-yl)-3-oxopropanenitrile (0.241 g, 0.555 mmol) and sodium acetate (0.228 g, 2.77 mmol) in ethanol (5 mL). The reaction mixture was stirred over 17 hours, then $NaHCO_3$ solution (25 mL) was added. The intermediate was extracted into DCM (3 x 25 mL). The combined organic extracts were washed (brine), dried ($MgSO_4$) and concentrated *in vacuo*. The crude residue was dissolved in ethanol (10 mL) and hydrazine monohydrate (0.270 mL, 5.55 mmol) was added. The reaction mixture was heated under reflux for 24 hours, then concentrated *in vacuo*. Purification by reverse phase column chromatography (0 – 50% acetonitrile in water (+ 0.1% formic acid)) was attempted, with the resultant fractions combined, adjusted to pH 8, and extracted into DCM/methanol (9:1, 6 x 50 mL). The combined organic extracts were dried ($MgSO_4$) and concentrated *in vacuo*. Purification by flash column chromatography (0 – 20% methanol (+ 0.1% $NH_3$) in DCM) gave 3-amino-5-(1-(4-((4-methylpiperazin-1-yl)methyl)benzyl)-1H-indol-6-yl)-1H-pyrazole-4-carbonitrile (0.140 g, 59% yield).

LCMS (ESI+): m/z 426.4 $[M + H]^+$, (ESI-): m/z 424.3 $[M - H]^-$, rt 1.36 minutes, 100%; $^1H$ NMR (500 MHz, $CD_3OD$) 7.84-7.81 (1H, m), 7.64 (1H, d, J = 8.3 Hz), 7.49 (1H, dd, J = 8.2, 1.4 Hz), 7.39 (1H, d, J = 3.1 Hz), 7.27-7.21 (2H, m), 7.21-7.16 (2H, m), 6.53 (1H, dd, J = 3.1, 0.7 Hz), 5.37 (2H, s), 3.46 (2H, s), 2.44 (8H, br s), 2.24 (3H, s); $^{13}C$ NMR (125 MHz, $CD_3OD$) 158.0 (br), 152.5 (br), 138.3, 137.7, 137.4, 131.6, 131.4, 131.1, 128.2, 124.0 (br), 122.2, 118.7, 117.0, 109.1, 102.7, 74.0 (br), 63.3, 55.5, 53.3, 50.8, 45.8; $\nu_{max}/cm^{-1}$ 3150 (br, N-H), 2919, 2808, 2207 (C≡N), 1612, 1581; HRMS (ESI+): m/z calculated for $[C_{25}H_{27}N_7 + H]^+$ = 426.2401, observed 426.2396.
